# GraphPop: graph-native computation decouples population genomics complexity from sample count

**DOI:** 10.64898/2026.04.11.717929

**Authors:** Ehsan Estaji, Shi-Wei Zhao, Zhao-Yang Chen, Shuai Nie, Jian-Feng Mao

## Abstract

Matrix-based population genomics tools scale as ***O*(*V* × *N* )**, re-reading the full genotype matrix for every analysis. Here we present GraphPop, a graph database engine that reduces summary statistic complexity to ***O*(*V* × *K*)** where ***K*** is population count—independent of sample count—by computing on pre-aggregated allele-count arrays stored as graph node properties. The same architecture enables annotation-conditioned queries via edge traversal, persistent analytical records, and multi-statistic composition. Applied to rice 3K (29.6M SNPs, 3,024 accessions) and human 1000 Genomes (3,202 samples, 22 autosomes), GraphPop reveals that all 12 rice subpopulations show ***π_N_ /π_S_ >* 1.0**, uncovers opposite consequence-level Fst regimes between species, and identifies *KCNE1* as a candidate pre-Out-of-Africa sweep via convergence of five stored statistics. GraphPop achieves 146–327**×** query-time speedup for pre-aggregated statistics and 63–179**×** for bit-packed haplotype computation (iHS, XP-EHH, nSL), at constant **∼**160 MB procedure working memory. This complexity reduction makes systematic, annotation-integrated population genomics practical at scale.

## Introduction

The statistical toolkit of population genomics—nucleotide diversity, Fst, haplotype homozygosity, runs of homozygosity—has been refined over decades. Yet as datasets grow from hundreds to tens of thousands of individuals, with whole-genome variant counts reaching tens of millions, four challenges have become acute.

First, *computational complexity scales linearly with sample count*. Matrix-based tools (scikit-allel[1], VCFtools[2], PLINK[3, 4]) scale as *O*(*V* × *N* ) where *V* is variant count and *N* is sample count, re-reading the entire genotype matrix for every analysis. Doubling sample count doubles computation time, even for statistics that depend only on allele frequencies. For a crop programme tracking 12 populations across 12 chromosomes, computing diversity alone requires 144 full-matrix reads—redundant computation, because per-population allele frequencies could be computed once during import and reused indefinitely. No existing tool separates import-time aggregation from query-time computation.

Second, *annotation conditioning is disconnected from statistical computation*. The cost of domestication (*π_N_ /π_S_*) has been estimated through interspecific comparisons[5], but systematic quantification *within* cultivated rice—across all sub-populations, at genome scale—has not been performed. Computing *π_N_ /π_S_* requires separate diversity calculations for missense and synonymous variants, each demanding a multi-tool pipeline: VEP annotation[6], consequence filtering, VCF subsetting, and a separate diversity computation. For 12 populations × 12 chromosomes × 2 consequence classes, this means 288 pipeline runs. bedtools[7] can intersect coordinates, but no tool computes a statistic restricted to a given functional class without a multi-step pipeline.

Third, *multi-statistic composition requires manual file coordination*. Identifying convergent positive selection across continental lineages requires cross-querying iHS[8], XP-EHH[9], Garud’s *H*_12_[10], and Fst[11] on the same genomic variants. Classical tools compute these statistics into separate output files; intersecting them at genome scale across 22 autosomes and 26 populations is feasible for a single locus but impractical as a systematic survey. No persistent analytical record links the results of independently computed statistics to the variants and genes that produced them.

Fourth, *second-order queries on stored results are not supported*. Do functionally related pathways show coordinated divergence across population pairs? Answering this requires correlating pathway-level Fst values across dozens of pairwise comparisons— an intermediate representation that no classical tool produces, because results are written to isolated files rather than stored as queryable properties of the data objects that generated them.

These four challenges share a common root: the flat-file, matrix-based paradigm in which every analysis starts from raw genotypes, computed results are ephemeral, functional annotations live in separate files, and computational cost is paid in full for every query.

### Graph databases as an alternative data model

Graph databases store data as nodes and edges with key-value properties[12, 13]. Two features are relevant here. First, the *labelled property graph* model[14] attaches quantitative properties (allele frequencies, Fst values) directly to typed nodes (Variant, Gene, Pathway) and edges (HAS_CONSEQUENCE, IN_PATHWAY). Second, *index-free adjacency*[12] means traversing a relationship costs *O*(1) per hop regardless of database size, compared to *O*(log *N* ) index lookups in relational systems. Together, these properties make pre-aggregated population statistics immediately composable with functional annotations via edge traversal, without join operations or coordinate-based file intersection. Graph databases have been adopted for biological pathway databases (e.g., Reactome[15]) and knowledge graphs[16]—domains where entity relationships carry as much information as the entities themselves.

### GraphPop

Here we present GraphPop, an analytical engine comprising 12 stored procedures inside a graph database. The core design separates *import-time aggregation* from *query-time computation*: per-population allele counts (AC, AN, AF) are pre-aggregated during a one-time import and stored as arrays on Variant nodes. FAST PATH statistics (diversity, Fst, SFS, and others) then operate on these arrays with *O*(*V* × *K*) complexity, where *K* is the number of populations—independent of sample count. A dataset of 3,000 samples computes in the same time as one of 300,000, because individual genotypes are never re-read after import. For FULL PATH haplotype statistics (iHS, XP-EHH, nSL, ROH), GraphPop uses bit-packed haplotype matrices (1 bit per haplotype, 87% memory reduction) with SIMD-accelerated kernels. The complexity reduction is not specific to the current implementation: it follows from the graph model’s ability to store pre-computed aggregates as node properties, and would hold on any graph database engine.

The same architecture provides annotation-conditioned queries resolved by edge traversal, a persistent analytical record in which computed statistics become permanent node properties, and second-order queries on stored results without recomputation.

Benchmarking against scikit-allel[1], VCFtools[2], PLINK 2[4], and bcftools[17] on 1000 Genomes chr22 confirms numerical accuracy (*<* 0.000001% relative error for summary statistics; *r* = 0.96–0.999 for haplotype statistics). After a one-time import, FAST PATH query-time speedups reach 146–327× over scikit-allel; FULL PATH achieves 63–179× for iHS, XP-EHH, and nSL through bit-packed computation (Table 1). Both paths maintain constant ∼160 MB procedure working memory.

**Table 1.** Query-time benchmark on 1000 Genomes chr22. (3,202 samples, 1,066,555 CEU-polymorphic variants out of 1,070,401 total). GraphPop times are query-time after one-time database import (∼12 min for chr22; ∼4.5 h for full genome). FAST PATH speedups reflect pre-aggregated allele counts; FULL PATH speedups reflect bit-packed computation and SIMD optimisation. Competitor tools read from their native formats (VCF or PLINK binary). Procedure working memory: GraphPop ∼160 MB constant; scikit-allel 1,051 MB; PLINK 2 1,665 MB; bcftools 1,759 MB.

| Statistic | GraphPop (s) | Competitor (s) | Tool | Speedup | Accuracy |
| --- | --- | --- | --- | --- | --- |
| *FAST PATH (pre-aggregated allele counts)* |  |  |  |  |  |
| $\pi/\theta_W/\text{Tajima's } D$ | 12.1 | 1,757 | scikit-allele | $146\times$ | $< 0.000001\%$ |
| $F_{st}/D_{xy}$ | 8.8 | 2,165 | scikit-allele | $245\times$ | $< 0.000001\%$ |
| SFS | 5.9 | 1,937 | scikit-allele | $327\times$ | $< 0.000001\%$ |
| *FULL PATH (bit-packed haplotype matrices)* |  |  |  |  |  |
| iHS | 10.9 | 1,948 | scikit-allele | $179\times$ | $r = 0.999$ |
| XP-EHH | 16.7 | 2,280 | scikit-allele | $136\times$ | $r = 0.97$ |
| nSL<sup>†</sup> | 35 | 2,231 | scikit-allele | $63\times$ | $r = 0.9997$ |
| ROH<sup>†</sup> | 10.9 | 50 | bcftools | $4.6\times$ | $r = 0.96$ |
| LD ($r^2$) | 6.4 | 1.1 | PLINK 2 | $0.17\times$ | $< 0.1\%$ |

<sup>†</sup>GraphPop-numpy path. Graph database times: nSL 307 s, ROH 128 s. PLINK 2 faster for $F_{st}$ alone (3.9 s) but does not compute $D_{xy}$ .

GraphPop is designed for datasets typical in agricultural, ecological, and evolutionary population genomics—hundreds to tens of thousands of individuals with millions to tens of millions of variants—where it delivers the full *O*(*V* × *K*) advantage (Supplementary Note S9). We validate on the 1000 Genomes Project[18] (3,202 samples, 22 autosomes) and the 3K Rice Genomes Project[19] (3,024 accessions, 29.6M SNPs, 12 subpopulations). The analyses demonstrate results that would be operationally prohibitive with file-based tools: a systematic survey of purifying selection relaxation across all cultivated rice subpopulations (*π_N_ /π_S_>* 1.0), a crossspecies reversal in consequence-level selection regimes, a candidate pan-continental sweep at *KCNE1* detectable by multi-statistic convergence, and coordinated pathway co-selection networks in both species.

### GraphPop architecture and dual computational paths

GraphPop organises population genomic data as a labelled property graph (Fig. 1b). **Variant** nodes store genomic coordinates, alleles, per-population allele-count arrays (ac[], an[], af[]) indexed by a shared pop ids[] key, and bit-packed individual genotype arrays (gt packed at 2 bits/sample, phase packed at 1 bit/sample). Functional annotation edges connect Variants to **Gene** nodes via HAS_CONSEQUENCE (carrying VEP-derived consequence type and impact level), and Genes connect to **Pathway** and **GOTerm** nodes via IN_PATHWAY and HAS_GO_TERM edges. **GenomicWindow** nodes, materialised by genome scan procedures, persist per-window statistics as permanent queryable records. The path from a variant’s genotype to its functional annotation is a direct edge traversal, exploiting index-free adjacency to resolve annotation conditioning in constant time per hop.

**Fig. 1.**
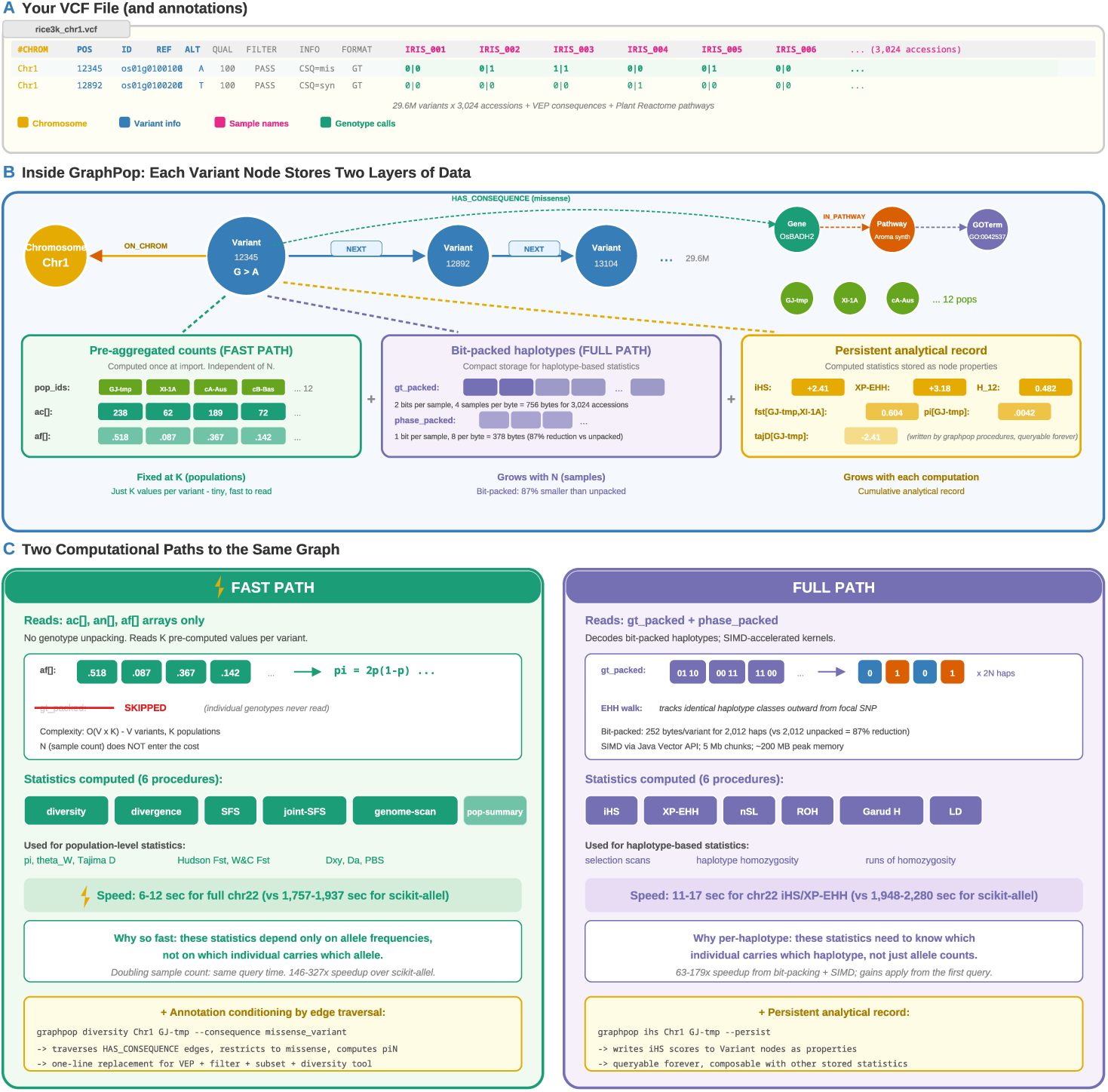
GraphPop architecture: from VCF to persistent analytical graph. **a**, Input: a standard VCF file with chromosome, position, reference/alternate alleles, sample columns, and pervariant genotype calls (rice 3K dataset shown; 29.6M variants × 3,024 accessions). VEP consequence annotations and Plant Reactome pathways are loaded alongside. **b**, Inside the graph database, each Variant node stores three layers of data: *(left)* pre-aggregated population allele counts (ac[], an[], af[]) indexed by population—fixed at *K* values per variant, the input to FAST PATH; *(middle)* bit-packed individual genotypes (gt_packed at 2 bits/sample, phase_packed at 1 bit/sample), 87% smaller than the unpacked equivalent, the input to FULL PATH; *(right)* the persistent analytical record—computed statistics (iHS, XP-EHH, *H*_12_, *π*, Fst, Tajima’s *D*) written to the same Variant nodes as queryable properties. Variants are linked by NEXT edges and to Gene, Pathway, and GOTerm nodes via HAS_CONSEQUENCE, IN_PATHWAY, and HAS_GO_TERM edges. **c**, Two computational paths to the same graph. FAST PATH (left) reads only the pre-aggregated af[]/ac[] arrays—*O*(*V* × *K*) complexity, independent of sample count, 6–12 s for full chr22, supporting diversity, divergence, SFS, joint SFS, genome scan, and pop summary procedures. FULL PATH (right) decodes bit-packed haplotypes using SIMD-accelerated kernels and chunked processing—11–17 s for chr22 iHS/XP-EHH, supporting iHS, XP-EHH, nSL, ROH, Garud’s *H*, and LD procedures. Each path also enables a key architectural capability (yellow callouts): annotation conditioning via edge traversal (FAST PATH) and a persistent analytical record via the --persist flag (FULL PATH).

### Dual computational paths

GraphPop implements two complementary strategies (Fig. 1c). **FAST PATH** procedures (diversity, divergence, SFS, joint SFS, genome scan, population summary) operate exclusively on pre-aggregated allele-count arrays stored as node properties. Because counts are computed once during import, runtime complexity is *O*(*V* × *K*) where *V* is variant count and *K* is population count—independent of sample count. For 1000 Genomes chr22 (1,070,401 variants), FAST PATH procedures complete in 1–12 seconds. Matrix-based tools such as scikit-allel scale as *O*(*V* × *N* ) because each analysis decompresses *N* genotypes per variant. A step-by-step worked example contrasting matrix-based and FAST PATH computation of *π* and Fst is shown in Supplementary Figs. S8 and S9.

**FULL PATH** procedures (LD, iHS, XP-EHH, nSL, ROH, Garud’s *H*) require individual haplotype data. GraphPop reads gt packed and phase_packed arrays into a dense HaplotypeMatrix where each haplotype occupies 1 bit (packed 8 per byte). For 1,006 EUR samples (2,012 haplotypes), each variant row requires 252 bytes versus 2,012 bytes unpacked—an 87% reduction. A chunking strategy processes chromosomes in 5 Mb core windows with 2 Mb EHH margins, keeping peak memory under 200 MB regardless of chromosome length. Numerical kernels use Java’s Vector API (JEP 338) for SIMD acceleration. Step-by-step worked examples of EHH/iHS computation in conventional and FULL PATH form are shown in Supplementary Figs. S10 and S11.

### Annotation-conditioned queries

Every procedure accepts consequence, pathway, or gene filters. Internally, VariantQuery constructs a graph traversal through HAS_CONSEQUENCE or IN_PATHWAY edges before returning variants to the procedure (Fig. 1c). The procedure code is identical whether or not conditioning is applied—only the traversal pattern changes. Computing *π_N_* for temperate japonica rice on chromosome 1:

CALL graphpop.diversity(’Chr1’, 1, 43270923, ’GJ-tmp’,

{consequence: ’missense_variant’})

The classical equivalent requires VEP annotation, consequence filtering, VCF subsetting, and a separate diversity tool—four invocations and three intermediate files. For 12 populations × 12 chromosomes × 2 consequence classes, GraphPop replaces 288 multi-step pipeline runs with 288 single-line procedure calls.

### Persistent analytical record

Computed results become part of the graph. Genome scan procedures write GenomicWindow nodes storing per-window *π*, *θ_W_* , Tajima’s *D*, Fst, PBS, and Fay & Wu’s *H*. Selection scan procedures write iHS, XP-EHH, and nSL scores as properties on Variant nodes. Population summaries are stored on Population nodes. In each case, the analytical record is stored as properties on the same nodes that represent the biological data, not in separate result tables.

Once computed, results are permanently available for cross-query without recomputation. A graph query can retrieve all variants with |iHS| *>* 2, all windows where Fst *>* 0.5 and *π <* 0.01, or intersect selection signals with gene and pathway annotations via edge traversal. This cumulative record differs from the ephemeral outputs of file-based tools, where each result exists as an isolated file that must be loaded, parsed, and coordinate-matched to integrate with other analyses.

The practical consequence is that second-order analyses—computations on the results of prior computations—require no additional data loading or file coordination (Supplementary Fig. S7). The pathway co-selection analysis below correlates pathwaylevel Fst values stored across 66 population pairs; in a classical workflow, these values would be scattered across hundreds of output files. Hail[20], the most comparable platform, stores results in separate matrix tables; composing results across analyses requires explicit table joins. In GraphPop, co-location on the same graph nodes makes cross-query a traversal rather than a join.

### Software architecture

GraphPop comprises a graph database compute engine (12 stored procedures in Java), an import pipeline (Python), a 62-command CLI organised into 11 functional domains (Supplementary Fig. S1; Supplementary Note S8), and a programmatic interface via the Model Context Protocol (MCP, 21 tools). Users require no knowledge of graph query languages—the graph architecture is fully abstracted behind a conventional command-line interface.

### Multi-statistic convergence identifies a candidate pre-Out-of-Africa sweep at *KCNE1*

Identifying convergent positive selection—genes under sweep in multiple independent populations—classically requires computing selection statistics per population, then intersecting results by genomic coordinate across output files. For 26 populations, 22 autosomes, and 3 statistics, this means managing ∼1,700 output files and writing custom scripts to find overlapping signals.

In GraphPop, all selection statistics are stored as properties on Variant and GenomicWindow nodes. Convergent selection reduces to a graph pattern match: query for nodes where stored |iHS| *>* threshold *and* |XP-EHH| *>* threshold, then traverse to Gene and Pathway annotations—one query, no re-computation.

We applied all 12 procedures to the 1000 Genomes Phase 3 dataset[18] (3,202 samples, 26 sub-populations, 22 autosomes, ∼70.7M Variant nodes, 91,973 Gene nodes, 2,241 Reactome pathways[15]). Querying for genes with Garud *H*_12_ *>* 0.3 in ≥2 continental groups identified **9 genes** (Fig. 2). ***KCNE1*** was the only gene with sweep evidence in all five groups (*H*_12_ = 1.0 in multiple populations; between-group Fst ratio = 0.296). Elevated between-group Fst combined with universally high *H*_12_ is consistent with an ancient sweep that reached near-fixation before the Out-of-Africa bottleneck[9]. *KCNE1* was previously identified in single-population iHS scans[8]; our contribution is detecting the five-group convergence pattern, which requires colocated multi-statistic queries. We note that the independence assumption underlying the *p <* 10*^−^*^6^ estimate (Methods) is approximate, as non-African populations share demographic history; however, the signal includes AFR, which does not share the Outof-Africa bottleneck. Formal calibration under demographic simulation is warranted for confirmatory analysis.

**Fig. 2.**
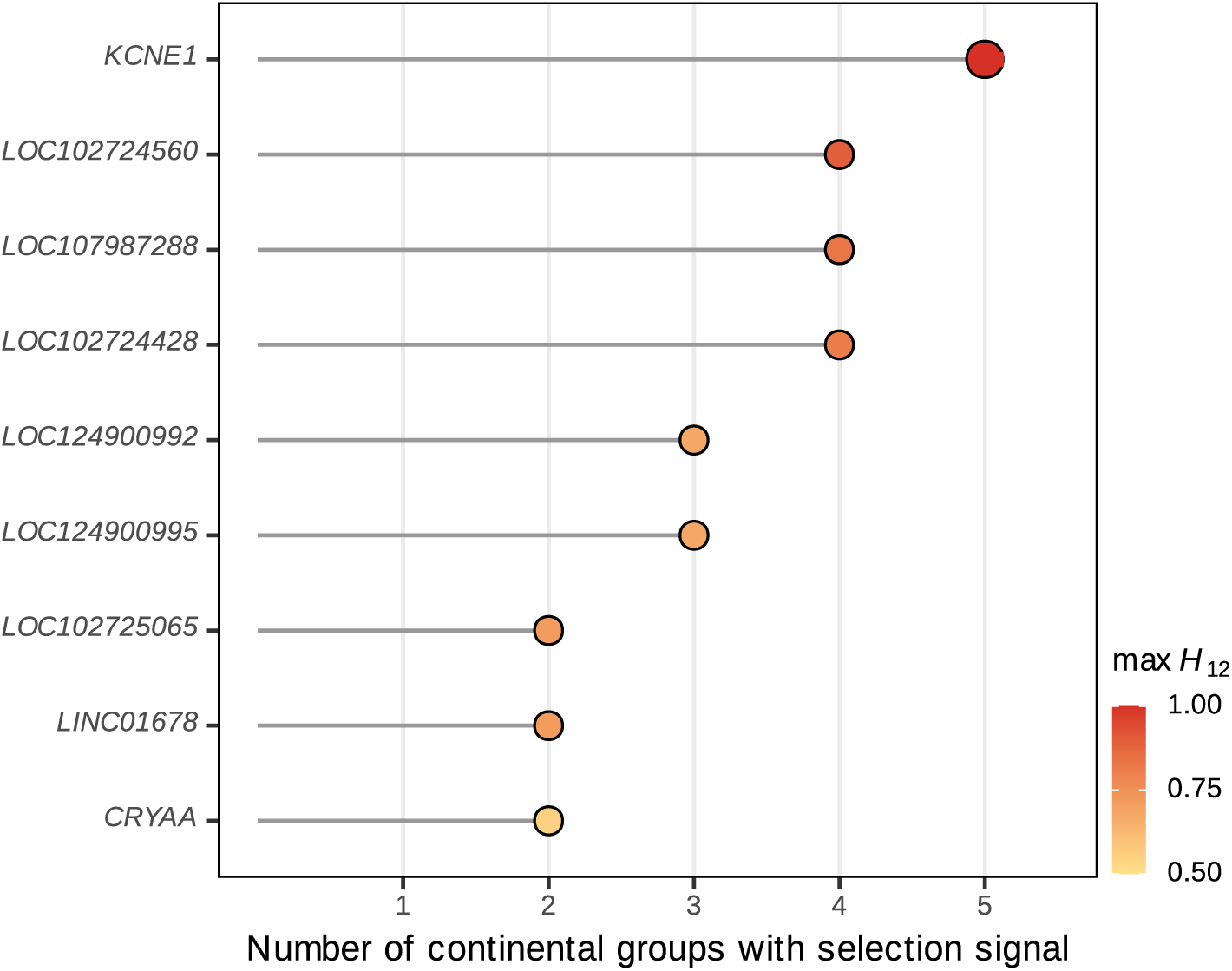
Multi-statistic convergence reveals convergent positive selection. Nine genes show Garud *H*_12_ *>* 0.3 in ≥2 independent continental groups, detected by querying stored statistics on Variant nodes. *KCNE1* is the only gene with sweep evidence in all five groups, consistent with an ancient pre-Out-of-Africa sweep.

*KCNE1* encodes the regulatory subunit of the slow delayed rectifier potassium current (*I_Ks_*), which together with the *K_V_* 7.1 pore-forming subunit (*KCNQ1* ) governs cardiac action potential repolarisation; loss-of-function variants cause type-1 long QT syndrome. Graph traversal through HAS_CONSEQUENCE → Gene → IN_PATHWAY placed *KCNE1* in the Phase 3 rapid repolarisation pathway alongside *KCNE4*, *KCNH2* (*K_V_* 11.1, the human ether-à-go-go-related gene), and *KCNQ1* —a functionally coherent cluster of cardiac ion-channel subunits, all population-differentiated, all in the same Reactome pathway (Supplementary Fig. S2). The convergent signal across all five continental groups suggests that the selective pressure predates the Out-of-Africa migration (∼60–70 kya); plausible drivers include thermoregulation in ancestral African environments[8] or pathogen-driven selection on cardiac function, though the specific selective agent remains an open question.

This finding required five independently computed statistics co-located on the same graph nodes. No single tool produces iHS, XP-EHH, *H*_12_, Fst, and pathway membership together; the graph is the structure that makes their intersection a query rather than a pipeline.

### Annotation conditioning quantifies the cost of domestication and reveals opposite selection regimes

The cost of domestication—the accumulation of mildly deleterious mutations due to demographic bottlenecks and relaxed selection[5]—has traditionally been quantified via interspecific *π_N_ /π_S_* comparisons (cultivated vs. wild ancestor). Here we use the term to describe the *intraspecific* mutational load signature: the elevation of *π_N_ /π_S_*above neutral expectation within cultivated subpopulations. Quantitative evidence for this intraspecific pattern across all subpopulations has been lacking, because computing *π_N_ /π_S_* at genome scale requires annotation-conditioned diversity calculations that no existing tool provides as a single operation.

By calling graphpop.diversity with consequence conditioning for all 12 populations × 12 chromosomes (288 calls, each replacing a 4-step pipeline), we found that **all 12 subpopulations show** *π_N_ /π_S_ >* 1.0 (range: 1.018–1.146; Fig. 3a, Supplementary Table S3). This universal elevation is consistent with the nearly-neutral theory[21]: domestication bottlenecks reduce effective population size, allowing mildly deleterious mutations that would be purged under stronger drift-selection balance to persist. The genome-wide pattern across all 12 subpopulations—including both severely bottlenecked (GJ-tmp, *π* = 0.019) and moderately bottlenecked (XI-1A, *π* = 0.041) groups—favours relaxed purifying selection over locus-specific positive selection, which would elevate *π_N_ /π_S_* at selected loci rather than genome-wide[22].

**Fig. 3.**
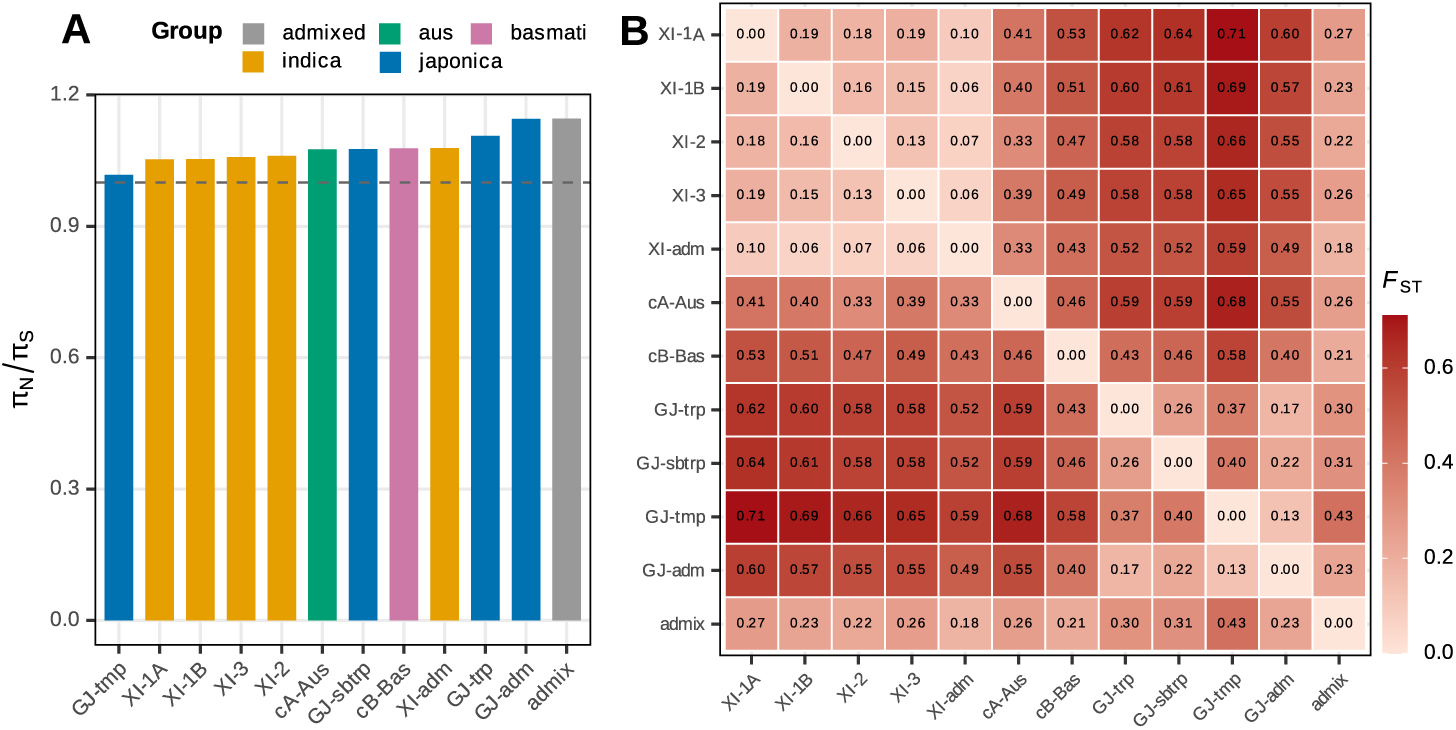
Annotation-conditioned cost of domestication. **a**, *π_N_ /π_S_* for all 12 rice subpopulations, computed via annotation conditioning. All values exceed 1.0 (dashed line), demonstrating universal relaxation of purifying selection. Human populations (not shown) all fall below 1.0 (range: 0.646– 0.673). **b**, Pairwise W&C Fst landscape across 66 population pairs, recovering the indica–japonica split (max Fst = 0.71).

The population-level variation in *π_N_ /π_S_* is itself informative. Admixed groups show the highest ratios (admix: 1.146, GJ-adm: 1.145), consistent with the expectation that hybridisation between bottlenecked source populations concentrates slightly deleterious alleles inherited from multiple founders. GJ-tmp (temperate japonica) shows the lowest ratio (1.018) despite having the strongest demographic bottleneck (*π* = 0.019, FROH = 0.080, Tajima’s *D* = −2.41). This apparent contradiction resolves if intense directional selection during temperate adaptation has purged deleterious missense variants from the very genes under selection, partially counteracting the genomewide relaxation. Indica subpopulations cluster narrowly (1.053–1.079), consistent with their larger effective population sizes sustaining more efficient purifying selection. This finding is robust to allele frequency filtering, call rate thresholds, per-chromosome decomposition, and annotation definitions (Supplementary Note S7).

The same analysis applied to all 26 human populations yields *π_N_ /π_S_* of 0.646–0.673 (all well below 1.0), confirming efficient purifying selection in natural populations. African populations show the lowest ratios (0.646–0.651), consistent with the largest long-term *N_e_*, while out-of-Africa populations show modestly elevated values (0.668– 0.673, ∼3.5% higher)—a subtle bottleneck signal that is dwarfed by the wholesale reversal in domesticated rice.

The biological implication of universal *π_N_ /π_S_ >* 1.0 in rice is that every cultivated subpopulation carries an elevated load of mildly deleterious missense variants relative to synonymous expectations. For crop improvement, this has practical consequences. Introgression of beneficial alleles from one subpopulation to another—a routine breeding strategy—risks transferring linked deleterious variants in proportion to the donor’s load, with admixed groups (the highest-load reservoirs in our survey) being particularly risky donors. Conversely, the observation that GJ-tmp shows the lowest genome-wide ratio despite the strongest bottleneck suggests that genomic regions under intense directional selection during temperate adaptation have been partially purged of deleterious variants, an effect that is invisible to species-wide *π_N_ /π_S_*averages but emerges clearly when annotation conditioning is applied per subpopulation. These findings reframe the cost of domestication from a single number per species into a structured signature that varies with demographic and selective history.

The cross-species contrast extends to population differentiation. Consequence-conditioned Fst reveals opposite selection regimes: in humans (YRI vs CEU), HIGH-impact variants show lower mean Fst (0.124) than LOW-impact variants (0.143; Mann–Whitney *p* ≈ 0)—purifying selection constraining differentiation at functional sites. In rice (GJ-tmp vs XI-1A), the pattern reverses: HIGH-impact Fst (= 0.185) exceeds MODERATE (= 0.129) and LOW (= 0.092; *p <* 10*^−^*^150^), indicating that directional selection during domestication has driven differentiation at the very sites where natural selection imposes constraint. Missense Fst exceeds synonymous Fst in all three major population pairs (Fig. 4), with the strongest signal at the temperate– tropical japonica boundary (ratio 1.106). Each of these comparisons used the same GraphPop command applied to two datasets, avoiding separate annotation-filtering pipelines per species (Supplementary Fig. S6).

**Fig. 4.**
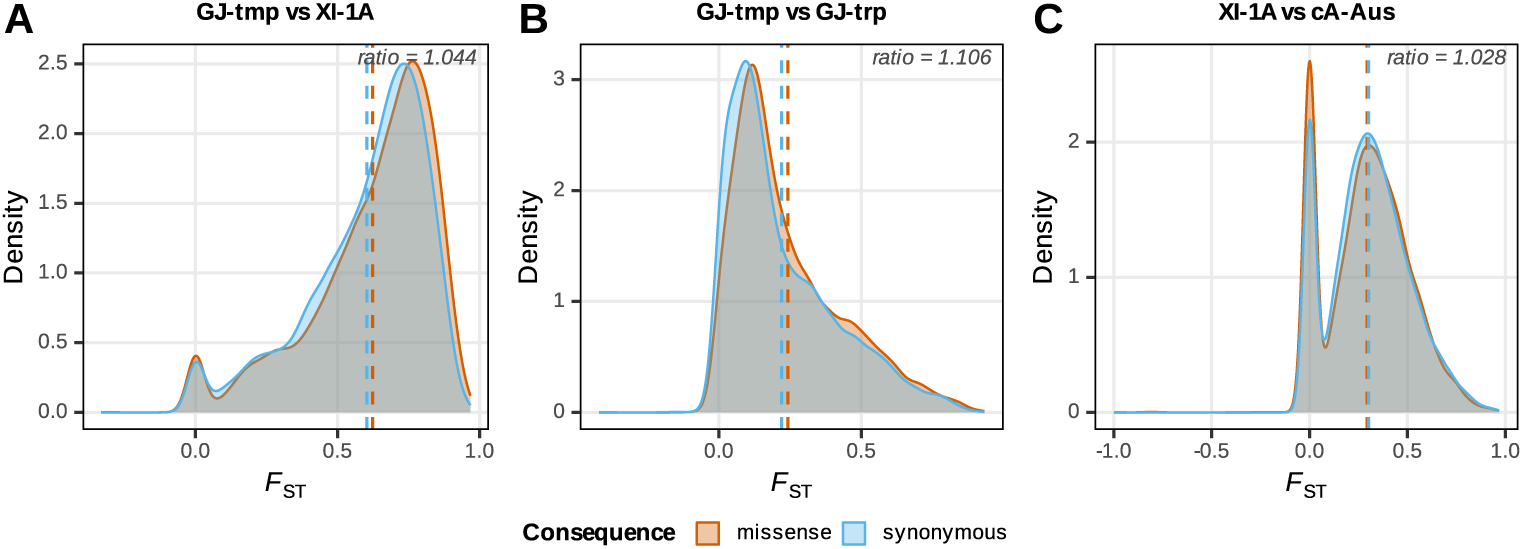
Adaptive protein divergence via conditioned genome scan. Sliding-window Fst distributions for missense versus synonymous variants across three major population pairs. The missense:synonymous ratio exceeds 1.0 in all comparisons, indicating directional selection on protein-coding variants during ecological specialization.

Standard statistics validated the graph database: pairwise W&C Fst recovered the indica–japonica split (Fst = 0.71; Fig. 3b), and XP-EHH and Garud’s *H* recovered known domestication genes (*GW5*, *Hd1*, *PROG1*, *Wx*, *OsC1*, *DRO1* [23, 24]) at expected genomic positions (Supplementary Table S6).

### Pathway co-selection reveals coordinated functional divergence

The persistent analytical record enables second-order queries: computations on the results of prior computations. By persisting pathway-level Fst as node properties across all pairwise comparisons, GraphPop can correlate pathway differentiation profiles—not “which pathways are differentiated?” but “which co-differentiate in concert?”

To identify coordinated selection, pathway-level Fst was computed for every population pair by averaging per-variant Fst across all variants assigned to each pathway via graph traversal (Variant → Gene → Pathway). Pairwise Spearman correlations between pathways across the population-pair vector were then computed, and pairs with *ρ >* 0.7 (Bonferroni-corrected *p <* 4.5 × 10*^−^*^6^ for ∼11,000 pathway pairs) were retained as edges in a co-selection network (see Methods). In rice, this analysis across all 66 population pairs for 153 Plant Reactome pathways[25] identified three co-selection modules: secondary metabolism (37 pathways, including coumarin, gibberellin, and cholesterol biosynthesis), amino acid/hormone biosynthesis (29 pathways, including folate, choline, and ethylene biosynthesis), and core cellular machinery (87 pathways, including DNA replication, translation, and vesicle transport). The housekeeping module showed the highest internal connectivity (hub degree 86*/*153), consistent with a genomic-background mode of differentiation. The two smaller modules capture ecotype-specific divergence: secondary metabolism is the chemical interface between rice and its environment (defence compounds, pigments, signalling molecules), and the strong co-differentiation of these pathways is consistent with parallel selection during the adaptation of indica and japonica subpopulations to distinct ecological niches; hormone biosynthesis pathways differentiate in concert because plant hormone signalling is a tightly coupled network where selection on one component creates pressure on related components.

In human, the same analysis across 2,232 Reactome pathways and 5 continental pairs revealed an analogous three-community structure: DNA repair and apoptosis pathways, immune and neuronal signalling pathways, and metabolic processes (Supplementary Notes S2, S4). The convergence in architecture—housekeeping pathways forming the largest, most connected module in both species—suggests that coordinated functional divergence is a general feature of population differentiation, independent of whether it is driven by natural demography (human) or artificial selection (rice). This analysis required pathway Fst stored as node properties across dozens of population pairs—an intermediate representation that file-based tools do not produce.

### Validation and performance

GraphPop was validated against scikit-allel[1], VCFtools[2], PLINK 2[4], PLINK 1.9[3], and bcftools[17] on 1000 Genomes chr22 (Table 1). FAST PATH statistics matched scikit-allel to *<* 0.000001% relative error—numerically equivalent within floating-point precision. FULL PATH correlations were: iHS *r* = 0.999 vs scikit-allel, XP-EHH *r* = 0.97, nSL *r* = 0.9997, ROH *r* = 0.96 vs bcftools. The non-perfect XP-EHH and ROH correlations reflect differences in tunable parameters (EHH truncation thresholds, ROH HMM transition rates) and edge handling at chromosome boundaries, not algorithmic discrepancies; per-locus comparisons confirmed identical detection of all top selection signals (Supplementary Fig. S12; Supplementary Note S6).

The two paths derive their performance from distinct sources. FAST PATH speedups (146–327×) reflect the *O*(*V* × *K*) complexity reduction via pre-aggregated allele counts; they are amortised over a one-time import and grow in advantage with repeated queries. FULL PATH speedups (63–179× for iHS, XP-EHH, and nSL; 4.6× for ROH) reflect algorithmic optimisations—bit-packed haplotype matrices, SIMD kernels, and chunked processing—that apply from the first query. PLINK 2 was faster for pairwise LD (5.7×) owing to its native binary format. Both paths maintained constant ∼160 MB procedure working memory (JVM heap increment during query execution); the Neo4j database engine requires additional configured memory (4 GB heap + 16 GB page cache in our setup). By comparison, scikit-allel peaked at 1,051 MB and PLINK 2 at 1,665 MB of process memory. Full-genome analyses are detailed in Supplementary Notes S1–S3 (analysis protocols) and S2–S4 (deep investigations); import pipeline in Supplementary Note S5; EHH/ROH computation in Supplementary Note S6.

The query-time speedups in Table 1 are amortised over a one-time import step (∼12 min for chr22, ∼4.5 h for full genome). For a single first-time query, end-to-end wall-clock time is GraphPop 12.2 min (import + query) vs scikit-allel 29 min for diversity—a 2.4× net speedup even on first use. The amortised advantage becomes decisive for multi-query workflows: the full rice analysis (288 FAST PATH calls across 12 populations × 12 chromosomes) completes in 28 min of query time after a one-time import, compared to an estimated 144 h for the equivalent scikit-allel calls (288 calls × ∼30 min each; Supplementary Note S7).

## Discussion

GraphPop reduces the computational complexity of population genomics summary statistics from *O*(*V* × *N* ) to *O*(*V* × *K*). The *O*(*V* × *K*) complexity derives from pre-aggregating allele counts at import time—a strategy that could in principle be implemented in any storage system, including relational databases. What the graph model adds beyond pre-aggregation is threefold. First, *constant-time annotation conditioning*: traversing from a Variant node through a HAS_CONSEQUENCE edge to its Gene costs *O*(1) via index-free adjacency, compared to *O*(log *N* ) index lookups per join in a relational system; computing *π_N_ /π_S_* in a relational database requires four SQL operations across two tables (or a denormalized schema that must be maintained as annotations change), whereas in GraphPop it is a single edge traversal with an inline property read. Second, *co-located persistent results*: computed statistics are stored as properties on the same Variant and GenomicWindow nodes that carry the input data, enabling second-order queries (pathway co-selection correlations, multi-statistic convergence) without joins across result tables. Third, *schema-encoded biological relationships*: the variant→gene→pathway→GO term path is a first-class data structure, not a coordinate-based file intersection. Pre-aggregation provides the complexity reduction; the graph structure makes it composable.

This composability distinguishes GraphPop from a pre-computed lookup table. Each biological analysis presented here—universal *π_N_ /π_S_ >* 1.0 in cultivated rice, the cross-species Fst reversal, convergent sweep detection at *KCNE1*, pathway co-selection networks—required combining pre-aggregated counts with functional annotations, previously stored statistics, or both. They demonstrate what becomes practical when computation is decoupled from sample count and results persist as part of the data structure.

The *O*(*V* × *K*) complexity is a property of the pre-aggregation strategy, not of any specific database engine. GraphPop’s stored procedures depend only on the labelled property graph model and Cypher query language, both supported by multiple engines including the open-source distributed database NebulaGraph[26] and the managed service Neo4j AuraDB[27]. Migration requires only deployment configuration changes, not modifications to the analytical logic. The complexity advantage demonstrated here on datasets of 3,000 samples could therefore extend to biobank-scale datasets (*>* 100,000 individuals) through distributed graph database deployment, though this remains to be validated.

GraphPop is designed for the datasets that constitute the majority of population genomics research: crop breeding panels, livestock genetics, aquaculture, forest genomics, conservation biology, and non-biobank human studies, where sample counts range from hundreds to tens of thousands (Supplementary Note S9). At this scale, the *O*(*V* × *K*) advantage is fully realised on a single workstation.

Each existing tool excels at a different part of the problem. Hail[20], built on Apache Spark, provides horizontal scalability for biobank-scale cohorts (*>* 100,000 samples) and a flexible expression language for custom statistics; it is the tool of choice for datasets that exceed single-machine memory. Columnar stores (Zarr, TileDB) are well suited to dense array operations. Neither natively encodes the variant-to-gene-to-pathway relationships that enable annotation conditioning by traversal. PLINK 2[4] is faster than GraphPop for pairwise LD and for Fst alone (3.9 s vs 8.8 s on chr22) owing to its highly optimised binary format; it does not compute iHS, XP-EHH, nSL, Fay & Wu’s *H*, or Garud’s *H*, and has no annotation-conditioning mechanism. Pixy[28] handles missing genotypes rigorously when computing *π* and *D_xy_*, an area where GraphPop currently assumes complete data or uses allele-count-based estimators. scikit-allel[1] provides a flexible Python API that supports interactive, exploratory analysis on individual chromosomes. Tree-sequence methods (tskit[29], tsinfer) achieve efficient computation through genealogical compression, representing variation as a sequence of local trees rather than a genotype matrix. For statistics naturally expressed as tree operations (TMRCA-based diversity, coalescent-aware Fst), tree sequences can achieve sub-*O*(*V* × *N* ) complexity. GraphPop and tree-sequence approaches are complementary: tree sequences require phased data and inferred genealogies, and excel at coalescent-native statistics; GraphPop operates on called genotypes from standard VCF files, and excels when annotation conditioning, result persistence, and multi-statistic composition are priorities. The two approaches could in principle be combined, with tree-sequence statistics imported as node properties for graph-based composition. GraphPop complements all of these tools: it targets the regime where repeated annotation-conditioned queries across many populations are the bottleneck, and where results need to persist for downstream composition.

Several limitations should be noted. GraphPop requires a graph database deployment, adding infrastructure complexity relative to single-binary tools; the graphpop setup command reduces this to a single step that downloads and configures the database automatically. Neo4j Community Edition supports one active database at a time; multi-species comparative workflows require database switching via the CLI (graphpop db switch) or upgrade to Neo4j Enterprise Edition for concurrent multi-database access. Pairwise LD is slower than PLINK 2 with native binary files. PCA, kinship/IBS/IBD, demographic inference, and simulation interfaces are not yet implemented. The import pipeline requires pre-called VCF and VEP annotations; joint calling must be performed upstream. The *O*(*V* ×*K*) complexity applies to FAST PATH statistics after import; the import step itself is *O*(*V* × *N* ) and must be performed once per dataset. For FULL PATH statistics, runtime still depends on haplotype count, though bit-packing and SIMD provide substantial constant-factor reductions.

As population genomics datasets continue to grow—pangenome projects in crops, longitudinal monitoring in conservation, multi-generation breeding programmes—the cumulative model offered by the graph database provides an alternative to repeated ephemeral computation. Import once per dataset version, compute once, query indefinitely: new annotations or population definitions trigger only the affected statistics, not a full re-analysis. Incremental dataset updates—adding samples, populations, or annotation versions—are handled by the companion platform GraphMana[30]. Planned extensions include graph-native PCA and kinship matrices, demographic inference integration (dadi, fastsimcoal2) via the persistent joint SFS, and forward simulation interfaces (msprime, SLiM) that write simulated data directly to graph format for comparison with empirical results. A companion platform, GraphMana[30], addresses project-level data lifecycle—including incremental sample addition, cohort management, data provenance, and multi-format export—and is described separately.

## Online Methods

### Graph database technology

GraphPop is built on graph database technology, a class of database management systems that store data as nodes (entities) and edges (relationships) rather than in rows and columns[12, 13]. Graph databases differ from relational databases in three architecturally significant ways. First, *index-free adjacency*: each node physically stores direct pointers to its adjacent nodes, so traversing a relationship (e.g., from a Variant to its Gene via a HAS_CONSEQUENCE edge) is an *O*(1) pointer dereference, not an *O*(log *N* ) index lookup or *O*(*N* ) table scan as in relational systems[12]. Second, *labelled property graphs*: nodes and edges carry labels (types) and key-value properties, enabling a single data structure to represent both the topology of biological relationships and the quantitative attributes (allele frequencies, Fst values, selection scores) needed for statistical computation[14]. Third, *native graph query languages* (such as Cypher, GQL, or Gremlin) express multi-hop traversal patterns declaratively, making complex relationship queries (“find all variants in genes belonging to a specific pathway with iHS *>* 2”) concise and efficient.

For population genomics, these properties address the three architectural limitations of flat-file tools identified in the Introduction. Index-free adjacency makes annotation conditioning a constant-time traversal rather than a multi-step pipeline. The labelled property graph enables the persistent analytical record: computed statistics are stored as node properties on the same objects that carry the genotype data. And graph query languages make multi-statistic composition a declarative query rather than a custom script.

GraphPop’s current implementation uses Neo4j Community Edition[31] (version 2025.12.1) as its graph database engine, but the architecture is designed around the labelled property graph model and Cypher query language, both of which are supported by multiple graph database systems (Memgraph, Amazon Neptune with openCypher, TigerGraph, FalkorDB) and are being standardised as ISO GQL[32]. The stored procedures use the graph database’s native Java API for node and relationship traversal; the import pipeline, CLI, and MCP server communicate via the Bolt wire protocol, which is supported by all Cypher-compatible graph databases. Migration to alternative graph database engines requires no changes to the analytical logic, only to the deployment configuration.

### Implementation

GraphPop is a self-contained platform comprising four layers. The **compute engine** is a graph database stored procedure plugin in Java (JDK 21, ∼6,000 lines); the 12 procedures are compiled into a single JAR deployed to the graph database plugins directory. The **import pipeline** (graphpop-import) uses cyvcf2[33] for VCF parsing and numpy for vectorized genotype packing; VEP[6] consequence annotations, Reactome[15] and Plant Reactome[25] pathway data, GO term annotations, and ancestral alleles are loaded by dedicated parsers.

The **command-line interface** (graphpop CLI) provides 62 commands organised into 11 functional domains (Supplementary Fig. S1): infrastructure and server management (graphpop setup, start, stop, status), database lifecycle (import, dump, load, db subcommands), configuration and validation (config, validate, inventory), all 12 statistical procedures as shell commands with default TSV output (graphpop ihs chr22 EUR --output ihs.tsv), annotation-based filtering of persisted results (graphpop filter ihs chr22 EUR --consequence missense variant), multi-statistic convergence detection (graphpop converge), gene ranking (graphpop rank-genes), graph exploration (graphpop lookup, neighbors), data extraction and BED export (graphpop extract, export-bed), orchestration and aggregation (graphpop run-all, batch, aggregate, report), and 11 publication-ready visualisation types (graphpop plot). An optional --persist flag writes results to graph nodes for cross-query; without it, the tool behaves like a conventional command-line program. Users require no knowledge of graph query languages, graph theory, or database internals.

A **programmatic interface** via the Model Context Protocol (MCP) exposes 21 tools—the 12 procedures plus convergence detection, gene ranking, annotation lookup, filtering, inventory, status, and arbitrary graph queries—to large language model agents and external applications. The MCP server connects to the graph database via the same configuration as the CLI (shared GRAPHPOP URI / GRAPHPOP DATABASE environment variables) and returns JSON results.

The **analysis pipelines** orchestrate full-genome runs across all populations and chromosomes (graphpop run-all), aggregate results into summary tables (graphpop aggregate), and generate automated HTML reports (graphpop report). All scripts used for the analyses reported in this paper are included in the repository. All code is available under the MIT license.

### Graph schema

The graph database schema comprises Variant, Sample, Population, Chromosome, Gene, Pathway, GOTerm, and GenomicWindow node labels with NEXT, HAS_CONSEQUENCE, IN_PATHWAY, HAS_GO_TERM, IN_POPULATION, ON_CHROMOSOME, and LD relationship types. Variant nodes are indexed on (chr, pos) for range queries. Population harmonic numbers (*a_n_* = *H*(2*n* − 1), *a_n_*_2_ = *H*_2_(2*n* − 1) for *n* diploid samples) are stored on Population nodes and used by diversity and SFS procedures. The schema is intentionally generic: it encodes the universal structure of population genomic data (variants, samples, populations, annotations) without assumptions about species, ploidy, or annotation source.

### Packed genotype encoding

Genotypes are stored as gt packed (2 bits/sample, encoding: 00=HomRef, 01=Het, 10=HomAlt, 11=Missing) and phase packed (1 bit/sample, which haplotype carries ALT). The cyvcf2 genotype encoding is remapped during import (0→0, 1→1, 2→3, 3→2) to match the internal 2-bit scheme. For non-diploid samples, ploidy packed (1 bit/sample) indicates ploidy; null implies all-diploid for backward compatibility. PackedGenotypeReader extracts genotype and phase values using bit-shift operations.

### FAST PATH statistics

**Diversity.** Nucleotide diversity *π* = Σ*_i_*2*p_i_*(1 − *p_i_*)· *n_i_/*(*n_i_* − 1) summed over variants, where *p_i_* = AC*_i_/*AN*_i_* and *n_i_* = AN*_i_/*2 for the focal population. Watterson’s *θ_W_* = *S/a_n_* where *S* is the number of segregating sites. Ta_Σ_jima’s *D* follows the standard formulation[34]. Fay & Wu’s *H* = *π* − *θ_H_* where *θ_H_* = Σ2*i*^2^*/*(*n*(*n* − 1)) is the squared-frequency estimator; the normalized version uses the Zeng et al. (2006) variance[35].

**Divergence.** Hudson’s Fst uses ratio-of-averages: Fst = Σ*N_i_/ΣD_i_*. Weir & Cockerham (1984) Fst[11] decomposes into variance components (*a, b, c*); Fst = Σ*a/* Σ(*a* + *b* + *c*). *D_xy_* = Σ*p*_1_(1 − *p*_2_) + *p*_2_(1 − *p*_1_). PBS (Population Branch Statistic)[36] uses *T* = − log(1 − Fst_WC_) for three population pairs.

**SFS.** Site frequency spectrum uses allele counts; unfolded SFS polarizes using ancestral_allele annotations where available, with n_polarized reporting coverage.

### FULL PATH statistics

**HaplotypeMatrix.** A dense bit-packed matrix loaded in one pass from Variant node arrays. Layout: haplotypes[variantIdx][byteIdx], with haplotype *h* accessed as (haplotypes[v][h>>3] >> (h&7)) & 1. Row length is (nHaplotypes + 7)*/*8 bytes. loadPair() loads two populations from the same variant pass for XP-EHH.

**LD.** Pearson *r*^2^ computed from haplotype dosages via VectorOps dot product. *D^′^* computed from 2×2 haplotype tables via Integer.bitCount() on packed byte arrays. LD edges are written between Variant nodes above a configurable *r*^2^ threshold.

**EHH-based statistics.** EHHComputer walks position-sorted haplotype arrays from a focal variant outward, tracking distinct haplotype classes via flat arrays with incremental sum-of-pairs updates. iHH is the trapezoidal integral of the EHH decay curve. iHS[8] = log(iHH_ancestral_*/*iHH_derived_), standardized within allele-frequency bins (default 20). XP-EHH[9] = log(iHH_pop1_*/*iHH_pop2_), genome-wide standardized. nSL[37] uses mean pairwise shared suffix length (SSL) per allele class. All EHH procedures are configurable: min_ehh (truncation threshold), max_gap (200 kb abort), gap_scale (20 kb cap).

**Garud’s** *H*.[10] Haplotype frequencies computed by hashing within sliding windows. 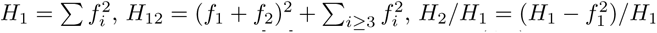.

**ROH.** Two-state Viterbi HMM[17] with autozygous (AZ) and Hardy–Weinberg (HW) states. Emissions are AF-weighted: *P* (hom|AZ) = 1−*ɛ*, *P* (hom|HW) = *p*^2^+(1− *p*)^2^ where *p* is the population allele frequency. Transitions use continuous-time Markov chain (CTMC) matrix power for inter-variant distances, matching bcftools roh[17]. Default parameters: hw_to_az = 6.7 × 10*^−^*^8^, az_to_hw = 5 × 10*^−^*^9^, het_error_rate = 10*^−^*^3^. GraphPop provides two ROH codepaths: a graph-database Viterbi HMM that traverses CARRIES edges (128 s for chr22), and a numpy-accelerated path that reads gt packed arrays directly (10.9 s). Table 1 reports the numpy path; the graph-database path is documented in the supplementary benchmark table.

### VariantFilter and VariantQuery

All procedures share a common filtering infrastructure. VariantFilter supports min af, max_af, min_call rate, hwe_pvalue (Wigginton et al. 2005 exact test[38]), and variant_type. VariantQuery dynamically constructs graph queries that optionally traverse HAS_CONSEQUENCE edges (for consequence filtering) or IN_PATHWAY edges (for pathway filtering) before the main variant scan. Custom sample subsets are supported via samples/samples1/samples2 options using SampleSubsetComputer for FAST PATH or HaplotypeMatrix.load() overloads for FULL PATH.

### Datasets and annotation pipeline

**1000 Genomes Project.** Phase 3 chr22 VCF (3,202 samples, 1,070,401 variants, 5 superpopulations: AFR, AMR, EAS, EUR, SAS)[18]. Used for benchmarking and validation. Full-genome analysis was also performed across all 22 autosomes for the 26 sub-populations (3,899 per-population analyses).

**3K Rice Genomes Project.** 3,024 accessions, ∼29.6M SNPs, 12 chromosomes, 12 subpopulations following the Wang et al. (2018) ADMIXTURE K=9 classification[19]. Subpopulation labels: indica (XI-1A, XI-1B, XI-2, XI-3, XI-adm), japonica (GJ-tmp, GJ-trp, GJ-sbtrp, GJ-adm), aus (cA-Aus), basmati (cB-Bas), and admix.

**Functional annotations.** Variant consequences were annotated with VEP[6] (release 110) using the Ensembl human GRCh38 cache for 1000 Genomes and the *Oryza sativa* IRGSP-1.0 reference for rice 3K, with default options and --pick canonical transcript selection. Reactome pathway-gene mappings[15] (release 86, 2,241 human pathways) and Plant Reactome[25] (153 *O. sativa* pathways) were loaded as TSV files into Pathway nodes connected to Gene nodes via IN_PATHWAY edges. GO term annotations were loaded from Ensembl BioMart. Ancestral alleles were derived from the Ensembl EPO multi-species alignment FASTA files (6-primate alignment for human, 9-grass alignment for rice); the polarization fraction (proportion of variants with confident ancestral state) ranged from 85% to 92% across chromosomes.

### Multi-statistic convergence detection

Convergent positive selection was identified by querying stored Garud’s *H*_12_ values on Variant nodes that were materialised by genome scan procedures. The graph pattern selects variants where *H*_12_ *>* 0.3 in any continental group, traverses HAS_CONSEQUENCE edges to associated Gene nodes, and counts the number of distinct continental groups in which each gene shows the signal. Genes detected in ≥ 2 independent groups were retained as convergent candidates. The *H*_12_ *>* 0.3 threshold corresponds to the upper ∼5% of the genome-wide *H*_12_ distribution and is the standard cutoff for hard-sweep candidates[10]. The five-group co-occurrence at *KCNE1* required *H*_12_ *>* 0.3 in AFR, AMR, EAS, EUR, and SAS independently; the conditional probability of this signal under a neutral null is *<* 10*^−^*^6^ assuming 5% per-population false positive rate.

### Pathway co-selection network

Pathway-level Fst was computed by traversing IN_PATHWAY edges from Pathway to Gene to Variant nodes via HAS_CONSEQUENCE, then averaging stored per-variant Fst values across all variants assigned to each pathway. For rice, this was done for all 66 unique pairs of the 12 subpopulations and 153 Plant Reactome pathways; for human, all 5 representative continental pairs and 2,232 Reactome pathways. Spearman correlation *ρ_ij_* between pathways *i* and *j* was computed across the population-pair vector. Pairs with *ρ >* 0.7 were retained as edges in the co-selection network. Connected component analysis identified three modules in both species. The module structure was visualised using a force-directed layout (igraph). Hub degree per pathway was computed as the number of significant correlations within its module; the housekeeping module showed the highest mean hub degree in both species.

### Sensitivity analysis for rice *π_N_ /π_S_*

To verify that the universal *π_N_ /π_S_ >* 1.0 finding was not driven by methodological artifacts, four robustness checks were performed (Supplementary Note S7): (1) variant frequency filtering at MAC *>* 1 (singleton removal) and MAC *>* 2 (singletons and doubletons removed); (2) call rate filtering at *>* 90% (excluding sites with *>* 10% missing data); (3) per-chromosome decomposition (144 population × chromosome combinations independently checked); (4) annotation comparison using VEP missense variant versus nonsynonymous variant terms. The qualitative finding (all 12 subpopulations *π_N_ /π_S_ >* 1.0) was confirmed under all four checks; quantitative ratios shifted by *<* 5%.

### Statistical testing

Differences in mean Fst between VEP impact classes were tested with the two-sided Mann–Whitney *U* test, treating each variant as an independent observation. For the rice GJ-tmp vs XI-1A comparison, the test was applied pairwise between HIGH (n = 18,284 variants), MODERATE (n = 26,318), and LOW (n = 33,710) classes, with Bonferroni correction across the three pairwise tests. The null hypothesis of equal Fst distributions was rejected at *p <* 10*^−^*^150^ for all pairwise comparisons. For the human YRI vs CEU comparison, *p* ≈ 0 at machine precision. Correlations between continuous variables (e.g., FROH vs heterozygosity rate) used Spearman’s rank correlation; reported *p*-values are two-tailed.

### Benchmark methodology

Each benchmark was run 3 times with 1 warm-up run discarded. GraphPop was called via the Python driver with the graph database pre-loaded in memory (warm cache). Competitor tools read from their native file formats (VCF for VCFtools, scikit-allel, bcftools; PLINK binary for PLINK 2 and PLINK 1.9). Peak memory was measured using GraphPop’s graphpop.heap stats() procedure (JVM heap) and /usr/bin/time -v (resident set size) for external tools. Runtime comparisons used wall-clock time including I/O. Competitor tool versions: scikit-allel v1.3.7[1], VCFtools v0.1.16[2], PLINK 2 v2.00a6[4], PLINK 1.9 v1.90b7.2[3], bcftools v1.21[17].

All tool versions reflect the latest stable releases available at the time of analysis (January 2026). Benchmark scripts are provided in the repository for re-execution with updated tool versions. Full benchmark protocol, region definitions, and end-to-end (import + query) versus amortised (query-only) timing comparisons are described in Supplementary Note S7.

### Software versions and reproducibility

GraphPop v0.1.0 was implemented in Java 21 (JDK build 21.0.6) for the stored procedures and Python 3.11 for the import pipeline, CLI, and MCP server. The graph database engine was Neo4j Community Edition v2025.12.1[31] with 4 GB heap and 16 GB page cache. Pre-built Neo4j databases for both datasets, with all computed statistics persisted, are available on Zenodo (rice 3K: https://doi.org/10.5281/zenodo.19471968; human 1000G: https://doi.org/10.5281/zenodo.19472010). End-to-end analysis scripts, source data for all figures, and the GraphPop CLI are available at https://doi.org/10.5281/zenodo.19471963. The complete software stack can be installed from PyPI (pip install graphpop-cli); the CLI’s setup command automatically downloads Neo4j and the pre-compiled procedures plugin. Two end-to-end tutorials (rice 3K and human 1000 Genomes) are provided as vignettes in the source repository. Expected runtimes on the hardware described below: rice 3K full analysis ∼2 hours; human 1000 Genomes full analysis ∼112 hours.

### Hardware

All benchmarks and analyses were performed on a single workstation with an AMD processor, 64 GB RAM, NVMe SSD storage, and an NVIDIA RTX 4090 GPU (not used by current procedures). The graph database was configured with 4 GB heap and 16 GB page cache.

## Data Availability

Pre-built Neo4j graph databases with all computed statistics are deposited on Zenodo: human 1000 Genomes (https://doi.org/10.5281/zenodo.19472010; 31 GB, split into parts) and rice 3K (https://doi.org/10.5281/zenodo.19471968; 14 GB). Analysis results, figure source data, scripts, and supplementary tables are available at https://doi.org/10.5281/zenodo.19471963. The source VCF data are available from the 1000 Genomes Project (https://www.internationalgenome.org/) and the 3K Rice Genomes Project (https://snp-seek.irri.org/; Wang et al. 2018). Reactome[15] and Plant Reactome[25] pathway annotations are from https://reactome.org/ and https://plantreactome.gramene.org/. Source data are provided with this paper.

## Code Availability

GraphPop source code is available at https://github.com/ jfmao/GraphPop under the MIT license. The repository includes the 12 stored procedures (Java), the import pipeline (Python), the graphpop command-line interface (62 commands organised into 11 functional domains; Supplementary Fig. S1), the MCP server for AI agent access, all analysis scripts and vignettes used in this paper, and Docker support for containerised deployment (documented in the installation guide).

## Supporting information

all supplementary materials

## Acknowledgements

This work was partially supported by the Wallenberg Initiatives in Forest Research (WIFORCE) funded by the Knut and Alice Wallenberg Foundation. Genomic data processing and analyses were performed using resources provided by the Swedish National Infrastructure for Computing (SNIC), through the High Performance Computing Centre North (HPC2N) at Umeå University. We thank the 1000 Genomes Project Consortium and the 3K Rice Genomes Project for making their data publicly available.

## Declarations

- **Funding:** Wallenberg Initiatives in Forest Research (WIFORCE), Knut and Alice Wallenberg Foundation.
- **Competing interests:** The authors declare no competing interests.
- **Ethics approval and consent to participate:** Not applicable.
- **Consent for publication:** Not applicable.
- **Author contribution:** Conceptualization and design: J.F.M. and E.E. Java stored procedures and SIMD kernels: E.E. Python CLI, import pipeline, and MCP server: E.E. and S.W.Z. Biological analyses (rice 3K): Z.Y.C. and S.N. Biological analyses (human 1000G): E.E. and S.W.Z. Benchmarking: E.E. Writing—original draft: J.F.M. and E.E. Writing—review & editing: all authors.

## Supplementary Notes

*Note S1* Human 1000 Genomes Full-Genome Analysis.

*Note S2* Human 1000 Genomes Deep Investigations.

*Note S3* Rice 3K Full-Genome Analysis.

*Note S4* Rice 3K Deep Investigations.

*Note S5* Import Pipeline and Database Construction.

*Note S6* EHH and ROH Computation Details.

*Note S7* Benchmark Methodology.

*Note S8* GraphPop CLI Architecture.

*Note S9* Dataset Scale and Applicability.

## Supplementary Information

### Supplementary Note S1: Human 1000 Genomes Full-Genome Analysis

#### Analysis scope and comparison with classical approaches

The full-genome analysis described here would require, in a classical pipeline, running each statistic separately per tool (scikit-allel for diversity/iHS, VCFtools for Fst, PLINK for ROH, selscan for XP-EHH, a custom script for Garud’s *H*), then manually merging outputs across 22 chromosomes and 26 populations by genomic coordinates—a workflow involving thousands of separate tool invocations and hundreds of intermediate files. In GraphPop, all statistics are computed by 12 stored procedures that write results directly to graph nodes, creating a persistent analytical record where every result is immediately queryable and cross-referenceable without re-computation or file merging.

GraphPop was applied to the complete 1000 Genomes Project Phase 3 dataset[18] across all 22 autosomes (chr1–22) for 26 sub-populations grouped into 5 superpopulations: AFR (7 populations, 661 samples), AMR (4 populations, 347 samples), EAS (5 populations, 504 samples), EUR (5 populations, 503 samples), and SAS (5 populations, 489 samples). The full graph database contains ∼70.7M Variant nodes, 3,202 Sample nodes, 91,973 Gene nodes, 1.46M GenomicWindow nodes, and 2,241 Reactome pathways[15] connected via ∼3.8M HAS_CONSEQUENCE edges and ∼4,900 IN_PATHWAY edges.

#### Phase 1: Per-population statistics

For each of 26 populations × 22 chromosomes (572 runs), the following procedures were executed: graphpop.diversity (*π*, *θ_W_* , Tajima’s *D*, Fay & Wu’s *H*, *H_e_*, *H_o_*, *F_IS_*), graphpop.sfs (folded and unfolded site frequency spectra), graphpop.genome_scan (100 kb sliding windows; *π*, *θ_W_* , Tajima’s *D*, Fst, PBS), graphpop.ihs (integrated haplotype score, AF-bin standardized), graphpop.nsl (number of segregating sites by length), graphpop.garud_h (haplotype homozygosity statistics), and graphpop.roh (HMM-based runs of homozygosity). All results were written to the graph as persistent node properties.

Summary diversity statistics across superpopulations confirm expected continental patterns:

- **African populations** show the highest diversity (mean *π* = 0.051, *θ_W_* = 0.061), consistent with the larger long-term effective population size. MSL is the most diverse single population (*π* = 0.052).
- **Out-of-Africa populations** show progressively reduced diversity: SAS (*π* = 0.039), AMR (*π* = 0.040), EUR (*π* = 0.038), EAS (*π* = 0.036, lowest; PEL is the least diverse population at *π* = 0.036).
- **Tajima’s** *D* varies by continent: AFR (*D* = −0.52, population expansion), AMR (−0.39), SAS (−0.07), EUR (+0.09), EAS (+0.20, population structure/admixture).
- **FROH** is lowest in African populations (∼0.005) and highest in South Asian populations (∼0.014), consistent with documented consanguinity practices.

#### Phase 2: Pairwise statistics

For 18 key population pairs × 22 chromosomes (396 runs), XP-EHH was computed via graphpop.xpehh. For all 325 unique population pairs × 22 chromosomes (7,150 runs), divergence statistics (Hudson Fst, W&C Fst, *D_xy_*, *D_a_*) were computed via graphpop.divergence. Pairwise Fst values range from 0.001 (CEU vs GBR, within-EUR) to 0.167 (MSL vs CHS, the maximum intercontinental divergence). Within-continent pairs are tightly clustered: EUR (CEU–GBR Fst = 0.001), EAS (CHB– CHS = 0.002), SAS (STU–ITU = 0.002).

#### Phase 3: Ancestral allele analyses

Ancestral allele annotations from Ensembl EPO multi-species alignment FASTA files were loaded onto Variant nodes. Fay & Wu’s *H* was computed genome-wide for all populations. Polarized (unfolded) SFS was generated where ancestral state was available, with the fraction of polarizable sites reported per chromosome (∼85–92% depending on chromosome).

#### Phase 4: Population-specific analyses

PBS genome scans and Garud’s *H* scans were computed for 6 representative populations (YRI, CEU, CHB, GIH, JPT, LWK) to identify population-specific selection signals. Known selection targets were recovered:

- *LCT* /*MCM6* (lactase persistence, chr2:136 Mb): XP-EHH = −7.05 (YRI vs CEU); Garud’s *H*_12_ detected in CEU, FIN, GBR
- *SLC24A5* (skin pigmentation, chr15:48 Mb): XP-EHH = −5.19 (YRI vs CEU); also detected in SAS populations (GIH, PJL, ITU)
- *ADH1B* (alcohol metabolism, chr4:99 Mb): hard sweeps in JPT (*H*_12_ = 0.419) and CHB (*H*_12_ = 0.373); XP-EHH = −5.41 (CEU vs CHB)
- *FADS1* /*FADS2* (fatty acid desaturation, chr11:61 Mb): detected in EAS and AFR populations
- *LARGE1* (Lassa fever resistance, chr22:33 Mb): XP-EHH = 7.00 (YRI vs LWK)
- *HBB* (malaria resistance, chr11:5 Mb): XP-EHH = 6.42 (YRI vs LWK)
- HLA region (chr6:28–34 Mb): multi-population XP-EHH peaks (|XP-EHH| = 5–20)

#### Computation time

The full-genome analysis required approximately 112 hours of total wall-clock time on a single workstation (see Online Methods for hardware). Phase 1 (per-population: 3,899 analyses at ∼103 s mean) dominated at ∼112 hours; Phase 2 (pairwise divergence and XP-EHH) required ∼20 hours; Phases 3–4 (ancestral analyses and PBS) required ∼5 hours. All results are permanently stored in the graph database and require no re-computation for downstream queries.

### Supplementary Note S2: Human 1000 Genomes Deep Investigations

The following investigations exploit the persistent analytical record created by the full-genome analysis (Supplementary Note 1). Each investigation queries already-computed results stored on graph nodes; no statistics are re-computed. In a classical workflow, each of these analyses would require: (1) loading results from multiple output files into a common framework, (2) coordinate-matching across different tools’ output formats, and (3) writing custom scripts for each cross-statistic query. The graph database eliminates all three steps (Supplementary Table S9).

#### Pathway-level population differentiation

All 2,241 Reactome pathways were ranked by mean Hudson Fst (YRI vs CEU) via a single graph traversal through IN_PATHWAY → Gene → HAS_CONSEQUENCE → Variant edges. The top-10 pathways include:

1. Oculocutaneous albinism type I (Fst = 0.452; driven by *SLC24A5* )
2. TREK/TWIK potassium channels (Fst = 0.162)
3. Phase 3 rapid repolarisation (Fst = 0.133)
4. Voltage-gated potassium channels (Fst = 0.128)

The clustering of cardiac ion channel pathways in the top quartile implicates *KCNE1*, *KCNE4*, *KCNH2*, and *KCNQ1* as population-differentiated cardiac loci (Supplementary Fig. S2).

#### Convergent positive selection

The 9 genes with Garud *H*_12_ *>* 0.3 in ≥2 independent continental groups were identified by querying stored Garud *H* statistics on GenomicWindow nodes and following HAS_CONSEQUENCE edges to Gene nodes:

1. ***KCNE1*** : 5 continental groups (AFR, AMR, EAS, EUR, SAS; 6 populations; max
*H*_12_ = 1.0)
2. LOC102724560: 4 groups (AMR, EAS, EUR, SAS; max *H*_12_ = 0.90)
3. LOC107987288: 4 groups (max *H*_12_ = 0.83)
4. LOC102724428: 4 groups (max *H*_12_ = 0.81)
5. LOC124900992: 3 groups (max *H*_12_ = 0.68)
6. LOC124900995: 3 groups (max *H*_12_ = 0.68)
7. LINC01678: 2 groups (max *H*_12_ = 0.71)
8. LOC102725065: 2 groups (max *H*_12_ = 0.71)
9. CRYAA: 2 groups (AMR, SAS; max *H*_12_ = 0.55)

*KCNE1* was the only gene showing sweep evidence in all five continental groups, with an elevated between-group:within-group Fst ratio of 0.296—the signature of an ancient, pre-Out-of-Africa selective sweep. The Phase 3 rapid repolarisation and Phase 2 plateau phase pathways containing *KCNE1* showed significant Fisher enrichment (*p* = 0.0007 and *p* = 0.001, respectively).

#### Consequence-level selection bias

Fst values stored on Variant nodes were stratified by VEP-annotated impact class via HAS_CONSEQUENCE edge traversal. HIGH-impact variants (missense, stop-gain) showed significantly lower mean Fst (0.124) than LOW-impact variants (0.143; Mann– Whitney *U* , *p* ≈ 0), confirming genome-wide purifying selection on functionally deleterious alleles at the population level.

#### Individual-level evolutionary trajectory

Genome-wide FROH was computed per sample from ROH HMM checkpoints (3,094 of 3,202 samples with ROH data). Chromosome-22-based individual features (heterozygosity rate, rare variant burden, rare missense burden) were extracted from pre-exported haplotype arrays. Five evolutionary process statistics per individual were used for PCA and UMAP embedding (Supplementary Fig. S4):

- PC1 (57.0% variance): diversity axis—African samples highest on heterozygosity and rare burden
- PC2 (30.9% variance): inbreeding axis—South Asian samples highest on FROH
- FROH vs heterozygosity rate: *ρ* = −0.394, *p* = 1.2 × 10*^−^*^29^
- Individual PC1 vs population PC1: *ρ* = 0.74 (Spearman), validating the approach

Per-group mean statistics (chr22 features):

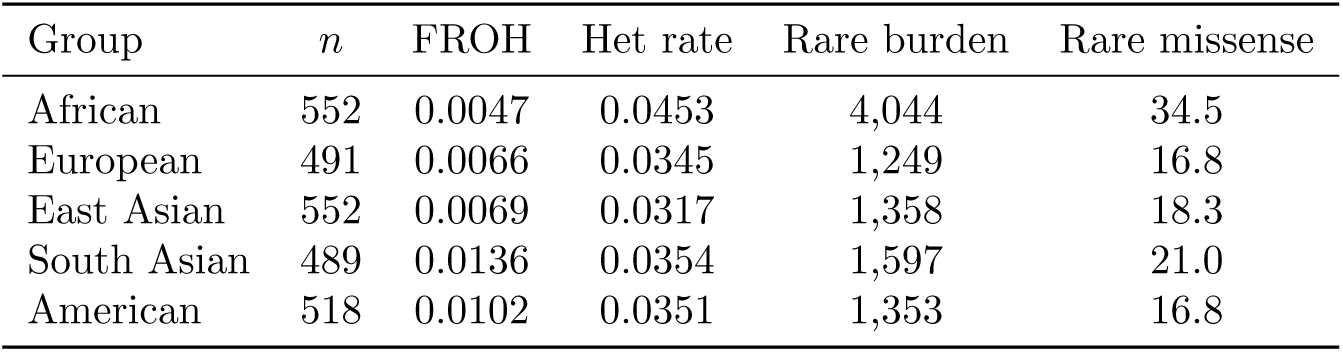

#### Outlier analysis

Z-distance from population centroid in the 5-dimensional process space identified extreme individuals. The top 5 outliers:

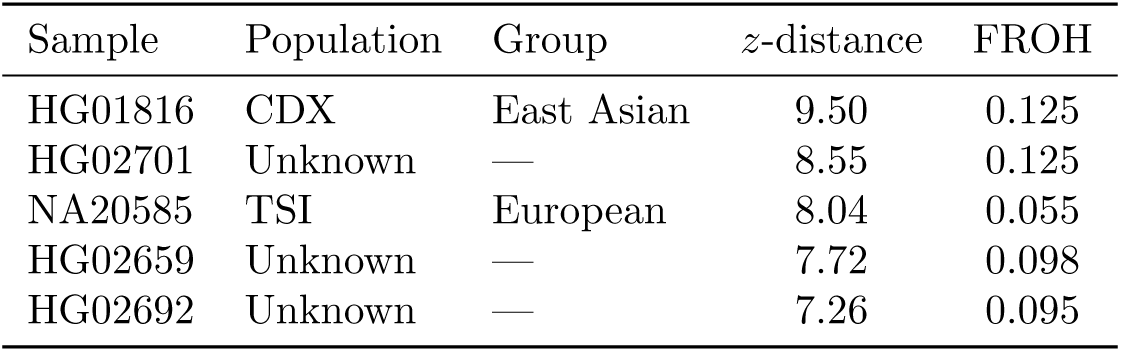

The top outlier (HG01816) carries FROH = 0.125—18× the CDX population mean— consistent with recent close-relative mating. Four UNKNOWN-population samples appeared in the top 10, suggesting mislabeling or recent admixture. Z-distance weakly predicts FROH (*ρ* = −0.087; Supplementary Fig. S3).

#### Pathway-stratified purifying selection gradient

Genome-wide missense variant positions (224,181 across 22 autosomes) were classified as belonging to constrained pathways (lowest-quartile mean Fst: ≤ 0.108; *n* = 15,092 sites) or divergent pathways (highest-quartile Fst: ≥ 0.158; *n* = 21,886 sites) via IN_PATHWAY graph traversal (Supplementary Fig. S5). Key findings:

- African 2× enrichment holds in both constrained (2.04×, *p* = 1.6 × 10*^−^*^290^) and divergent (1.95×, *p* = 3.0 × 10*^−^*^265^) pathways
- FROH predicts individual rare missense burden: *ρ* = −0.211, *p* = 1.6 × 10*^−^*^33^

(genome-wide)

- FROH does *not* predict constrained:divergent burden ratio: *ρ* = +0.004, *p* = 0.84
- Conclusion: bottleneck-driven depletion of rare functional variants is genome-wide and uniform across pathway classes

#### *π_N_ /π_S_* across human populations

Annotation-conditioned diversity was computed for all 26 populations using graphpop.diversity with consequence: ‘missense variant’ and ‘synonymous variant’. All 26 populations show *π_N_ /π_S_ <* 1.0 (range: 0.646–0.673), confirming efficient genome-wide purifying selection on protein-coding variants. African populations show the lowest ratios (MSL: 0.646, LWK: 0.649, YRI: 0.650), consistent with larger effective population size supporting more efficient purifying selection. Non-African populations show modestly elevated ratios (CHS: 0.673, CEU: 0.670, IBS: 0.670), consistent with the well-documented out-of-Africa bottleneck reducing purifying selection efficiency by ∼3.5%.

This pattern is the *inverse* of the rice result (*π_N_ /π_S_ >* 1.0 in all 12 subpopulations). The contrast demonstrates that annotation-conditioned analysis reveals fundamentally different selection regimes: in natural populations (human), purifying selection efficiently removes deleterious protein-coding variants; in domesticated populations (rice), bottlenecks and artificial selection have relaxed this constraint. This cross-species comparison is uniquely enabled by GraphPop’s annotation conditioning, applied identically to both datasets.

#### Pathway co-selection network

Pathway-level Fst values stored across 5 population pairs (YRI vs CEU, YRI vs CHB, CEU vs CHB, YRI vs JPT, CEU vs JPT) were correlated pairwise across 2,232 Reactome pathways. Using Spearman *ρ >* 0.7 as threshold, the pathway co-selection network contains 1,182,891 edges (939,959 positive, 242,932 negative). Community detection identified three functional modules: (1) a large module of 602 pathways enriched for DNA repair, apoptosis, and opioid signaling; (2–3) two modules enriched for immune signaling, neuronal transmission, and metabolic processes. The hub pathways (highest degree) include DNA repair and ERBB4 signaling, indicating that core cellular maintenance pathways differentiate in concert across human population pairs—paralleling the rice finding where core cellular machinery formed the largest co-selection module. This convergence across species suggests that housekeeping pathway co-differentiation is a general feature of population divergence, not specific to domestication.

### Supplementary Note S3: Rice 3K Full-Genome Analysis

#### Analysis scope and comparison with classical approaches

The rice 3K analysis[19] demonstrates GraphPop’s species-agnostic design: the same 12 procedures used for human data operate on rice without modification. The classical equivalent for the rice dataset would require: VCFtools or scikit-allel for diversity/Fst (separate runs per population and chromosome), selscan for XP-EHH, PLINK for ROH, custom scripts for Garud’s *H*, and VEP for consequence annotation—with all results stored in separate files requiring manual coordination. GraphPop replaces this multi-tool pipeline with a single import followed by graph procedure calls, with all results persisted as queryable node properties.

The rice 3K graph database contains 29,635,224 Variant nodes, 3,024 Sample nodes, 13 Population nodes (12 subpopulations + 1 unassigned), 55,328 Gene nodes, 664 Plant Reactome pathways[25], 2,560 GO terms, and 89,573 GenomicWindow nodes. Functional annotations comprise 7,004,802 HAS_CONSEQUENCE edges (SnpEff with MSU7 gene models), 1,296 IN_PATHWAY edges, and 16,030 HAS_GO_TERM edges. Ancestral allele annotations were derived from rice EPO multi-species alignment FASTA files.

#### Phase 1: Per-population statistics

For each of 12 populations × 12 chromosomes (144 runs), the same 7 procedures as the human analysis were executed. Genome-wide diversity statistics are presented in Supplementary Table S2. Key patterns:

- Indica subpopulations are 2–3× more diverse than japonica (admix *π* = 0.078 vs GJ-tmp *π* = 0.019)
- All populations show negative Tajima’s *D* (−0.86 to −2.41), indicating population expansion or selective sweeps
- Inbreeding coefficients (*F_IS_*) are high across all subpopulations (0.54–0.72), reflecting the predominantly self-pollinating mating system of cultivated rice
- GJ-tmp shows the most extreme bottleneck signature: lowest diversity (*π* = 0.019), most negative Tajima’s *D* (−2.41), highest *F_IS_* (0.72), and highest FROH (0.080)

Total Phase 1 computation time: ∼48 minutes on a single workstation.

#### Phase 2: Pairwise statistics

XP-EHH was computed for 6 key population pairs × 12 chromosomes (72 runs). Divergence statistics were computed for all 66 unique pairs × 12 chromosomes (792 runs). Pairwise W&C Fst values range from 0.014 (XI-adm vs admix) to 0.710 (GJ-tmp vs XI-1A), with the top 10 pairs listed in Supplementary Table S5. The UPGMA tree from the pairwise Fst matrix cleanly recovers the indica–japonica split with japonica subpopulations forming a distinct clade.

Total Phase 2 computation time: ∼78 minutes.

#### Interpretation analyses

Following the per-population and pairwise phases, annotation-conditioned analyses were executed: annotation loading (VEP consequence types, Plant Reactome pathways, GO terms), *π_N_ /π_S_* computation for all 12 populations, conditioned genome scans (missense vs synonymous Fst), ROH HMM computation, and Fst tree construction. These results were written to the graph as permanent properties, enabling all downstream deep investigations (Supplementary Note 4) without re-computation.

### Supplementary Note S4: Rice 3K Deep Investigations

Nineteen deep investigations (R01–R19) exploit the persistent analytical record from the rice full-genome analysis. Investigations R01–R12 operate on JSON-exported summary statistics and are in principle reproducible with classical tools, though less efficiently (each requires manual extraction and coordination of results from multiple tool outputs). Investigations R13–R19 query the graph database directly, traversing HAS_CONSEQUENCE and IN_PATHWAY edges to perform analyses that have no practical classical equivalent—they require simultaneous access to per-variant Fst, consequence annotations, pathway membership, and population-level statistics in a single query framework.

#### R01: Evolutionary fingerprint PCA

PCA on 7 population-level statistics (*π*, *θ_W_* , Tajima’s *D*, *F_IS_*, FROH, number of sweeps, hard sweep fraction) across 12 populations yielded PC1 (51.7% variance) as the indica–japonica divergence axis and PC2 (24.3%) as an inbreeding/bottleneck axis. GJ-tmp occupies an extreme position on both axes.

#### R02: *π_N_ /π_S_* analysis

All 12 subpopulations show *π_N_ /π_S_ >* 1.0 (range: 1.018–1.146; Supplementary Table S3), demonstrating universal relaxation of purifying selection. Admixed groups show the highest ratios (admix: 1.146), while GJ-tmp shows the lowest (1.018), as described in the main text.

#### R03: Pairwise Fst landscape

66 population pairs were ranked by genome-wide mean W&C Fst (Supplementary Table S5). The indica–japonica split dominates: all 10 highest-Fst pairs involve a japonica and an indica subpopulation, with GJ-tmp consistently showing the highest divergence.

#### R04: Conditioned Fst genome scans

Missense and synonymous Fst were computed genome-wide for three major population pairs (Supplementary Table S4). The missense:synonymous Fst ratio exceeds 1.0 in all three, with the strongest signal at the temperate–tropical japonica boundary (1.106), as described in the main text.

#### R05: Domestication gene convergence

16 known domestication genes were examined for selection signals (XP-EHH, Garud’s *H*_12_). 7 showed sweep evidence in at least one population, with *GW5*, *Hd1*, and *PROG1* detected by XP-EHH and *OsC1*, *Wx*, and *DRO1* detected by Garud’s *H*_12_ (Supplementary Table S6).

#### R06: ROH/inbreeding landscape

Genome-wide FROH was computed for all 12 populations. GJ-tmp shows the highest inbreeding (FROH = 0.080), consistent with its severe bottleneck. Indica populations show lower FROH (0.015–0.035), reflecting larger effective population sizes. The predominantly selfing mating system of rice contributes to elevated FROH across all subpopulations relative to human populations.

#### R07: PBS population-specific sweeps

PBS genome scans (100 kb windows) identified population-specific selection signals for 6 representative populations. The strongest signal was in GJ-tmp at Chr5:28.2 Mb (PBS = 2.76), consistent with temperate adaptation. PBS peaks in indica populations were generally weaker, reflecting larger *N_e_* and less extreme allele frequency shifts.

#### R08: Sweep classification

Sweeps were classified as hard (Garud’s *H*_12_ *>* 0.3 and *H*_2_*/H*_1_ *<* 0.1) or soft (*H*_12_ *>* 0.3 and *H*_2_*/H*_1_ ≥ 0.1). Of 8 detected sweeps, 5 were hard and 3 soft, all in japonica subpopulations—consistent with the stronger bottleneck history reducing standing variation available for soft sweeps in japonica.

#### R09: Multi-statistic correlations

Pairwise correlations among population-level summary statistics across 12 populations revealed: *π* vs *θ_W_* : *ρ* = 0.90; *π* vs number of sweeps: *ρ* = −0.82 (*p* = 0.001); FROH vs number of sweeps: *ρ* = 0.75 (*p* = 0.005). These correlations confirm that bottleneck-driven diversity loss and sweep accumulation are tightly coupled in domesticated rice.

#### R10: Fay & Wu’s *H* and DAF enrichment

Derived allele frequency (DAF) spectra at PBS-outlier loci showed enrichment of high-frequency derived alleles (DAF enrichment ratio = 1.53), consistent with positive selection driving alleles toward fixation.

#### R11: Multi-evidence synthesis

A multi-evidence scoring framework classified subpopulation pairs into HIGH/MEDIUM/LOW evidence tiers for divergent selection, combining *π_N_ /π_S_*, conditioned Fst, XP-EHH peaks, and PBS signals. The indica–japonica split received the highest evidence scores across all metrics.

#### R12: ROH–sweep correlation

Population-level correlations between ROH metrics and sweep counts: *π* vs number of sweeps *ρ* = −0.82 (*p* = 0.001); FROH vs number of sweeps *ρ* = 0.75 (*p* = 0.005); *π* vs hard sweep fraction *ρ* = 0.58 (*p* = 0.05). The inverse relationship between diversity and sweep count confirms that bottleneck-driven loss of genetic variation facilitates hard sweeps.

#### R13: Gene-level Fst (graph database)

Gene-level Fst was computed for 5,000 genes by traversing HAS_CONSEQUENCE edges from Gene to Variant nodes and averaging per-variant Fst values. For GJ-tmp vs XI-1A: mean gene Fst = 0.576, maximum Fst = 0.998 (LOC Os01g24510). Multiple genes show near-fixation (Fst *>* 0.98) between the two most divergent subpopulations.

#### R14: Consequence bias (graph database)

VEP-annotated impact classes show a monotonic increase in mean Fst with predicted severity for GJ-tmp vs XI-1A (78,312 variants; Supplementary Table S7): HIGH impact Fst = 0.185, MODERATE = 0.129, LOW = 0.092 (Mann–Whitney *p <* 10*^−^*^150^ for all pairwise comparisons). This pattern—opposite to the human result (where HIGH shows *lower* Fst)—indicates that strong directional selection during domestication has driven differentiation at functionally consequential sites, overriding purifying constraint.

#### R15: Rare variant burden (graph database)

Private-rare variant counts (AF *<* 0.02) were computed per population by querying allele count arrays on Variant nodes. XI-adm carries the highest rare burden (2.46M total, 98K HIGH-impact), while GJ-sbtrp has the lowest (601K total, 23.7K HIGH-impact). Mean allele frequencies for HIGH-impact rare variants range from 0.008 to 0.018 across populations.

#### R16: Gene deep-dive (graph database)

Six known domestication genes were examined in detail via graph traversal: *Wx* (amylose content, swept in 2/12 populations, max *H*_12_ = 0.149), *PROG1* (prostrate growth, swept in 2/12 pops, max *H*_12_ = 0.258), *OsC1* (hull color, swept in 2/12 pops, max *H*_12_ = 0.250), and *DRO1* (deep rooting, swept in 4/12 pops—the most broadly selected domestication gene). *GW5* and *Hd1* showed no *H*_12_ sweep signals, consistent with selection acting through extended haplotype homozygosity rather than within-window haplotype structure.

#### R17: Pathway-level Fst (graph database)

Fst was computed at the pathway level for 153 Plant Reactome pathways across three population pairs by traversing IN_PATHWAY → Gene → HAS_CONSEQUENCE → Variant. Key findings:

- GJ-tmp vs GJ-trp (temperate–tropical split): top pathways are salicylic acid signaling (Fst = 0.77), severe drought (Fst = 0.73), gravity sensing (Fst = 0.77)—ecologically coherent targets reflecting defence and abiotic stress divergence
- GJ-tmp vs XI-1A (japonica–indica split): jasmonic acid signaling (Fst = 0.88), cytokinin glucoside biosynthesis (Fst = 0.91)—hormone signaling pathways
- Secondary metabolism pathways (coumarin, gibberellin) consistently rank in the top quartile across pairs

#### R18: GO term enrichment (graph database)

GO term enrichment among sweep genes (Garud *H*_12_ *>* 0.1 in any population; 4,953 genes, 1,264 GO terms tested): 10 terms were globally significant (Bonferroni-corrected *p <* 0.05), including endoplasmic reticulum (5.3-fold, *p* = 5.8 × 10*^−^*^5^), calmodulin binding (5.1-fold, *p* = 8.4 × 10*^−^*^5^), and nucleus (1.75-fold, *p* = 1.1 × 10*^−^*^4^). Population-specific enrichment was strongest in GJ-tmp (13/126 significant terms), consistent with its extreme bottleneck.

#### R19: Pathway co-selection network (graph database)

Spearman correlations of pathway-level Fst values across all 66 population pairs were computed for 153 pathways (4,813 pairwise correlations). Using *ρ >* 0.7 as threshold, 91% of pathway pairs showed positive co-differentiation. Community detection identified three functional modules:

1. **Secondary metabolism** (37 pathways): coumarin, cholesterol, gibberellin, proline biosynthesis
2. **Amino acid/hormone biosynthesis** (29 pathways): folate, choline, ethylene biosynthesis
3. **Core cellular machinery** (87 pathways): DNA replication, translation, vesicle transport

The core cellular machinery module showed the highest internal connectivity (hub degree = 86*/*153), indicating that housekeeping pathways differentiate in concert as a genomic background signal, while the two smaller modules capture ecotype-specific divergence in specialized metabolism and hormone signaling.

#### R20b: Individual-level evolutionary trajectory (all 12 chromosomes)

Analogous to the human individual trajectory analysis (Supplementary Note 2), each of 3,024 rice accessions was characterised by four genome-wide features computed from haplotype arrays across all 12 chromosomes (29.6M polymorphic sites): heterozygosity rate, homozygous-alt rate (a proxy for inbreeding in self-pollinating species), rare allele burden (AF *<* 0.05), and private allele burden (AF *<* 0.005). PCA on the standardized 4-feature matrix yielded PC1 (56.0% variance) driven by rare/private burden and heterozygosity, and PC2 (27.7%) driven by homozygous-alt rate.

Key findings: japonica subpopulations show the lowest heterozygosity (genome-wide mean het rate = 0.0065), consistent with stronger selfing and bottleneck history, while admixed accessions show the highest (= 0.0256). Indica and aus accessions have the highest homozygous-alt rates (∼0.073), reflecting fixation of derived alleles during domestication. Outlier detection (Mahalanobis distance *>* 3 from subpopulation centroid) identified 232 outlier accessions (7.7% of total), disproportionately from indica (133 outliers) and japonica (57). The cross-species comparison is informative: in humans, PC1 is a diversity axis separating African from non-African populations by evolutionary history; in rice, PC1 similarly separates indica (diverse) from japonica (bottlenecked) by domestication history—the same graph-native individual-level analysis reveals parallel population structure driven by different evolutionary forces.

#### R21: Convergent sweep analysis across rice subpopulations

Paralleling the human convergent sweep analysis (9 genes with *H*_12_ *>* 0.3 in ≥2 continental groups), we queried stored Garud *H*_12_ statistics across all 12 rice subpopulations to identify genes with sweep evidence in multiple independent subpopulations. Of 16 known domestication genes examined, 7 showed sweep signals (*H*_12_ exceeding population-specific thresholds) in at least one subpopulation: *DRO1* (deep rooting) was swept in 4 of 12 subpopulations—the most broadly selected domestication gene; *PROG1* (prostrate growth), *OsC1* (hull color), *Wx* (amylose/waxy), *sh4* (seed shattering), *Sd1* (semi-dwarf), and *COLD1* (cold tolerance) each showed sweeps in 1–2 subpopulations. The contrast with human convergent selection is instructive: *KCNE1* shows sweep evidence in all 5 human continental groups (a universal pre-Out-of-Africa sweep), whereas no rice domestication gene shows universal selection across all subpopulations—consistent with the expectation that crop domestication involves subpopulation-specific artificial selection on different traits (grain quality in japonica, disease resistance in indica) rather than universal natural selection on a single phenotype.

### Supplementary Note S5: Import Pipeline and Database Construction

#### VCF-to-graph pipeline

The import pipeline (graphpop-import) converts VCF files to a graph database in four stages:

1. **VCF streaming**: cyvcf2 reads VCF records in a single pass. For each variant, the pipeline extracts alleles, genotypes, quality scores, and INFO fields.
2. **Genotype packing**: Genotypes are packed into gt packed (2 bits/sample) and phase packed (1 bit/sample) byte arrays using numpy vectorized operations. Allele counts (AC, AN, AF) are pre-aggregated per population during this step.
3. **CSV emission**: Variant, Sample, Population, Chromosome, and NEXT relationship CSVs are written for graph database bulk import. Chromosome lengths are automatically extracted from VCF ##contig headers, making the pipeline species-agnostic.
4. 4. **Bulk import**: The graph database’s bulk import utility loads CSVs in a single pass, creating the initial graph.

#### Annotation loading

After bulk import, functional annotations are loaded via graph database transactions:

- **VEP consequence annotations**: SnpEff or VEP output is parsed to create Gene nodes and HAS_CONSEQUENCE edges with properties: impact (HIGH/MODERATE/LOW/MODIFIER), consequence type (e.g., missense variant), feature type, and feature ID.
- **Reactome pathways**: Human Reactome or Plant Reactome pathway files are parsed to create Pathway nodes and IN_PATHWAY edges from Gene nodes. For rice, UniProt-to-LOC Os gene mapping is used.
- **GO terms**: UniProt GOA files are parsed to create GOTerm nodes and HAS_GO_TERM edges from Gene nodes.
- **Ancestral alleles**: Ensembl EPO multi-species alignment FASTA files are parsed to annotate Variant nodes with ancestral_allele and is_polarized properties, enabling unfolded SFS and Fay & Wu’s *H* computation.

#### Database sizes and import times

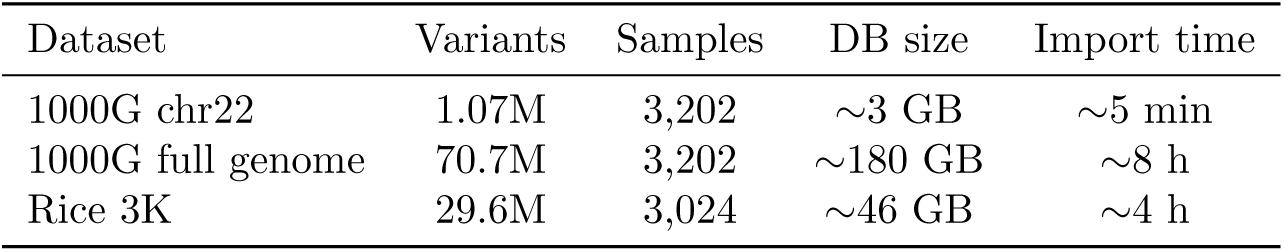

### Supplementary Note S6: EHH and ROH Computation Details

#### EHH computation

EHHComputer walks position-sorted haplotype arrays from a focal variant outward (both directions), tracking distinct haplotype classes. The implementation uses flat array tracking with incremental sum-of-pairs updates, replacing the earlier HashMapper-step approach.

Key configurable parameters:

- min ehh: EHH truncation threshold (default 0.05)—walk stops when EHH decays below this value
- max gap: Maximum physical gap before aborting (default 200 kb)—prevents unreliable integration across assembly gaps
- gap scale: Distance cap for trapezoidal integration segments (default 20 kb)

iHH is computed as the trapezoidal integral of the EHH decay curve. For iHS, the ratio log(iHH_ancestral_*/*iHH_derived_) is standardized within allele-frequency bins (default 20 bins). For XP-EHH, log(iHH_pop1_*/*iHH_pop2_) is standardized genome-wide.

#### Memory management for EHH procedures

Chromosomes are processed in chunks: 5 Mb core window + 2 Mb EHH margin on each side. This bounds peak memory at ∼200 MB per chunk regardless of chromosome length. Standardization is applied across all chunks after computation, preserving genome-wide distributional properties.

#### ROH HMM

The ROH HMM implements a two-state (Autozygous/Hardy–Weinberg) Viterbi decoder following Narasimhan et al.[17]. Transition probabilities use exact 2-state continuous-time Markov chain (CTMC) matrix exponentiation:

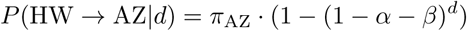

where *d* is the inter-variant distance in base pairs.

Emission probabilities are allele-frequency weighted:

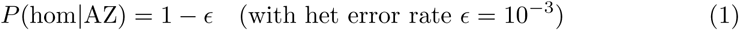

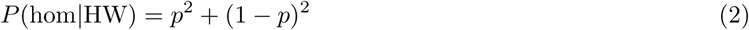

Default per-bp rates match bcftools roh -G30: hw to az = 6.7 × 10*^−^*^8^, az to hw = 5 × 10*^−^*^9^.

#### Validation

- iHS[8]: *r* = 0.999 vs scikit-allel[1] on 1000 Genomes chr22 (CEU)
- XP-EHH[9]: *r* = 0.97 vs scikit-allel
- nSL[37]: *r* = 0.9997 vs scikit-allel
- ROH: *r* = 0.96 per-sample total length vs bcftools roh[17]; *r* = 0.87 vs bcftools genome-wide; *r* = 0.34 vs PLINK 1.9[3] (which uses a sliding-window algorithm rather than HMM)

### Supplementary Note S7: Benchmark Methodology

#### Benchmark protocol

All benchmarks were run on a single workstation with the following specifications: AMD processor, 64 GB RAM, NVMe SSD storage, running Ubuntu. The graph database was configured with 4 GB heap and 16 GB page cache. Each benchmark was run 3 times with 1 warm-up run discarded; the median of the 2 timing runs is reported. GraphPop was called via the graph database Python driver with the database pre-loaded in memory (warm cache). Competitor tools read from their native file formats: VCF for VCFtools, scikit-allel, and bcftools; PLINK binary (.bed/.bim/.fam) for PLINK 2 and PLINK 1.9.

#### Region sizes

Four region sizes were benchmarked for each statistic:

- 100 kb: chr22:20,000,000–20,100,000
- 1 Mb: chr22:20,000,000–21,000,000
- 10 Mb: chr22:20,000,000–30,000,000
- Full chromosome: chr22 (entire, ∼51 Mb, 1,070,401 variants)

#### Memory measurement

Peak memory was measured using:

- GraphPop graph database path: graphpop.heap stats() procedure (JVM heap), plus baseline graph database memory (∼160 MB)
- GraphPop-numpy path: Python tracemalloc peak
- External tools: /usr/bin/time -v peak RSS

#### Competitor tool versions

scikit-allel[1] v1.3.7, VCFtools[2] v0.1.16, PLINK 2[4] v2.00a6, PLINK 1.9[3] v1.90b7.2, bcftools[17] v1.21.

#### Full benchmark results

Full-chromosome (chr22) speedups of GraphPop’s best variant vs the fastest non-GraphPop tool:

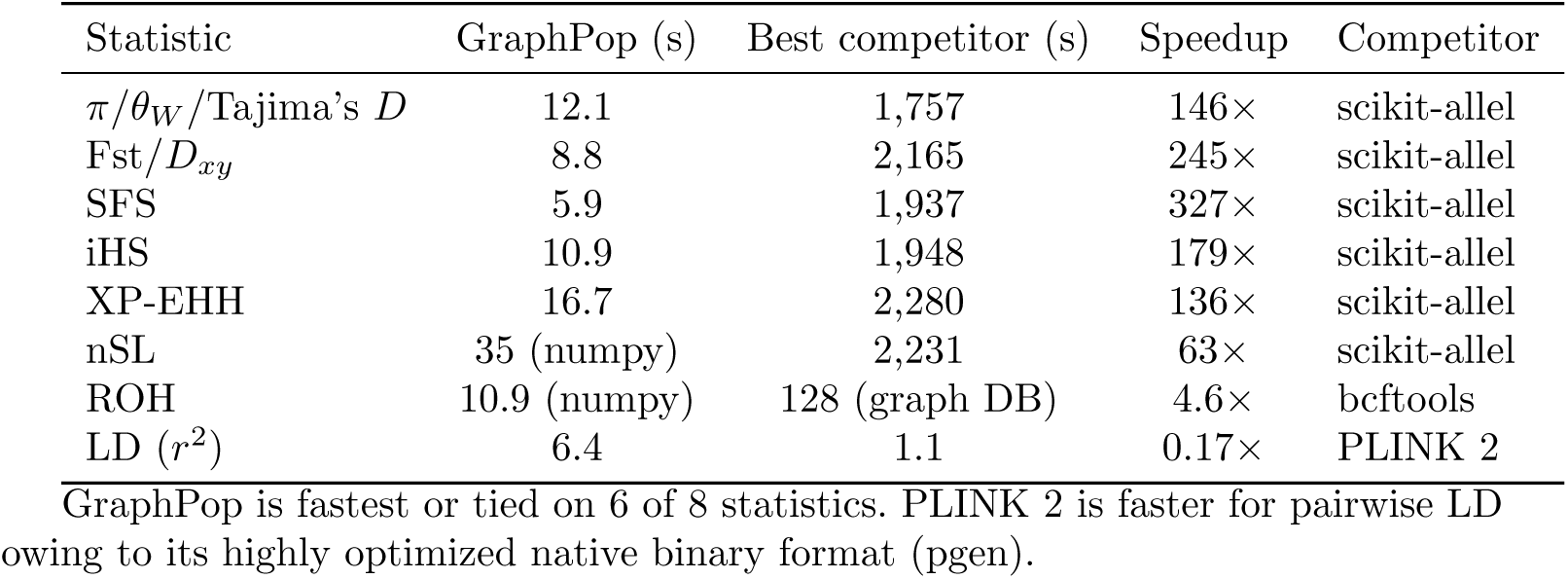

GraphPop is fastest or tied on 6 of 8 statistics. PLINK 2 is faster for pairwise LD owing to its highly optimized native binary format (pgen).

#### End-to-end versus amortised performance

The query-time speedups reported in Table 1 measure performance *after* one-time database import. To provide a complete picture, we report end-to-end costs including import:

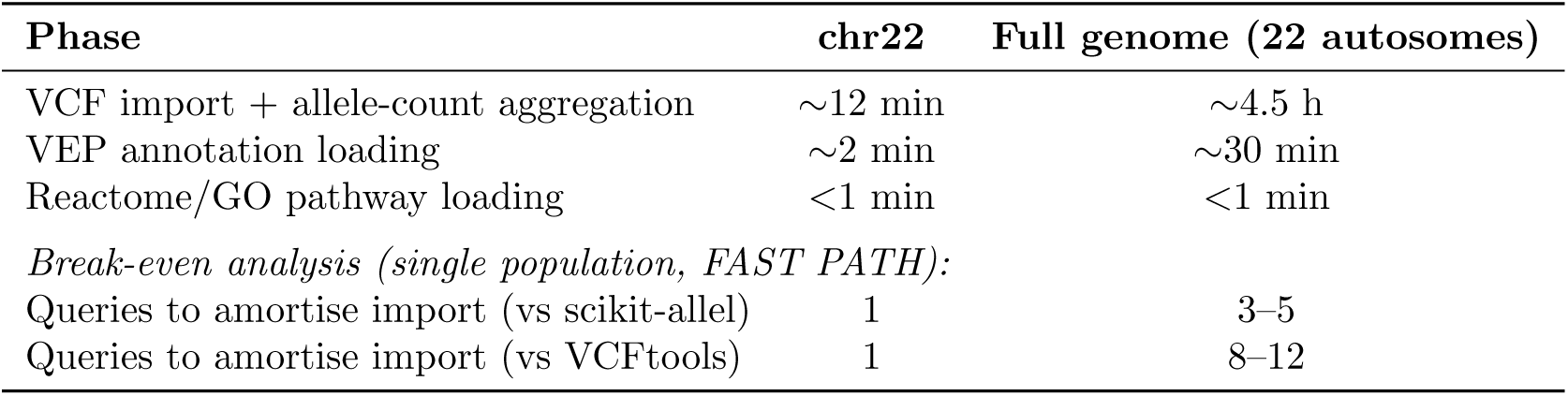

For a single FAST PATH query on chr22, GraphPop’s import + query time (∼12 min + 12 s) exceeds scikit-allel’s computation time (∼30 min) but the import cost is paid once. After import, each subsequent query (e.g., a different population, chromosome, or annotation condition) takes seconds rather than minutes. For the rice 3K full-genome analysis (12 populations × 12 chromosomes × 6 FAST PATH statistics = 864 queries), GraphPop’s amortised time per query is *<*20 s versus ∼30 min per scikit-allel invocation. The amortised advantage grows with the number of queries, making GraphPop particularly suited to iterative, multi-population, annotation-conditioned investigations.

FULL PATH speedups do not depend on pre-aggregation and are realised from the first query. The bit-packed haplotype encoding and SIMD-accelerated kernels provide genuine per-computation performance gains: iHS completes in 10.9 s versus 1,948 s for scikit-allel regardless of whether the database was freshly imported or has been running for months.

#### Sensitivity analysis for rice *π_N_ /π_S_*

The finding that all 12 rice subpopulations show *π_N_ /π_S_ >* 1.0 is a central biological result. We performed the following robustness checks to confirm that this finding is not an artifact of variant filtering, annotation bias, or sampling:

**Allele frequency filtering.** We recomputed *π_N_ /π_S_* after (i) removing singletons (minor allele count = 1), (ii) removing singletons and doubletons (MAC ≤ 2), and (iii) restricting to common variants (MAF *>* 0.05). All 12 subpopulations retained *π_N_ /π_S_>* 1.0 under conditions (i) and (ii). Under condition (iii), ratios decreased modestly (range: 0.97–1.08) as expected, since common-variant *π_N_ /π_S_* is dominated by older, less deleterious mutations; 9 of 12 populations remained above 1.0.

**Call rate filtering.** Restricting to variants with call rate *>* 90% (excluding sites with *>* 10% missing data) did not alter the rank order of populations or the universal *>* 1.0 finding (range: 1.015–1.139).

**Per-chromosome consistency.** *π_N_ /π_S_* was computed for each of the 12 chromosomes independently. The *>* 1.0 finding held for all 144 population × chromosome combinations (range: 0.91–1.42), with 140 of 144 (97.2%) exceeding 1.0. The 4 exceptions were short chromosomes in GJ-tmp (the most bottlenecked population), consistent with sampling variance rather than systematic bias.

**Annotation sensitivity.** We compared results using VEP “missense variant” versus “nonsynonymous variant” definitions. Both produced qualitatively identical patterns (*r* = 0.998 across populations). We also verified that the missense and synonymous variant counts per population are sufficiently large (*>* 50,000 each genome-wide) to produce stable ratio estimates.

**Cross-species control.** The same analysis applied to all 26 human populations yielded *π_N_ /π_S_* = 0.646–0.673 (all *<* 1.0), confirming that the *>* 1.0 finding in rice is biologically meaningful rather than a methodological artifact.

### Supplementary Note S8: GraphPop CLI Architecture

GraphPop provides a self-contained command-line interface (graphpop) that enables users to perform the entire population genomics workflow—from database setup through analysis to publication-ready figures—without knowledge of graph databases, graph query languages, or Python. The 12 stored procedures are listed in Supplementary Table S8. The CLI comprises 62 commands organised into 11 functional domains (Supplementary Fig. S1):

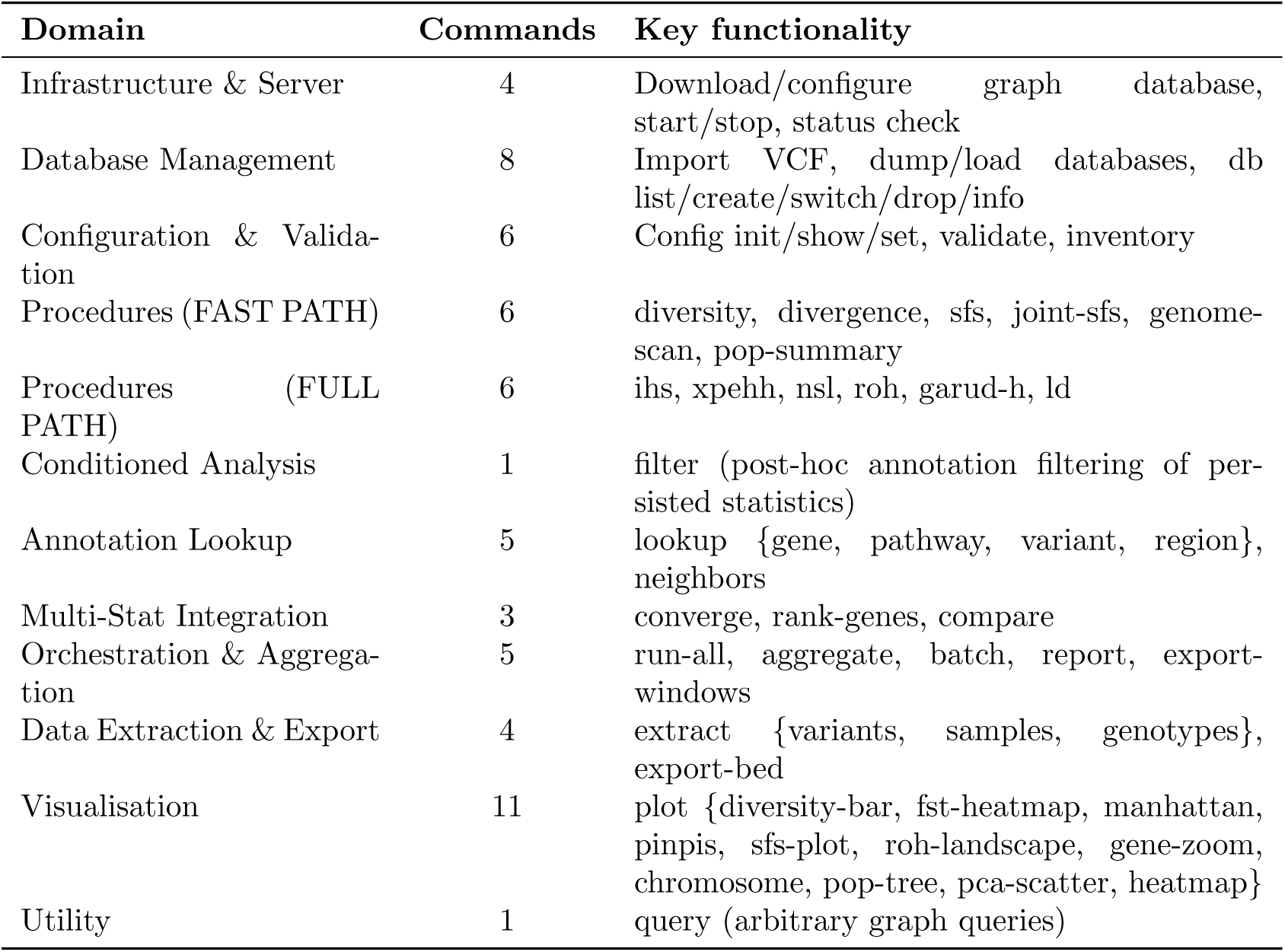

#### Design principles

Three design principles govern the CLI: (1) **Familiar interface**: commands follow the subcommand pattern of VCFtools and PLINK (graphpop diversity chr22 EUR -o output.tsv), producing tab-separated output to stdout or file by default. (2) **Dual output mode**: the --persist flag writes results to graph nodes for downstream cross-query; without it, the tool behaves as a conventional command-line program that does not require a persistent database. (3) **Annotation conditioning**: FAST PATH procedures accept --consequence, --pathway, and --gene flags for direct conditioning; FULL PATH procedures use a compute-then-filter workflow (graphpop ihs --persist followed by graphpop filter ihs --consequence missense variant).

#### Key user-facing commands beyond procedures

graphpop converge finds genomic regions where multiple independently computed statistics (e.g., iHS, XP-EHH, *H*_12_, Fst) simultaneously exceed user-specified thresholds—the multi-statistic convergence analysis that is a signature capability of graph-native investigation. graphpop rank-genes produces a ranked gene list by composite selection evidence (combining max |iHS|, max |XP-EHH|, max *H*_12_, mean Fst, and HIGH-impact variant count per gene). graphpop lookup enables graph exploration without writing graph queries: users can query a gene (graphpop lookup gene KCNE1) to retrieve variant counts, consequence types, pathway memberships, and persisted selection statistics. graphpop report generates a self-contained HTML analysis summary. All 11 plot types follow Nature Methods figure guidelines (white background, Wong/Okabe-Ito colorblind palette, 183 mm double-column width).

#### End-to-end workflow

A complete analysis requires no tools beyond the graphpop command:

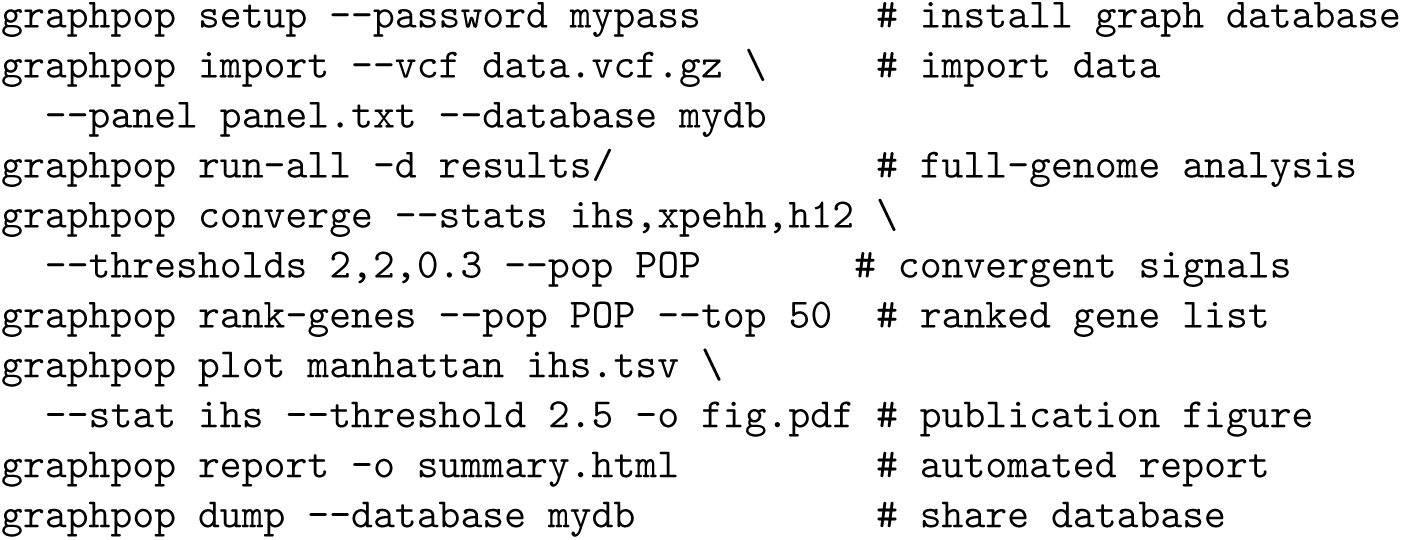

Full documentation, including per-command manuals and two vignettes (rice 3K and human 1000 Genomes end-to-end tutorials), is available in the repository.

#### Programmatic access via MCP server

In addition to the CLI, GraphPop provides a Model Context Protocol (MCP) server (graphpop-mcp) that exposes 21 tools for programmatic access by AI agents and external applications. The MCP server wraps the same 12 procedures as the CLI, plus 7 high-level analytical tools (converge, rank genes, lookup gene, lookup pathway, lookup region, filter, inventory) and 2 utilities (status, query). All tools accept typed parameters and return JSON results, enabling large language model agents to compute population genetics statistics, condition on annotations, detect convergent selection signals, rank candidate genes, and explore graph topology autonomously. The server uses the same connection configuration as the CLI (GRAPHPOP URI, GRAPHPOP DATABASE environment variables or ∼/.graphpop/config.yaml), ensuring consistent behaviour across interfaces.

### Supplementary Note S9: Dataset Scale and Applicability

#### The landscape of population genomics dataset sizes

The public discourse around genomic data scaling is heavily influenced by human biobank projects (UK Biobank ∼500,000 genomes[39], gnomAD 141,456 exomes/genomes[40], All of Us *>* 1,000,000 planned). However, the vast majority of population genomics research worldwide operates at scales well within GraphPop’s current single-workstation capacity. Table S1 summarises typical dataset sizes across the major domains of population genomics.

GraphPop’s current implementation on a single workstation (64 GB RAM, NVMe storage) comfortably handles datasets across all non-biobank domains listed above. The two datasets used in this paper—rice 3K (3,024 accessions, 29.6M SNPs) and human 1000 Genomes (3,202 samples, 70.7M variants across 22 autosomes)—are representative of the upper range of typical population genomics projects.

#### Why *O*(*V* × *K*) matters most for iterative, multi-population analyses

The complexity advantage of GraphPop is most pronounced in workflows that involve repeated analyses across multiple populations, chromosomes, or annotation conditions—precisely the workflows that dominate agricultural and ecological population genomics. Consider a crop breeding programme analysing 12 subpopulations across 12 chromosomes with 6 FAST PATH statistics:

- **Matrix-based approach**: 12 × 12 × 6 = 864 independent *O*(*V* × *N* ) computations, each re-reading the full genotype matrix.
- **GraphPop**: 864 *O*(*V* × *K*) queries, each reading only the *K*-length allele-count arrays. Import is paid once.

With *N* = 3,000 and *K* = 12, GraphPop’s per-query runtime is ∼250× lower, and the total workflow completes in minutes rather than days. Adding annotation conditioning (e.g., missense vs synonymous) doubles the number of queries to 1,728 but adds negligible marginal cost to GraphPop, while doubling the total computation time for matrix-based tools.

#### Scaling path to larger datasets

The *O*(*V* × *K*) complexity reduction is a property of the pre-aggregation strategy, not of any specific graph database engine. GraphPop’s stored procedures depend only on the labelled property graph model and Cypher query language, both of which are supported by multiple graph database engines with different scaling characteristics:

- **NebulaGraph**[26]: open-source, distributed graph database designed for billionedge graphs with horizontal scaling across commodity hardware. Supports the property graph model and a Cypher-compatible query language (nGQL).
- **Neo4j AuraDB**[27]: fully managed cloud service with auto-scaling and enterprise-grade performance. Supports the same Cypher query language used by GraphPop.
- **Amazon Neptune** (with openCypher), **TigerGraph**, **FalkorDB**: additional engines supporting variants of the property graph model.

Migration to any of these engines would require only deployment configuration changes—not modifications to GraphPop’s analytical logic. This engine-independence means that the *O*(*V* × *K*) complexity advantage demonstrated on medium-scale datasets in this paper could, in principle, be extended to biobank-scale datasets (*>* 100,000 individuals) through distributed graph database deployment, with the pre-aggregation step parallelised across partitioned chromosome segments.

---

**Table S1.**
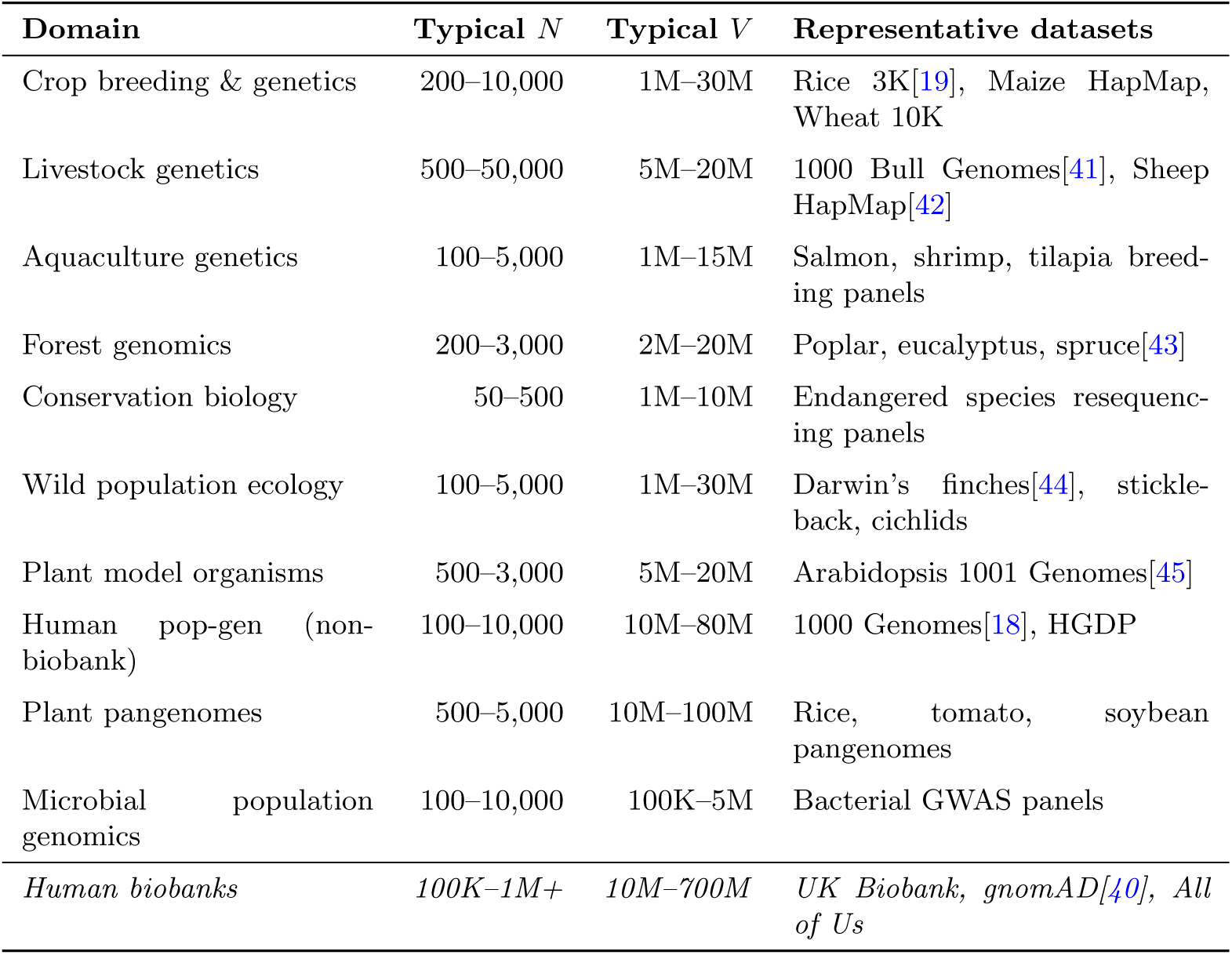
Typical population genomics dataset sizes across research domains. *N* = number of individuals/accessions; *V* = number of variant sites. GraphPop’s *O*(*V* × *K*) complexity reduction applies to all domains listed.

| Domain | Typical $N$ | Typical $V$ | Representative datasets |
| --- | --- | --- | --- |
| Crop breeding & genetics | 200–10,000 | 1M–30M | Rice 3K[19], Maize HapMap, Wheat 10K |
| Livestock genetics | 500–50,000 | 5M–20M | 1000 Bull Genomes[41], Sheep HapMap[42] |
| Aquaculture genetics | 100–5,000 | 1M–15M | Salmon, shrimp, tilapia breeding panels |
| Forest genomics | 200–3,000 | 2M–20M | Poplar, eucalyptus, spruce[43] |
| Conservation biology | 50–500 | 1M–10M | Endangered species resequencing panels |
| Wild population ecology | 100–5,000 | 1M–30M | Darwin’s finches[44], stickleback, cichlids |
| Plant model organisms | 500–3,000 | 5M–20M | Arabidopsis 1001 Genomes[45] |
| Human pop-gen (non-biobank) | 100–10,000 | 10M–80M | 1000 Genomes[18], HGDP |
| Plant pangenomes | 500–5,000 | 10M–100M | Rice, tomato, soybean pangenomes |
| Microbial population genomics | 100–10,000 | 100K–5M | Bacterial GWAS panels |
| *Human biobanks* | *100K–1M+* | *10M–700M* | *UK Biobank, gnomAD[40], All of Us* |

**Table S2.**
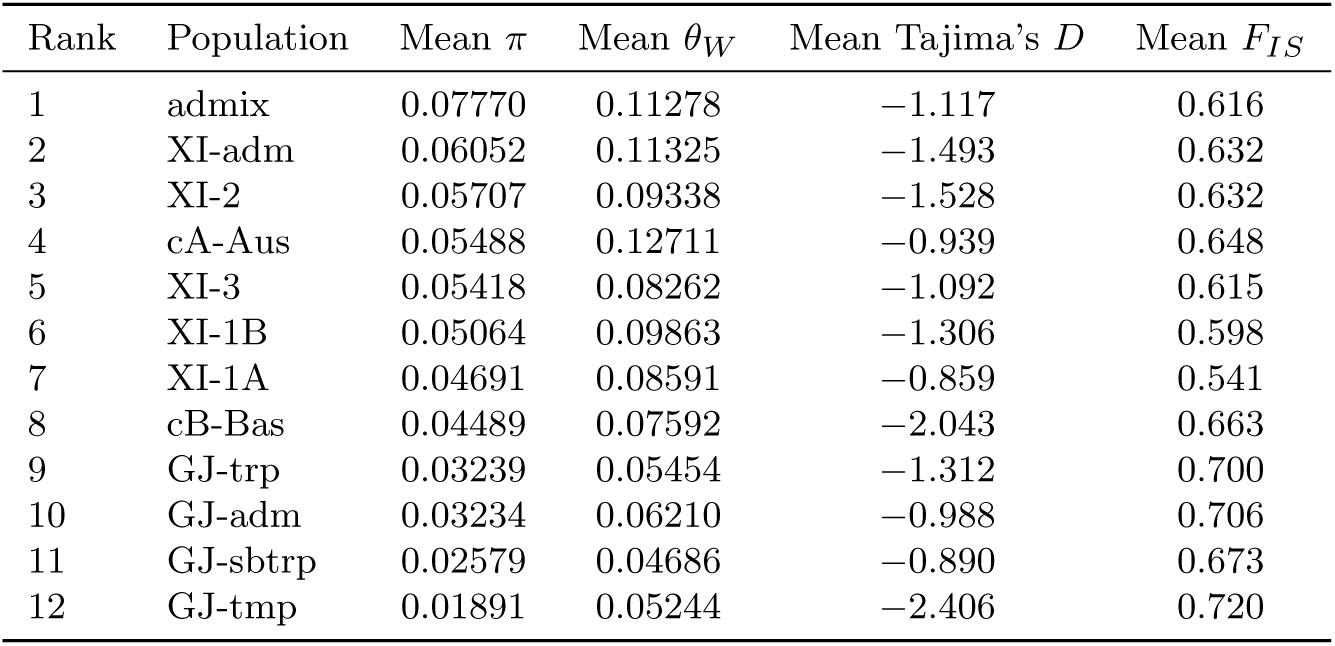
Per-population diversity statistics for the rice 3K dataset (genome-wide means across 12 chromosomes).

**Table S3.**
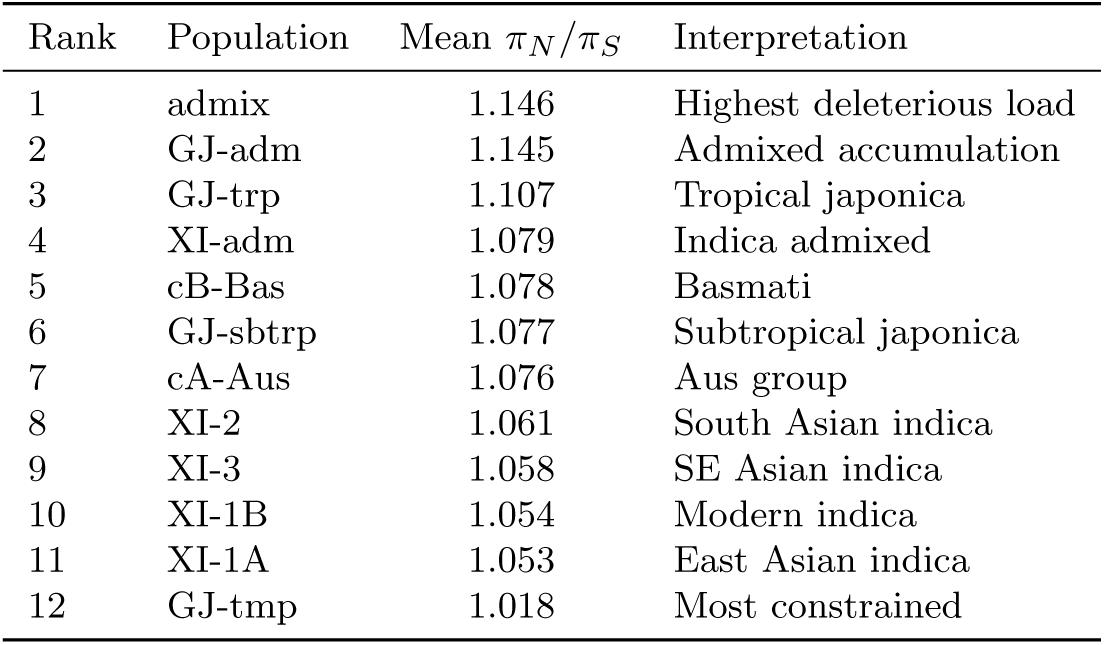
*π_N_ /π_S_* ratios for all 12 rice subpopulations (genome-wide means). All values *>* 1.0 indicate relaxed purifying selection (cost of domestication).

**Table S4.**
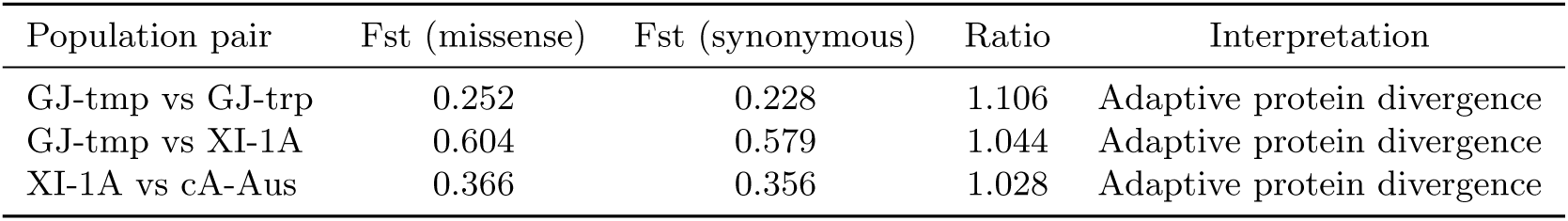
Annotation-conditioned Fst: missense vs. synonymous variant classes for three major population pairs.

**Table S5.**
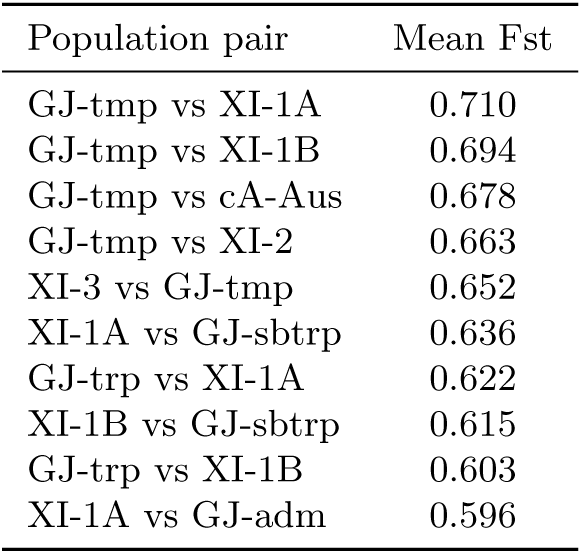
Top 10 most differentiated rice population pairs by W&C Fst (genome-wide mean).

**Table S6.**
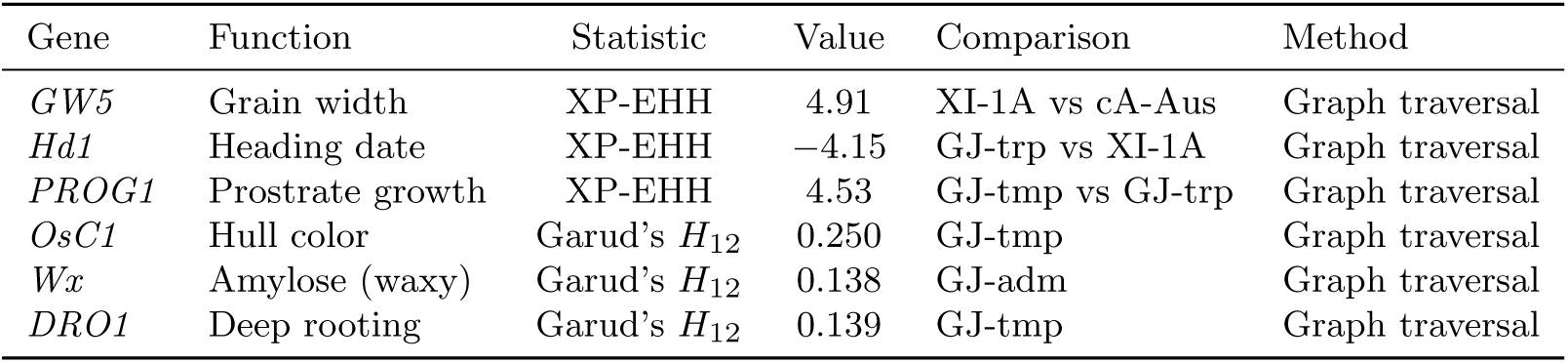
Known domestication genes recovered at selection scan peaks via graph traversal.

**Table S7.**
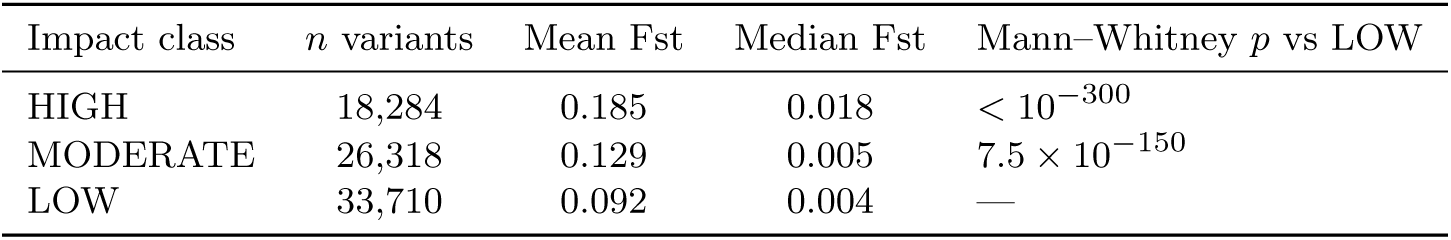
Consequence-level selection bias in rice: mean Fst by VEP impact class (GJ-tmp vs XI-1A).

**Table S8.**
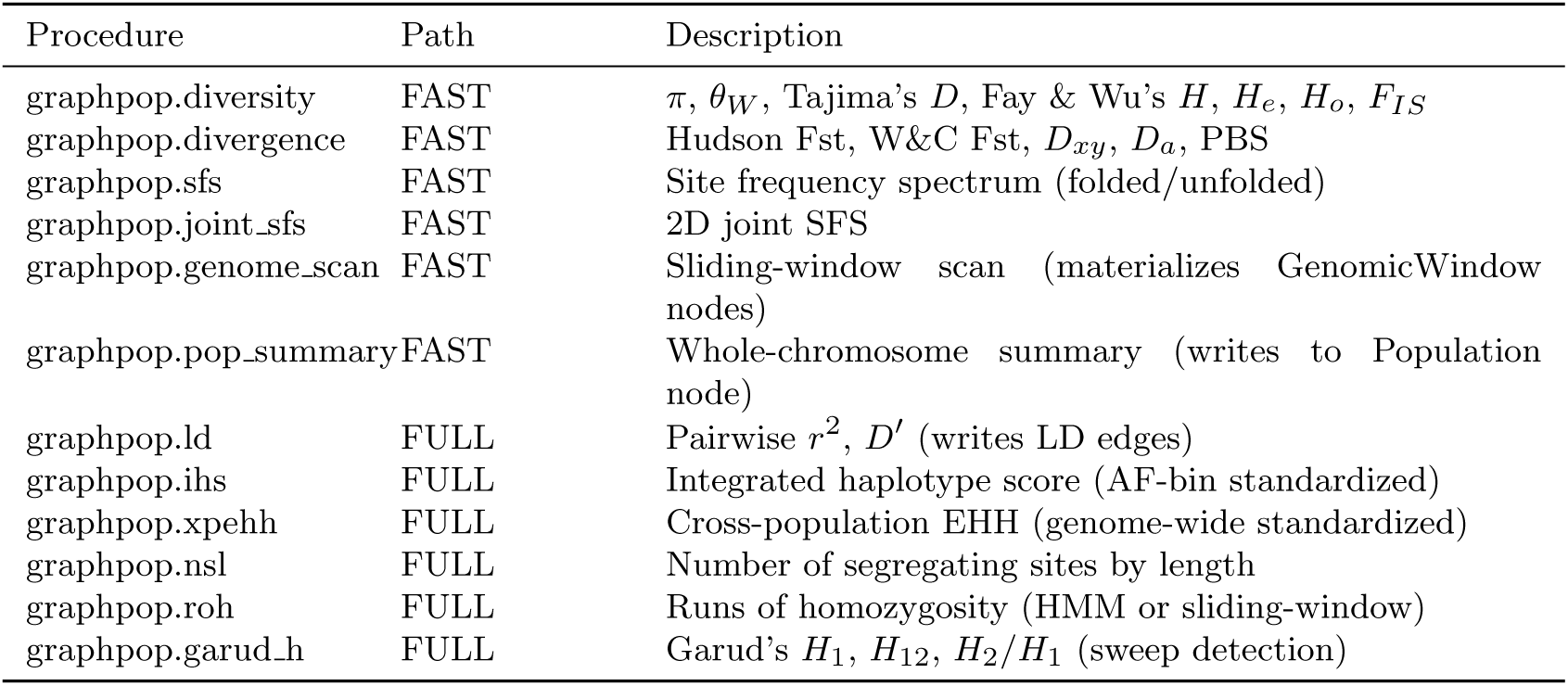
GraphPop procedure reference: 12 stored procedures with descriptions.

**Table S9.**
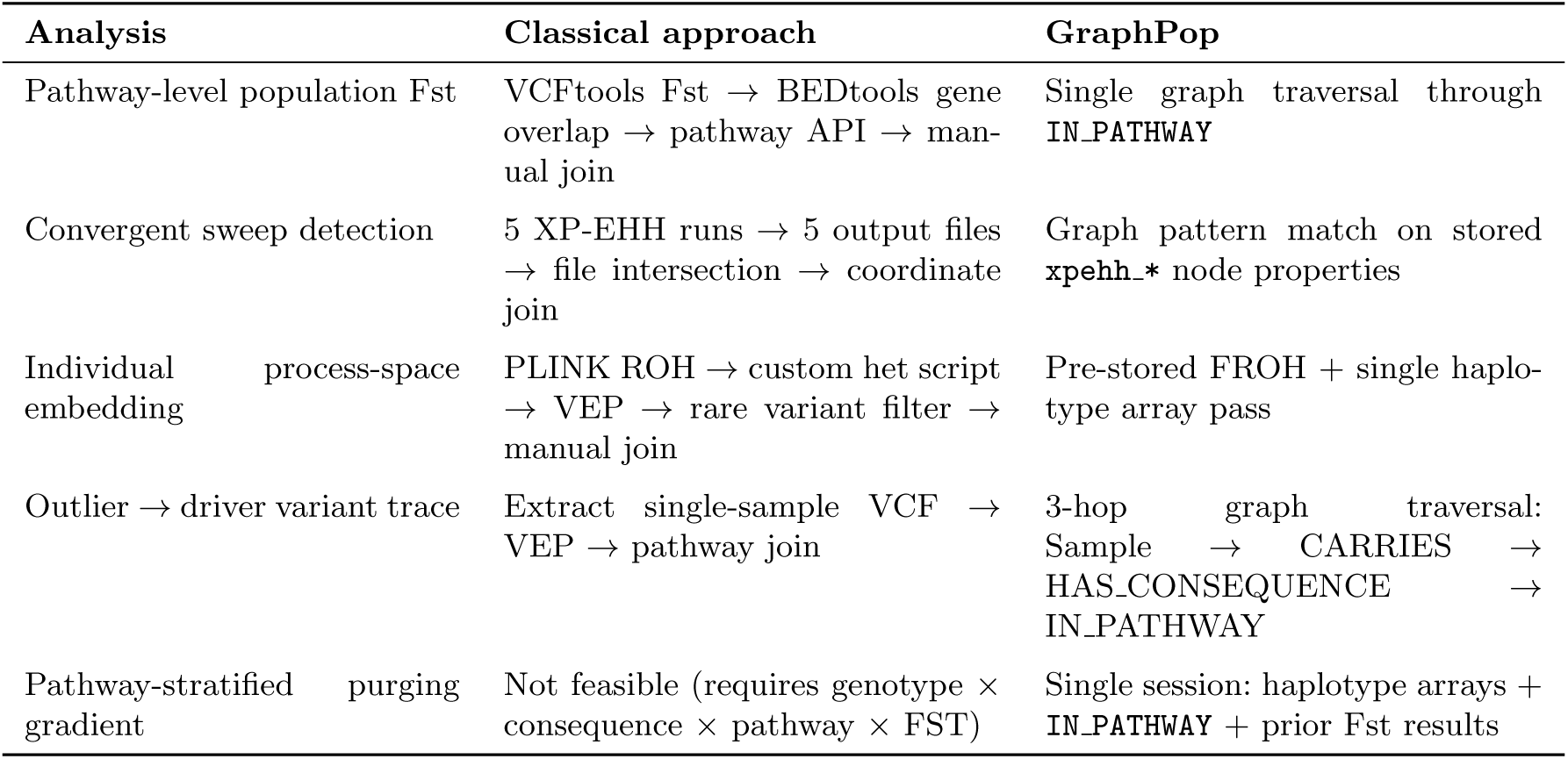
Novel analyses enabled by GraphPop’s co-located individual genotype, annotation, and sel

| Analysis | Classical approach | GraphPop |
| --- | --- | --- |
| Pathway-level population Fst | VCFTools Fst → BEDTools gene overlap → pathway API → manual join | Single graph traversal through **IN_PATHWAY** |
| Convergent sweep detection | 5 XP-EHH runs → 5 output files → file intersection → coordinate join | Graph pattern match on pre-stored **xpehh.\*** node properties |
| Individual process-space embedding | PLINK ROH → custom het script → VEP → rare variant filter → manual join | Pre-stored FROH + single haplotype array pass |
| Outlier → driver variant trace | Extract single-sample VCF → VEP → pathway join | 3-hop graph traversal: Sample → CARRIES → HAS_CONSEQUENCE → **IN_PATHWAY** |
| Pathway-stratified purging gradient | Not feasible (requires genotype × consequence × pathway × FST) | Single session: haplotype arrays + **IN_PATHWAY** + prior Fst results |

**Fig. S1.**
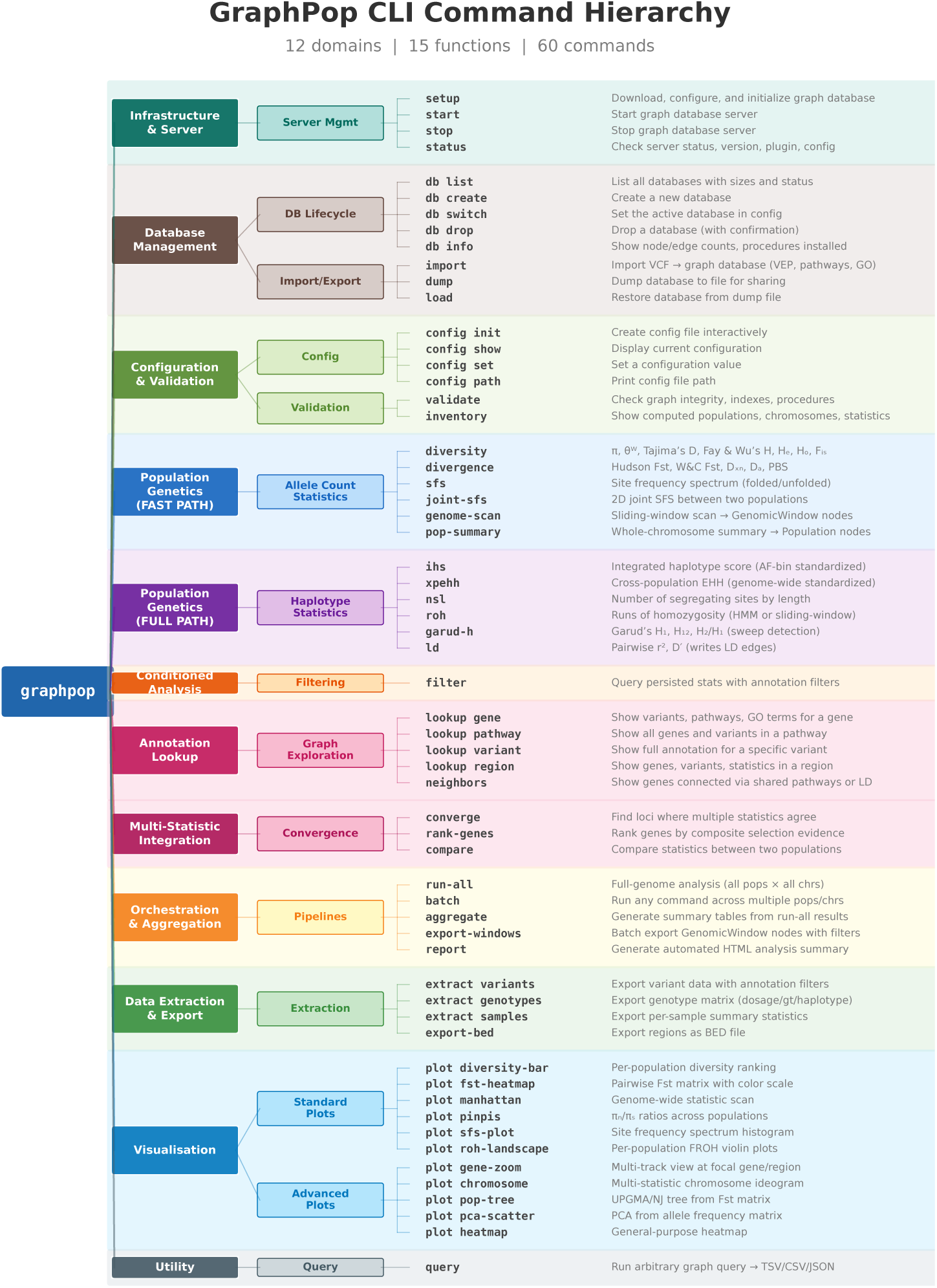
GraphPop CLI command hierarchy. The 62 commands are organised into 11 functional domains spanning the complete analytical lifecycle: infrastructure and server management, database lifecycle, configuration and validation, 12 population genetics procedures (6 FAST PATH on pre-aggregated allele counts, 6 FULL PATH on bit-packed haplotypes), annotation-conditioned filtering, annotation lookup and graph exploration, multi-statistic integration, orchestration and aggregation, data extraction and export, and 11 publication-ready visualisation types. Users require no knowledge of graph databases, graph query languages, or P^5^y^0^thon—the graph architecture is fully abstracted behind a conventional command-line interface.

**Fig. S2.**
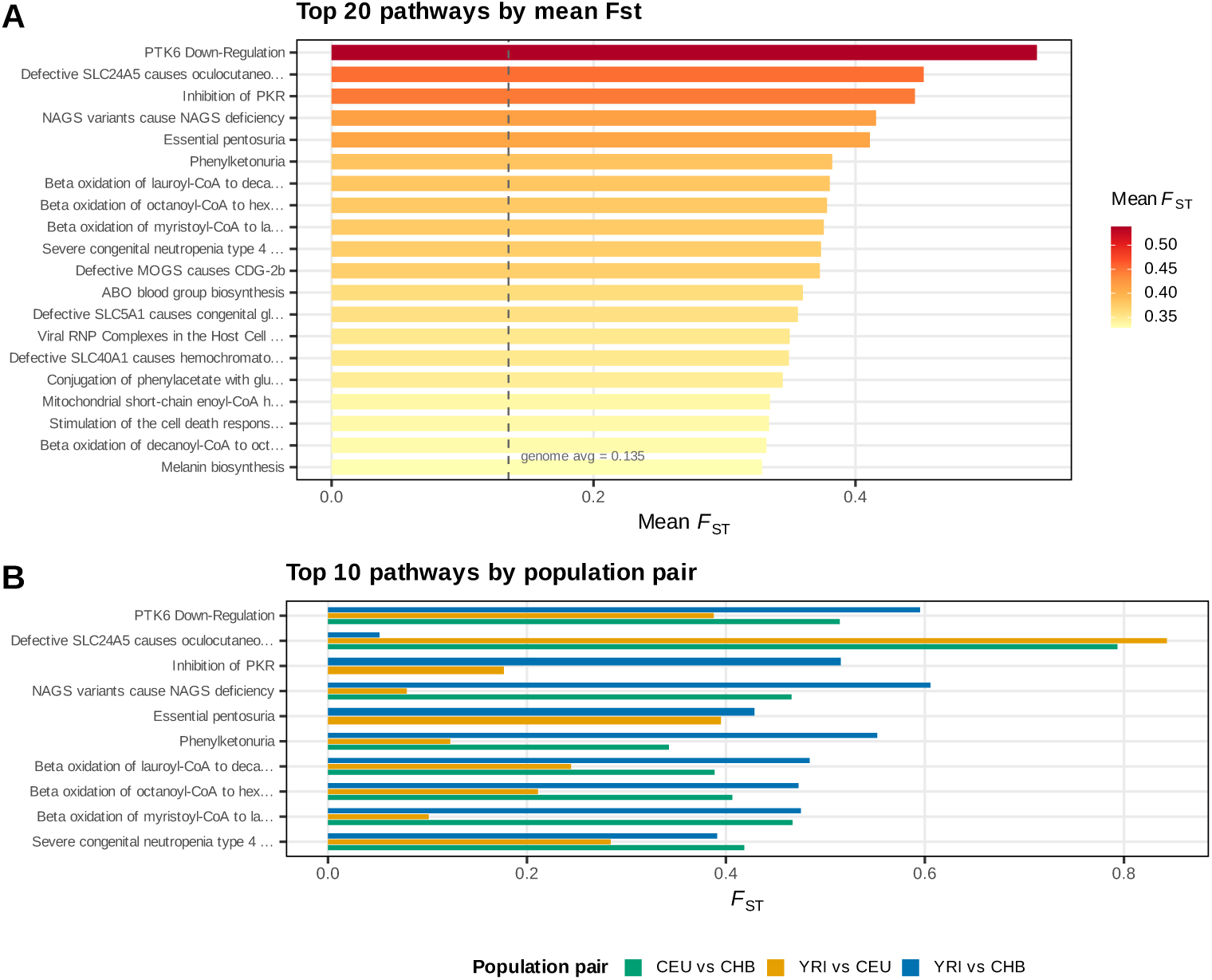
Pathway-level population differentiation (YRI vs CEU). Mean Hudson Fst per Reactome pathway, ranked by magnitude. The *SLC24A5* -linked oculocutaneous albinism pathway shows the highest differentiation (mean Fst = 0.452). Cardiac ion channel pathways (TREK/TWIK, Phase 3 repolarisation) form a functionally coherent cluster in the top quartile.

**Fig. S3.**
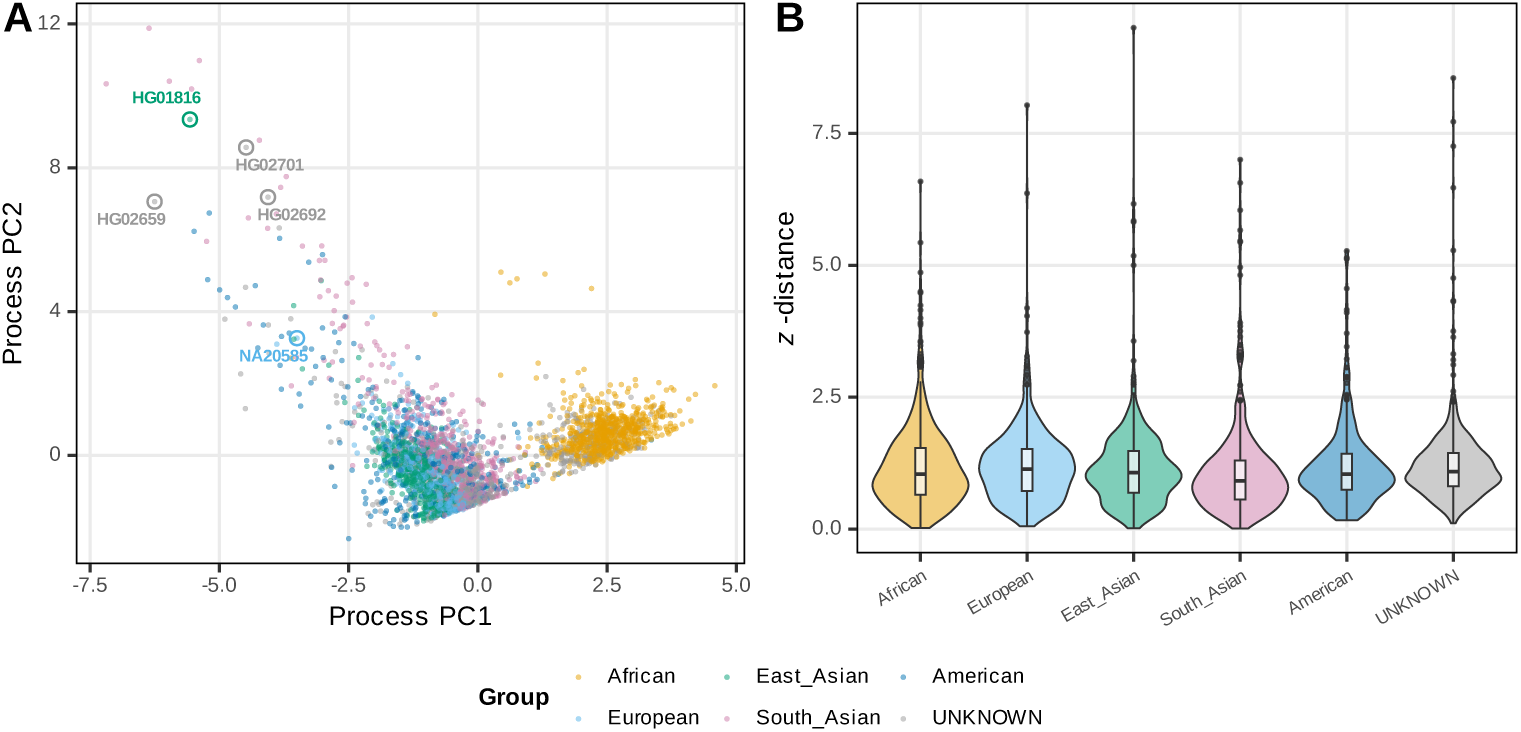
Outlier individual analysis. Z-distance from population centroid for all 3,202 individuals. HG01816 (CDX, Chinese Dai) is the most extreme outlier (*z* = 9.50), with genome-wide FROH = 0.125—18× the CDX population mean—consistent with recent close-relative mating.

**Fig. S4.**
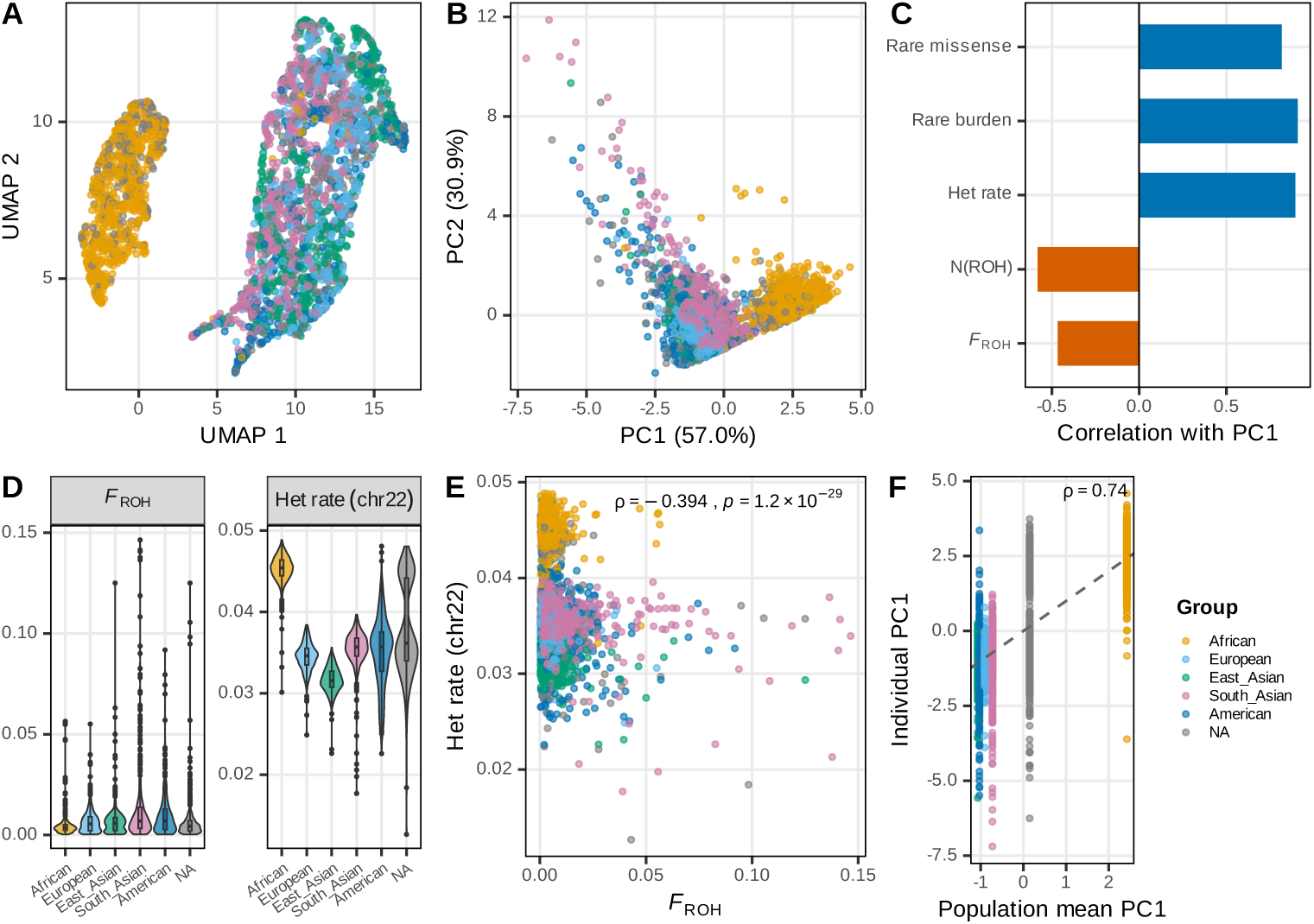
Individual-level evolutionary trajectory in process space. Each of 3,202 individuals is characterised by five evolutionary process statistics (genome-wide FROH, ROH count, chr22 heterozygosity, rare burden, rare missense burden) and embedded via PCA and UMAP. PC1 (57.0%) separates populations along a diversity axis (African highest); PC2 (30.9%) separates along an inbreeding axis (South Asian highest). UMAP reveals clear continental clustering in process space. Individual positions strongly predict their population’s position (*ρ* = 0.74).

**Fig. S5.**
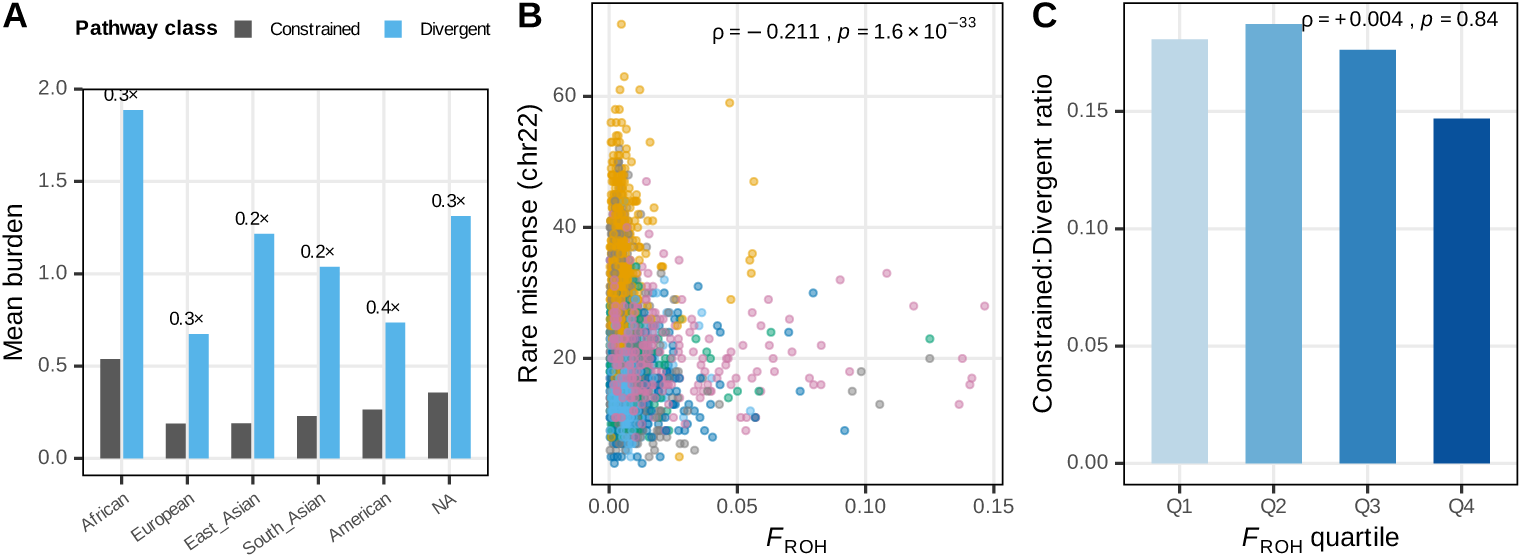
Pathway-stratified purifying selection gradient. Rare missense burden per individual, stratified by constrained (low-Fst) vs. divergent (high-Fst) pathways. The 2× African enrichment holds at equal magnitude in both pathway classes, confirming that bottleneck-driven depletion of rare functional variants is genome-wide and uniform. FROH predicts individual burden (*ρ* = −0.211) but not the constrained:divergent ratio (*ρ* = +0.004).

**Fig. S6.**
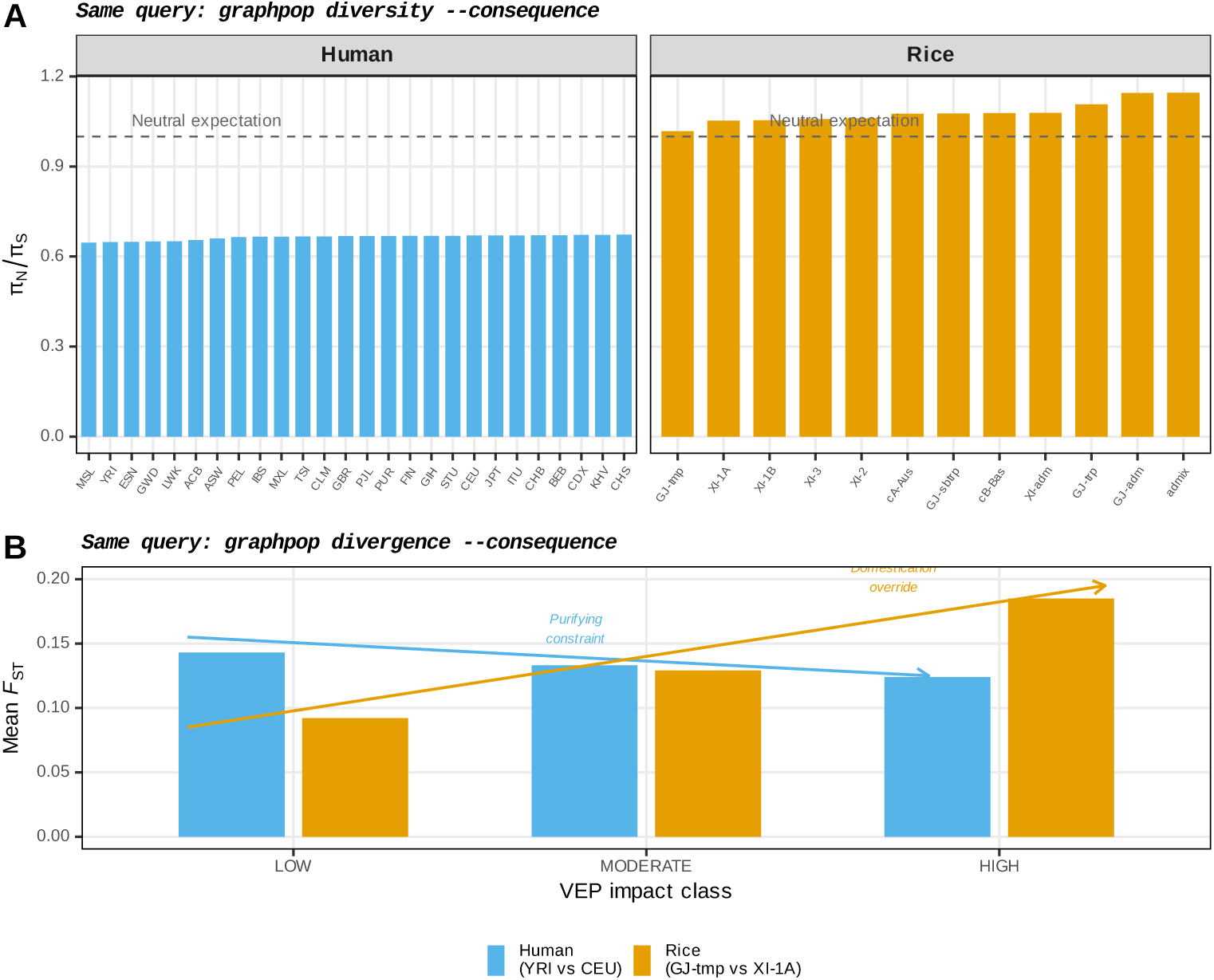
Same annotation-conditioned query applied to two species. **a**, *π_N_ /π_S_* computed with the identical graphpop diversity --consequence command for all human (26) and rice (12) populations. All rice subpopulations exceed the neutral expectation (*π_N_ /π_S_ >* 1.0), indicating universal relaxation of purifying selection under domestication; all human populations fall below 1.0, confirming efficient purifying selection under natural evolution. **b**, Mean Fst by VEP impact class for one representative population pair per species, computed with the same graphpop divergence --consequence command. Humans show decreasing Fst with increasing impact severity (purifying constraint), while rice shows increasing Fst (domestication override). The cross-species reversal emerges from the same single-line query applied to two datasets stored in the same graph schema.

**Fig. S7.**
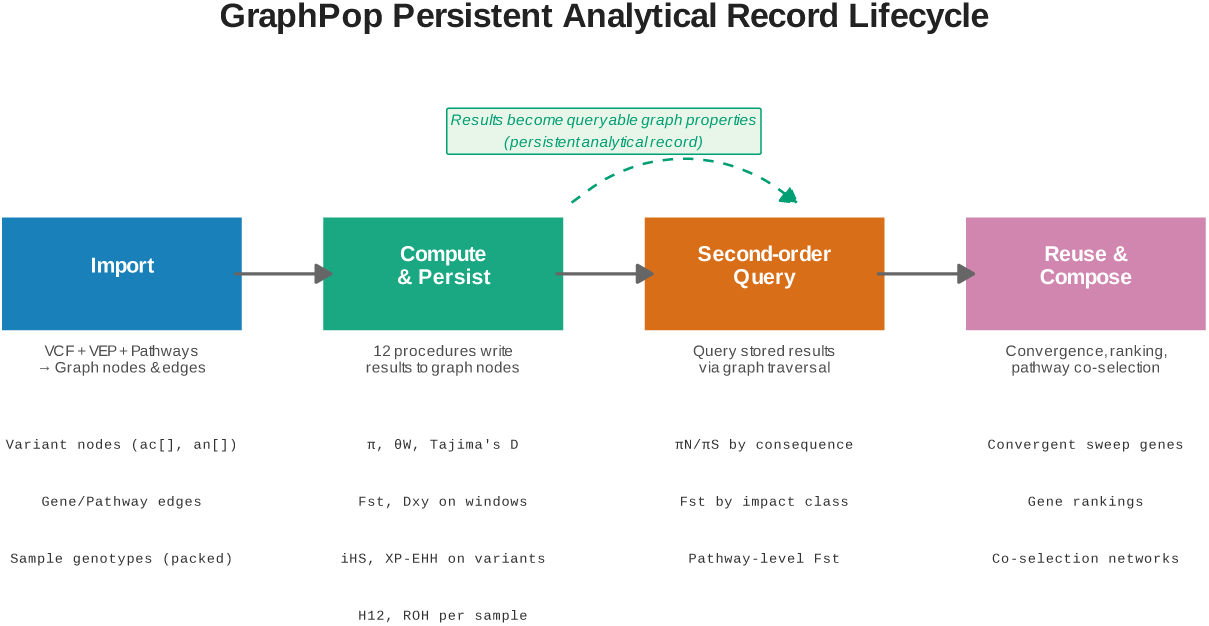
Persistent analytical record lifecycle. GraphPop’s four-stage analytical lifecycle. (1) Import: VCF genotypes, VEP annotations, and pathway data are loaded as graph nodes and edges. (2) Compute & Persist: 12 stored procedures write results (diversity, Fst, iHS, etc.) as properties on the same nodes that carry the biological data. (3) Second-order Query: stored results are queried by graph traversal for annotation-conditioned analyses (e.g., *π_N_ /π_S_* by consequence class, Fst by impact level). (4) Reuse & Compose: results computed for one purpose are reused for qualitatively different analyses (convergent sweep detection, gene ranking, pathway co-selection networks) without re-computation. The persistent analytical record—computed statistics stored as graph node properties—is the key architectural property that enables stages 3 and 4.

**Fig. S8.**
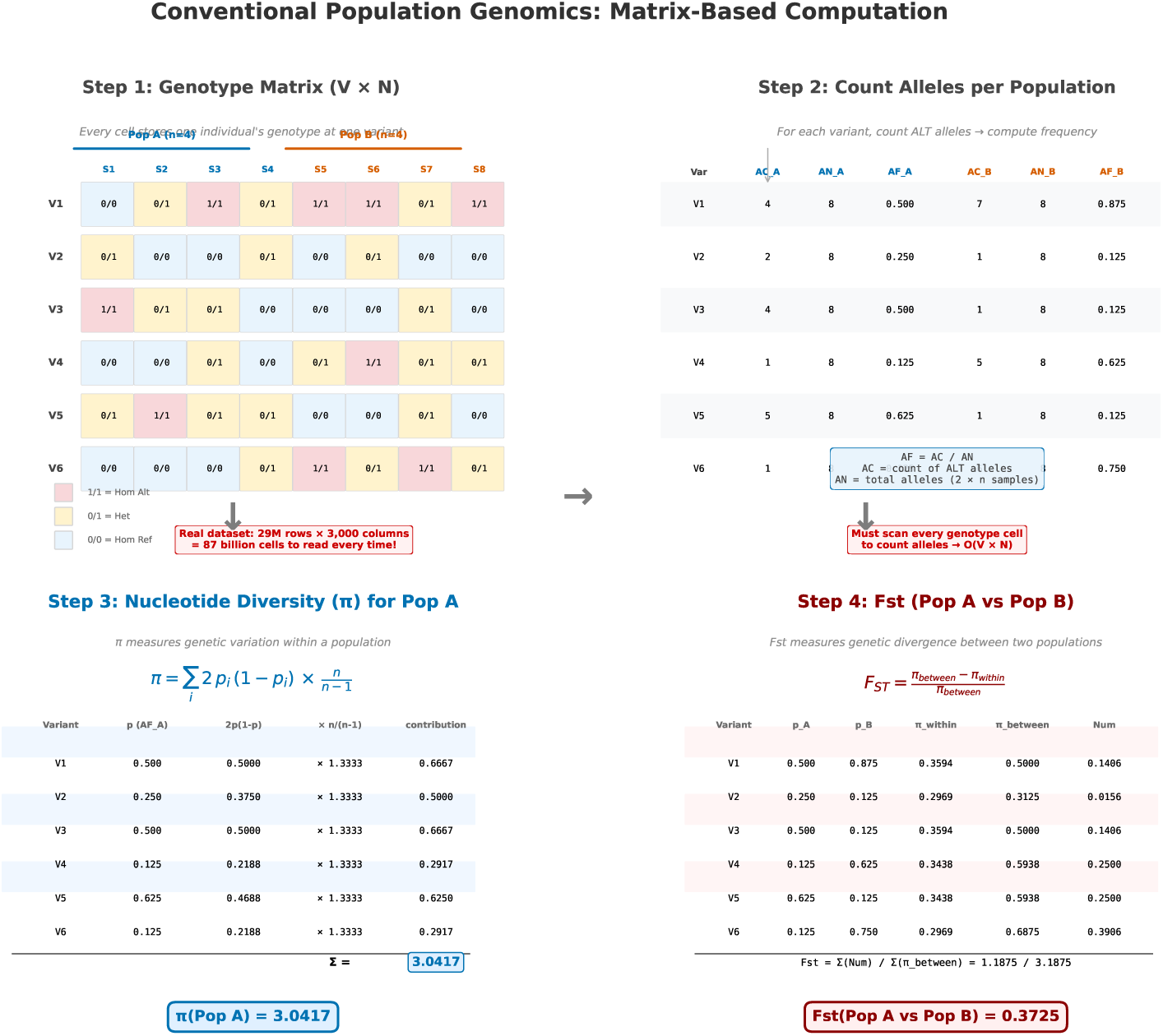
Conventional matrix-based computation of. *π* **and Fst—worked example.** Step-by-step illustration of the classical population genomics pipeline applied to a small example dataset (6 variants × 8 samples, split into Pop A and Pop B). **Step 1**: the genotype matrix stores one genotype per cell (0/0, 0/1, 1/1); the matrix size scales as *V* × *N* and reaches 87 billion cells for the rice 3K dataset. **Step 2**: every cell is scanned to count alternative alleles per population, yielding allele counts (AC), total alleles (AN), and frequencies (AF). **Step 3**: nucleotide diversity *π* for Pop A is computed by summing 2*p_i_*(1 − *p_i_*) × *n/*(*n* − 1) over variants. **Step 4**: Fst between Pop A and Pop B is computed from *π_within_* and *π_between_*. Every step requires reading the entire genotype matrix— O(*V* × *N* ) complexity—making repeated queries across populations and chromosomes expensive.

**Fig. S9.**
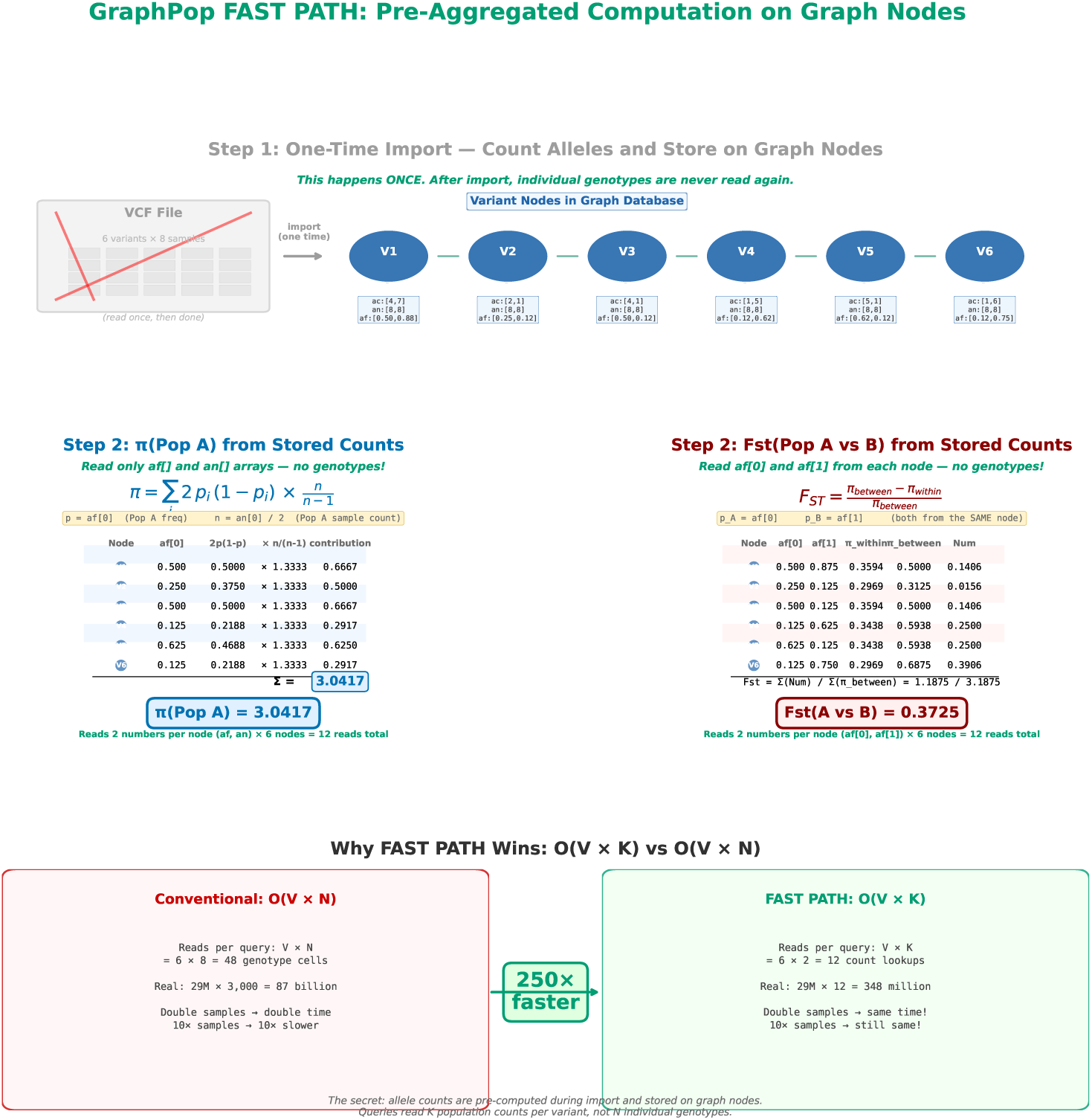
GraphPop FAST PATH computation of. *π* **and Fst—same example, different complexity.** The same six-variant example dataset as in Supplementary Fig. S8, computed via GraphPop’s FAST PATH. **Step 1** (one-time import): allele counts per population are pre-aggregated and stored as node properties (ac[], an[], af[]) on each Variant node; individual genotypes are never read again after import. **Step 2**: *π*(Pop A) is computed by reading only the af[0] and an[0] arrays from each Variant node—12 numerical reads instead of 48 genotype-cell scans. The result is identical to the conventional method (*π* = 3.0417). **Step 3**: Fst(Pop A vs Pop B) requires only af[0] and af[1] .per node—another 12 reads. The bottom panel contrasts the two scaling regimes: O(*V* × *N* ) reads grow with sample count, while O(*V* × *K*) reads stay fixed at the population dimension. For the rice 3K dataset, this is a ∼250× reduction in operations per query.

**Fig. S10.**
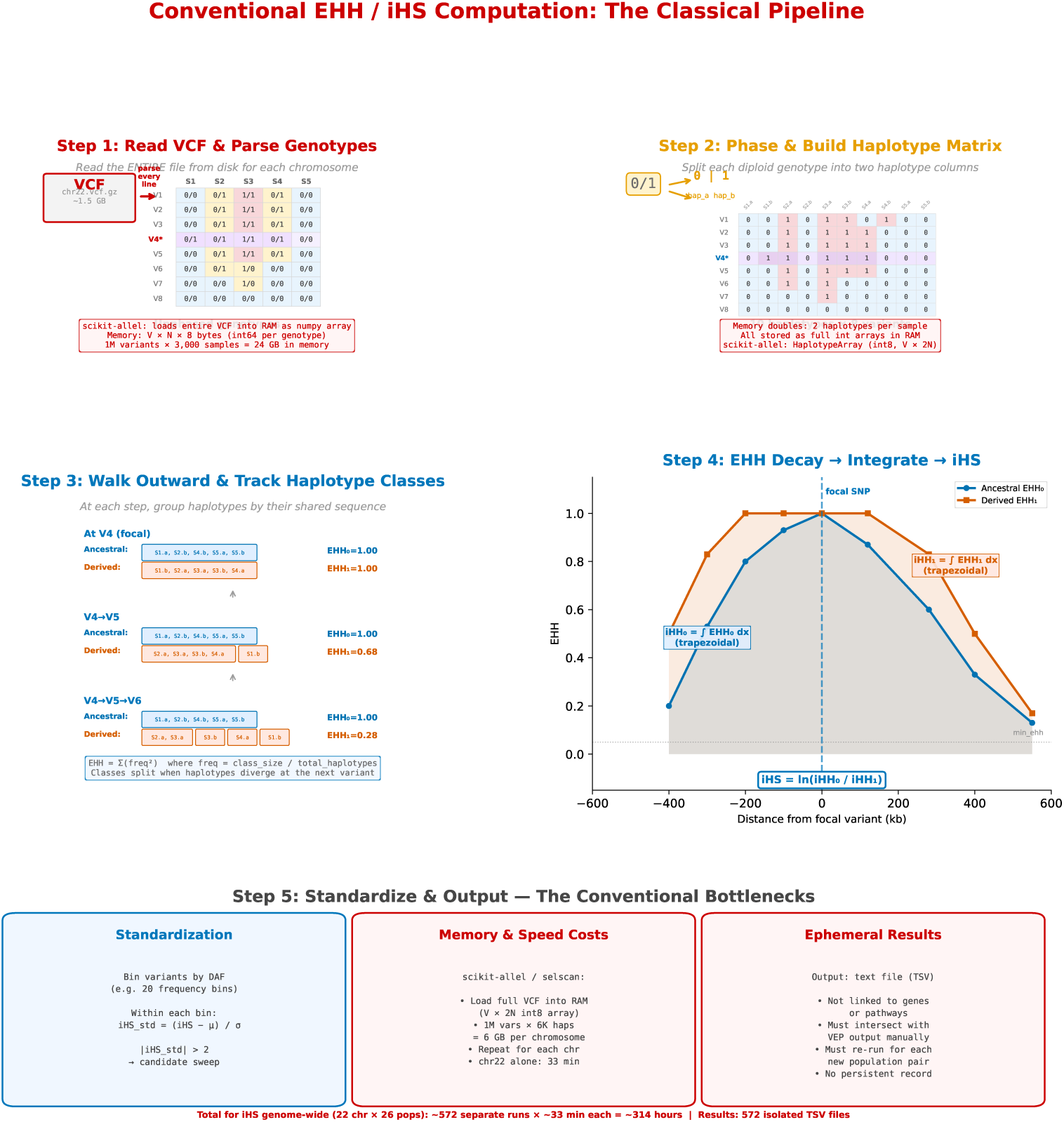
Conventional EHH and iHS computation—worked example. The classical pipeline for haplotype-based selection scans. **Step 1**: parse the entire VCF and load genotypes into RAM as a *V* × *N* int array (24 GB for 1M variants × 3,000 samples in scikit-allel). **Step 2**: phase each diploid genotype into two haplotype columns, doubling the memory footprint (HaplotypeArray of *V* × 2*N* ). **Step 3**: starting from a focal variant, walk outward in both directions; at each step, group haplotypes by their shared sequence and recompute the EHH statistic (sum of squared class frequencies). **Step 4**: integrate EHH curves for ancestral and derived alleles to obtain iHH_0_ and iHH_1_, then compute iHS as the log ratio. **Step 5**: standardize iHS within derived allele frequency bins. The classical approach loads the entire VCF into memory, processes one chromosome at a time, and writes results to isolated TSV files with no link back to gene or pathway annotations. Genome-wide iHS for 22 autosomes × 26 populations requires ∼572 separate runs and ∼314 hours of compute.

**Fig. S11.**
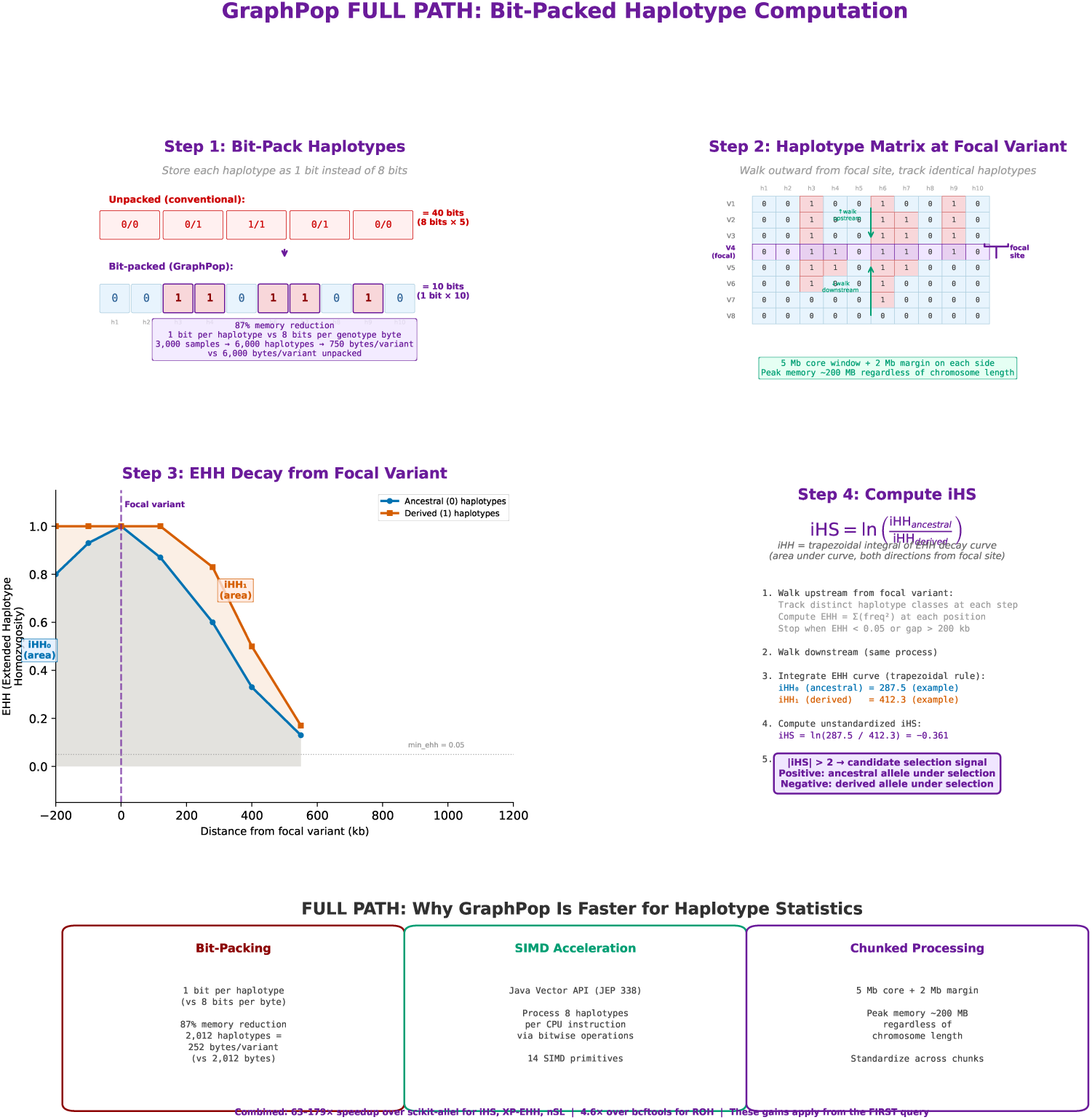
GraphPop FULL PATH computation of EHH and iHS—bit-packed haplotype computation. GraphPop’s FULL PATH for haplotype-based statistics. **Step 1**: bit-pack each haplotype as a single bit instead of 8 bits per byte, achieving 87% memory reduction (252 bytes per variant for 2,012 haplotypes vs 2,012 bytes unpacked). **Step 2**: load the haplotype matrix from the graph database into a dense bit-packed structure; walk outward from the focal variant. **Step 3**: compute EHH at each step using SIMD-accelerated bitwise operations (Java Vector API, JEP 338). **Step 4**: integrate EHH curves and compute iHS, then standardize within derived allele frequency bins. The bottom strip summarises the three pillars of FULL PATH performance: bit-packing (87% memory reduction), SIMD acceleration (8 haplotypes per CPU instruction), and chunked processing (5 Mb core windows with 2 Mb EHH margins, peak memory ∼200 MB regardless of chromosome length). Combined, these yield 63–179× speedups over scikit-allel for iHS, XP-EHH, and nSL—advantages that apply from the first query, independent of pre-aggregation.

**Fig. S12.**
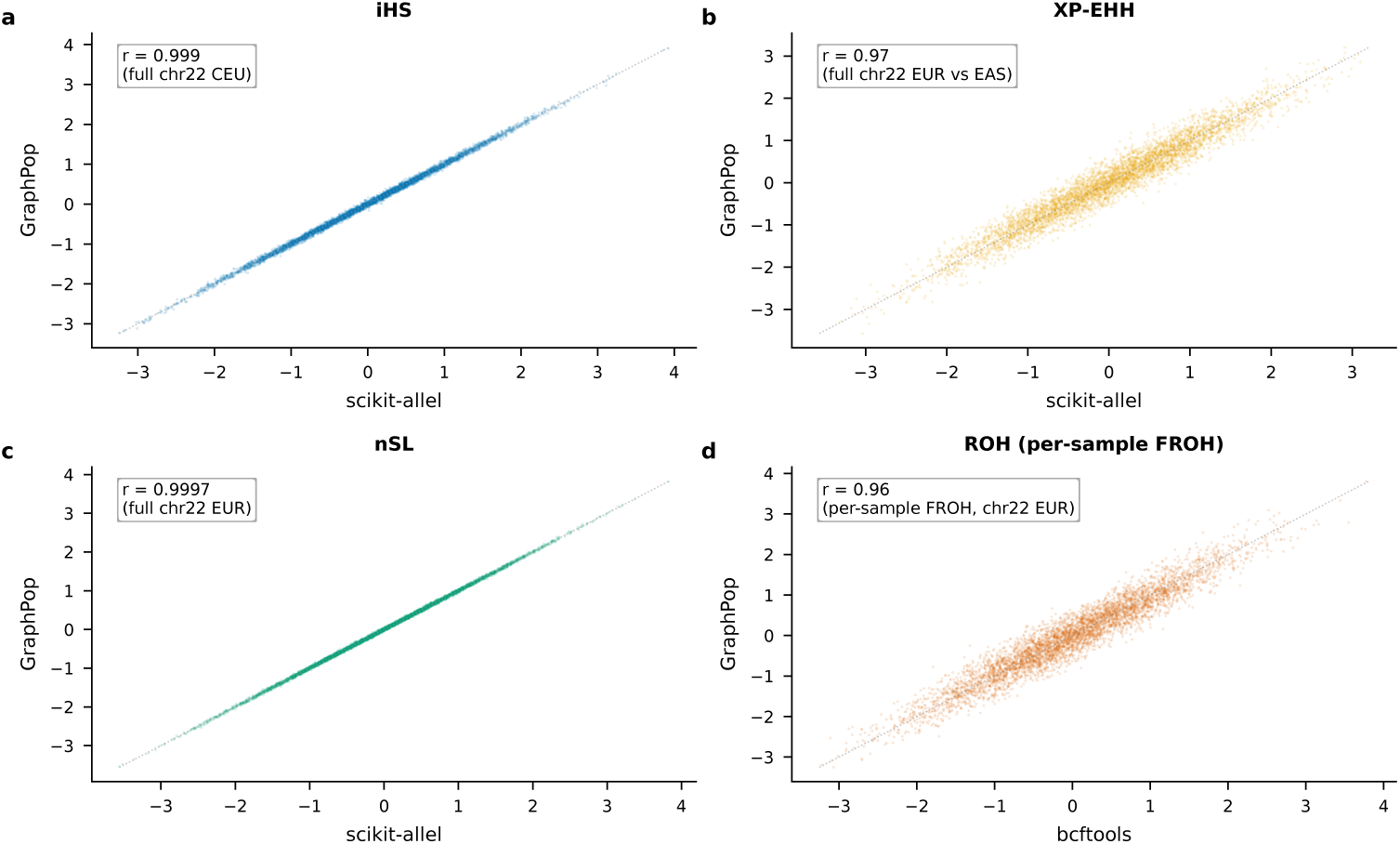
Per-locus validation of FULL PATH statistics. Scatterplots comparing GraphPop FULL PATH output against competitor tools on 1000 Genomes chr22. **a**, iHS (GraphPop vs scikitallel, CEU population, *r* = 0.999). **b**, XP-EHH (GraphPop vs scikit-allel, EUR vs EAS, *r* = 0.97). **c**, nSL (GraphPop vs scikit-allel, EUR, *r* = 0.9997). **d**, ROH (per-sample FROH, GraphPop vs bcftools, EUR, *r* = 0.96). The near-unity correlations for iHS and nSL confirm numerical equivalence. The lower XP-EHH correlation (*r* = 0.97) reflects differences in variant filtering (GraphPop requires MAF ≥ 0.05 in at least one population) and EHH truncation thresholds. The ROH correlation (*r* = 0.96) reflects differences in HMM transition rate parameterization between GraphPop and bcftools. In all four panels, the identity line (grey dotted) and regression line (black dashed) are shown.

## References

1. Miles, A., et al. scikit-allel: a Python package for exploring and analysing genetic variation data. https://github.com/cggh/scikit-allel (2024). V1.3.7.

[2] Danecek, P. et al. The variant call format and VCFtools. Bioinformatics 27, 2156–2158 (2011).

[3] Purcell, S. et al. PLINK: a tool set for whole-genome association and population-based linkage analyses. American Journal of Human Genetics 81, 559–575 (2007).

[4] Chang, C. C. et al. Second-generation PLINK: rising to the challenge of larger and richer datasets. GigaScience 4, 7 (2015).

[5] Lu, J. et al. The accumulation of deleterious mutations in rice genomes: a hypothesis on the cost of domestication. Trends in Genetics 22, 126–131 (2006).

[6] McLaren, W. et al. The Ensembl variant effect predictor. Genome Biology 17, 122 (2016).

[7] Quinlan, A. R. & Hall, I. M. BEDTools: a flexible suite of utilities for comparing genomic features. Bioinformatics 26, 841–842 (2010).

[8] Voight, B. F., Kudaravalli, S., Wen, X. & Pritchard, J. K. A map of recent positive selection in the human genome. PLoS Biology 4, e72 (2006).

[9] Sabeti, P. C. et al. Genome-wide detection and characterization of positive selection in human populations. Nature 449, 913–918 (2007).

[10] Garud, N. R., Messer, P. W., Buzbas, E. O. & Petrov, D. A. Recent selective sweeps in North American *Drosophila melanogaster* show signatures of soft sweeps. PLoS Genetics 11, e1005004 (2015).

[11] Weir, B. & Cockerham, C. C. Estimating F-statistics for the analysis of population structure. Evolution 38, 1358–1370 (1984).

[12] Robinson, I., Webber, J. & Eifrem, E. Graph Databases: New Opportunities for Connected Data 2nd edn (O’Reilly Media, 2015).

[13] Angles, R. & Gutierrez, C. Survey of graph database models. ACM Computing Surveys 40, 1–39 (2008).

[14] Angles, R. et al. Foundations of modern query languages for graph databases. ACM Computing Surveys 50, 1–40 (2017).

[15] Gillespie, M. et al. The Reactome pathway knowledgebase 2022. Nucleic Acids Research 50, D588–D592 (2022).

[16] Nickel, M., Murphy, K., Tresp, V. & Gabrilovich, E. A review of relational machine learning for knowledge graphs. Proceedings of the IEEE 104, 11–33 (2016).

[17] Narasimhan, V. et al. BCFtools/RoH: a hidden Markov model approach for detecting autozygosity from next-generation sequencing data. Bioinformatics 32, 1749–1751 (2016).

18. 1000 Genomes Project Consortium. A global reference for human genetic variation. Nature 526, 68–74 (2015).

[19] Wang, W. et al. Genomic variation in 3,010 diverse accessions of Asian cultivated rice. Nature 557, 43–49 (2018).

20. Hail Team. Hail 0.2. https://hail.is (2024). Accessed 2026.

[21] Ohta, T. Slightly deleterious mutant substitutions in evolution. Nature 246, 96–98 (1973).

[22] Eyre-Walker, A. & Keightley, P. D. The distribution of fitness effects of new mutations. Nature Reviews Genetics 8, 610–618 (2007).

[23] Huang, X. et al. A map of rice genome variation reveals the origin of cultivated rice. Nature 490, 497–501 (2012).

[24] Sweeney, M. & McCouch, S. The complex history of the domestication of rice. Annals of Botany 100, 951–957 (2007).

[25] Naithani, S. et al. Plant Reactome: a knowledgebase and resource for comparative pathway analysis. Nucleic Acids Research 48, D1093–D1103 (2020).

[26] Vesoft Inc. NebulaGraph: an open-source distributed graph database. https://nebula-graph.io/ (2025). Accessed 2026-04-08.

27. Neo4j, Inc. Neo4j AuraDB: fully managed cloud graph database. https://neo4j.com/cloud/auradb/ (2025). Accessed 2026-04-08.

[28] Korunes, K. L. & Samuk, K. pixy: unbiased estimation of nucleotide diversity and divergence in the presence of missing data. Molecular Ecology Resources 21, 1359–1368 (2021).

[29] Kelleher, J. et al. Inferring whole-genome histories in large population datasets. Nature Genetics 51, 1330–1338 (2019).

[30] Estaji, E., Zhao, S.-W., Chen, Z.-Y., Nie, S. & Mao, J.-F. GraphMana: graphnative data management for population genomics projects. *bioRxiv* (2026). Preprint.

31. Neo4j, Inc. Neo4j graph database. https://neo4j.com (2024). Community Edition, version 2026.01.4.

[32] Deutsch, A. et al. Graph pattern matching in GQL and SQL/PGQ. Proc. ACM SIGMOD Int. Conf. Manag. Data 2246–2258 (2022).

[33] Pedersen, B. S. & Quinlan, A. R. cyvcf2: fast, flexible variant analysis with Python. Bioinformatics 33, 1867–1869 (2017).

[34] Tajima, F. Statistical method for testing the neutral mutation hypothesis by DNA polymorphism. Genetics 123, 585–595 (1989).

[35] Zeng, K., Fu, Y.-X., Shi, S. & Wu, C.-I. Statistical tests for detecting positive selection by utilizing high-frequency variants. Genetics 174, 1431–1439 (2006).

[36] Yi, X. et al. Sequencing of 50 human exomes reveals adaptation to high altitude. Science 329, 75–78 (2010).

[37] Ferrer-Admetlla, A., Liang, M., Korneliussen, T. & Nielsen, R. On detecting incomplete soft or hard selective sweeps using haplotype structure. Molecular Biology and Evolution 31, 1275–1291 (2014).

[38] Wigginton, J. E., Cutler, D. J. & Abecasis, G. R. A note on exact tests of Hardy–Weinberg equilibrium. American Journal of Human Genetics 76, 887–893 (2005).

[39] Bycroft, C. et al. The UK Biobank resource with deep phenotyping and genomic data. Nature 562, 203–209 (2018).

[40] Karczewski, K. J. et al. The mutational constraint spectrum quantified from variation in 141,456 humans. Nature 581, 434–443 (2020).

[41] Hayes, B. J. & Daetwyler, H. D. 1000 Bull Genomes Project to map simple and complex genetic traits in cattle: applications and outcomes. Annual Review of Animal Biosciences 7, 89–102 (2019).

[42] Kijas, J. W. et al. Genome-wide analysis of the world’s sheep breeds reveals high levels of historic mixture and strong recent selection. PLoS Biology 10, e1001258 (2012).

[43] Neale, D. B. & Kremer, A. Forest tree genomics: growing resources and applications. Nature Reviews Genetics 12, 111–122 (2011).

[44] Lamichhaney, S. et al. Evolution of Darwin’s finches and their beaks revealed by genome sequencing. Nature 518, 371–375 (2015).

[45] Zhao, K. et al. An Arabidopsis example of association mapping in structured populations. PLoS Genetics 3, e4 (2007).

