## Supplementary material for "GraphPop: graph-native computation decouples population genomics complexity from sample count": all supplementary materials

### Supplementary Information for GraphPop: graph-native computation decouples population genomics complexity from sample count

Ehsan Estaji[1], Shi-Wei Zhao[1], Zhao-Yang Chen[1], Shuai Nie[2], Jian-Feng Mao[1*]

[1]Umeå Plant Science Centre, Department of Plant Physiology, Umeå University, SE-90187, Umeå, Sweden.
[2]Rice Research Institute, Guangdong Academy of Agricultural Sciences, 510640, Guangzhou, China.

Contributing authors:;

**Abstract**

Supplementary Notes, Tables, and Figures for the GraphPop manuscript.

Summary diversity statistics across superpopulations confirm expected continental patterns:

- **African populations** show the highest diversity (mean $\pi = 0.051$, $\theta_W = 0.061$), consistent with the larger long-term effective population size. MSL is the most diverse single population ($\pi = 0.052$).
- **Out-of-Africa populations** show progressively reduced diversity: SAS ($\pi = 0.039$), AMR ($\pi = 0.040$), EUR ($\pi = 0.038$), EAS ($\pi = 0.036$, lowest; PEL is the least diverse population at $\pi = 0.036$).
- **Tajima's** $D$ varies by continent: AFR ($D = -0.52$, population expansion), AMR ($-0.39$), SAS ($-0.07$), EUR ($+0.09$), EAS ($+0.20$, population structure/admixture).

- PC1 (57.0% variance): diversity axis—African samples highest on heterozygosity and rare burden
- PC2 (30.9% variance): inbreeding axis—South Asian samples highest on FROH
- FROH vs heterozygosity rate: $\rho = -0.394$, $p = 1.2 \times 10^{-29}$
- Individual PC1 vs population PC1: $\rho = 0.74$ (Spearman), validating the approach

Per-group mean statistics (chr22 features):

| Group | $n$ | FROH | Het rate | Rare burden | Rare missense |
|---|---|---|---|---|---|
| African | 552 | 0.0047 | 0.0453 | 4,044 | 34.5 |
| European | 491 | 0.0066 | 0.0345 | 1,249 | 16.8 |
| East Asian | 552 | 0.0069 | 0.0317 | 1,358 | 18.3 |
| South Asian | 489 | 0.0136 | 0.0354 | 1,597 | 21.0 |
| American | 518 | 0.0102 | 0.0351 | 1,353 | 16.8 |

#### Outlier analysis

Z-distance from population centroid in the 5-dimensional process space identified extreme individuals. The top 5 outliers:

| Sample | Population | Group | $z$-distance | FROH |
|---|---|---|---|---|
| HG01816 | CDX | East Asian | 9.50 | 0.125 |
| HG02701 | Unknown | — | 8.55 | 0.125 |
| NA20585 | TSI | European | 8.04 | 0.055 |
| HG02659 | Unknown | — | 7.72 | 0.098 |
| HG02692 | Unknown | — | 7.26 | 0.095 |

The top outlier (HG01816) carries FROH = 0.125—18× the CDX population mean—consistent with recent close-relative mating. Four UNKNOWN-population samples appeared in the top 10, suggesting mislabeling or recent admixture. Z-distance weakly predicts FROH ($\rho = -0.087$; Supplementary Fig. S3).

#### Pathway-stratified purifying selection gradient

Genome-wide missense variant positions (224,181 across 22 autosomes) were classified as belonging to constrained pathways (lowest-quartile mean Fst: $\leq 0.108$; $n = 15{,}092$ sites) or divergent pathways (highest-quartile Fst: $\geq 0.158$; $n = 21{,}886$ sites) via `IN_PATHWAY` graph traversal (Supplementary Fig. S5). Key findings:

- African 2× enrichment holds in both constrained (2.04×, $p = 1.6 \times 10^{-290}$) and divergent (1.95×, $p = 3.0 \times 10^{-265}$) pathways
- FROH predicts individual rare missense burden: $\rho = -0.211$, $p = 1.6 \times 10^{-33}$ (genome-wide)
- FROH does *not* predict constrained:divergent burden ratio: $\rho = +0.004$, $p = 0.84$
- Conclusion: bottleneck-driven depletion of rare functional variants is genome-wide and uniform across pathway classes

#### R09: Multi-statistic correlations

Pairwise correlations among population-level summary statistics across 12 populations revealed: $\pi$ vs $\theta_W$: $\rho = 0.90$; $\pi$ vs number of sweeps: $\rho = -0.82$ ($p = 0.001$); FROH vs number of sweeps: $\rho = 0.75$ ($p = 0.005$). These correlations confirm that bottleneck-driven diversity loss and sweep accumulation are tightly coupled in domesticated rice.

#### R10: Fay & Wu's *H* and DAF enrichment

Derived allele frequency (DAF) spectra at PBS-outlier loci showed enrichment of high-frequency derived alleles (DAF enrichment ratio = 1.53), consistent with positive selection driving alleles toward fixation.

#### R11: Multi-evidence synthesis

A multi-evidence scoring framework classified subpopulation pairs into HIGH/MEDIUM/LOW evidence tiers for divergent selection, combining $\pi_N/\pi_S$, conditioned Fst, XP-EHH peaks, and PBS signals. The indica–japonica split received the highest evidence scores across all metrics.

#### R12: ROH–sweep correlation

Population-level correlations between ROH metrics and sweep counts: $\pi$ vs number of sweeps $\rho = -0.82$ ($p = 0.001$); FROH vs number of sweeps $\rho = 0.75$ ($p = 0.005$); $\pi$ vs hard sweep fraction $\rho = 0.58$ ($p = 0.05$). The inverse relationship between diversity and sweep count confirms that bottleneck-driven loss of genetic variation facilitates hard sweeps.

#### R18: GO term enrichment (graph database)

GO term enrichment among sweep genes (Garud $H_{12} > 0.1$ in any population; 4,953 genes, 1,264 GO terms tested): 10 terms were globally significant (Bonferroni-corrected $p < 0.05$), including endoplasmic reticulum (5.3-fold, $p = 5.8 \times 10^{-5}$), calmodulin binding (5.1-fold, $p = 8.4 \times 10^{-5}$), and nucleus (1.75-fold, $p = 1.1 \times 10^{-4}$). Population-specific enrichment was strongest in GJ-tmp (13/126 significant terms), consistent with its extreme bottleneck.

#### Database sizes and import times

| Dataset | Variants | Samples | DB size | Import time |
|---|---|---|---|---|
| 1000G chr22 | 1.07M | 3,202 | ∼3 GB | ∼5 min |
| 1000G full genome | 70.7M | 3,202 | ∼180 GB | ∼8 h |
| Rice 3K | 29.6M | 3,024 | ∼46 GB | ∼4 h |

### Supplementary Note S6: EHH and ROH Computation Details

#### EHH computation

EHHComputer walks position-sorted haplotype arrays from a focal variant outward (both directions), tracking distinct haplotype classes. The implementation uses flat array tracking with incremental sum-of-pairs updates, replacing the earlier HashMap-per-step approach.

Key configurable parameters:

- `min_ehh`: EHH truncation threshold (default 0.05)—walk stops when EHH decays below this value
- `max_gap`: Maximum physical gap before aborting (default 200 kb)—prevents unreliable integration across assembly gaps
- `gap_scale`: Distance cap for trapezoidal integration segments (default 20 kb)

iHH is computed as the trapezoidal integral of the EHH decay curve. For iHS, the ratio $\log(\mathrm{iHH}_{\mathrm{ancestral}}/\mathrm{iHH}_{\mathrm{derived}})$ is standardized within allele-frequency bins (default 20 bins). For XP-EHH, $\log(\mathrm{iHH}_{\mathrm{pop1}}/\mathrm{iHH}_{\mathrm{pop2}})$ is standardized genome-wide.

#### Memory management for EHH procedures

Chromosomes are processed in chunks: 5 Mb core window + 2 Mb EHH margin on each side. This bounds peak memory at ∼200 MB per chunk regardless of chromosome length. Standardization is applied across all chunks after computation, preserving genome-wide distributional properties.

#### ROH HMM

The ROH HMM implements a two-state (Autozygous/Hardy–Weinberg) Viterbi decoder following Narasimhan et al.[5]. Transition probabilities use exact 2-state continuous-time Markov chain (CTMC) matrix exponentiation:

$$P(\mathrm{HW} \to \mathrm{AZ}|d) = \pi_{\mathrm{AZ}} \cdot (1 - (1 - \alpha - \beta)^d)$$

where $d$ is the inter-variant distance in base pairs.

Emission probabilities are allele-frequency weighted:

$$P(\mathrm{hom}|\mathrm{AZ}) = 1 - \epsilon \quad \text{(with het error rate } \epsilon = 10^{-3}\text{)} \tag{1}$$

$$P(\mathrm{hom}|\mathrm{HW}) = p^2 + (1-p)^2 \tag{2}$$

Default per-bp rates match bcftools roh -G30: `hw_to_az` $= 6.7 \times 10^{-8}$, `az_to_hw` $= 5 \times 10^{-9}$.

#### Validation

- iHS[6]: $r = 0.999$ vs scikit-allel[7] on 1000 Genomes chr22 (CEU)

- XP-EHH[8]: $r = 0.97$ vs scikit-allel
- nSL[9]: $r = 0.9997$ vs scikit-allel
- ROH: $r = 0.96$ per-sample total length vs bcftools roh[5]; $r = 0.87$ vs bcftools genome-wide; $r = 0.34$ vs PLINK 1.9[10] (which uses a sliding-window algorithm rather than HMM)

### Supplementary Note S7: Benchmark Methodology

#### Benchmark protocol

All benchmarks were run on a single workstation with the following specifications: AMD processor, 64 GB RAM, NVMe SSD storage, running Ubuntu. The graph database was configured with 4 GB heap and 16 GB page cache. Each benchmark was run 3 times with 1 warm-up run discarded; the median of the 2 timing runs is reported.

GraphPop was called via the graph database Python driver with the database pre-loaded in memory (warm cache). Competitor tools read from their native file formats: VCF for VCFtools, scikit-allel, and bcftools; PLINK binary (.bed/.bim/.fam) for PLINK 2 and PLINK 1.9.

#### Region sizes

Four region sizes were benchmarked for each statistic:

- 100 kb: chr22:20,000,000–20,100,000
- 1 Mb: chr22:20,000,000–21,000,000
- 10 Mb: chr22:20,000,000–30,000,000
- Full chromosome: chr22 (entire, ~51 Mb, 1,070,401 variants)

#### Memory measurement

Peak memory was measured using:

- GraphPop graph database path: `graphpop.heap_stats()` procedure (JVM heap), plus baseline graph database memory (~160 MB)
- GraphPop-numpy path: Python `tracemalloc` peak
- External tools: `/usr/bin/time -v` peak RSS

#### Competitor tool versions

scikit-allel[7] v1.3.7, VCFtools[11] v0.1.16, PLINK 2[12] v2.00a6, PLINK 1.9[10] v1.90b7.2, bcftools[5] v1.21.

#### Full benchmark results

Full-chromosome (chr22) speedups of GraphPop's best variant vs the fastest non-GraphPop tool:

| Statistic | GraphPop (s) | Best competitor (s) | Speedup | Competitor |
|---|---|---|---|---|
| $\pi/\theta_W$/Tajima's $D$ | 12.1 | 1,757 | 146× | scikit-allel |
| Fst/$D_{xy}$ | 8.8 | 2,165 | 245× | scikit-allel |
| SFS | 5.9 | 1,937 | 327× | scikit-allel |
| iHS | 10.9 | 1,948 | 179× | scikit-allel |
| XP-EHH | 16.7 | 2,280 | 136× | scikit-allel |
| nSL | 35 (numpy) | 2,231 | 63× | scikit-allel |
| ROH | 10.9 (numpy) | 128 (graph DB) | 4.6× | bcftools |
| LD ($r^2$) | 6.4 | 1.1 | 0.17× | PLINK 2 |

GraphPop is fastest or tied on 6 of 8 statistics. PLINK 2 is faster for pairwise LD owing to its highly optimized native binary format (pgen).

#### End-to-end versus amortised performance

The query-time speedups reported in Table 1 measure performance *after* one-time database import. To provide a complete picture, we report end-to-end costs including import:

| **Phase** | **chr22** | **Full genome (22 autosomes)** |
|---|---|---|
| VCF import + allele-count aggregation | ∼12 min | ∼4.5 h |
| VEP annotation loading | ∼2 min | ∼30 min |
| Reactome/GO pathway loading | <1 min | <1 min |
| *Break-even analysis (single population, FAST PATH):* | | |
| Queries to amortise import (vs scikit-allel) | 1 | 3–5 |
| Queries to amortise import (vs VCFtools) | 1 | 8–12 |

For a single FAST PATH query on chr22, GraphPop's import + query time (∼12 min + 12 s) exceeds scikit-allel's computation time (∼30 min) but the import cost is paid once. After import, each subsequent query (e.g., a different population, chromosome, or annotation condition) takes seconds rather than minutes. For the rice 3K full-genome analysis (12 populations × 12 chromosomes × 6 FAST PATH statistics = 864 queries), GraphPop's amortised time per query is <20 s versus ∼30 min per scikit-allel invocation. The amortised advantage grows with the number of queries, making GraphPop particularly suited to iterative, multi-population, annotation-conditioned investigations.

FULL PATH speedups do not depend on pre-aggregation and are realised from the first query. The bit-packed haplotype encoding and SIMD-accelerated kernels provide genuine per-computation performance gains: iHS completes in 10.9 s versus 1,948 s for scikit-allel regardless of whether the database was freshly imported or has been running for months.

#### Sensitivity analysis for rice $\pi_N/\pi_S$

The finding that all 12 rice subpopulations show $\pi_N/\pi_S > 1.0$ is a central biological result. We performed the following robustness checks to confirm that this finding is not an artifact of variant filtering, annotation bias, or sampling:

**Allele frequency filtering.** We recomputed $\pi_N/\pi_S$ after (i) removing singletons (minor allele count = 1), (ii) removing singletons and doubletons (MAC $\leq$ 2), and (iii) restricting to common variants (MAF > 0.05). All 12 subpopulations retained $\pi_N/\pi_S > 1.0$ under conditions (i) and (ii). Under condition (iii), ratios decreased modestly (range: 0.97–1.08) as expected, since common-variant $\pi_N/\pi_S$ is dominated by older, less deleterious mutations; 9 of 12 populations remained above 1.0.

**Call rate filtering.** Restricting to variants with call rate > 90% (excluding sites with > 10% missing data) did not alter the rank order of populations or the universal > 1.0 finding (range: 1.015–1.139).

**Per-chromosome consistency.** $\pi_N/\pi_S$ was computed for each of the 12 chromosomes independently. The > 1.0 finding held for all 144 population × chromosome combinations (range: 0.91–1.42), with 140 of 144 (97.2%) exceeding 1.0. The 4 exceptions were short chromosomes in GJ-tmp (the most bottlenecked population), consistent with sampling variance rather than systematic bias.

**Annotation sensitivity.** We compared results using VEP "missense_variant" versus "nonsynonymous_variant" definitions. Both produced qualitatively identical patterns ($r = 0.998$ across populations). We also verified that the missense and synonymous variant counts per population are sufficiently large (> 50,000 each genome-wide) to produce stable ratio estimates.

**Cross-species control.** The same analysis applied to all 26 human populations yielded $\pi_N/\pi_S = 0.646$–$0.673$ (all < 1.0), confirming that the > 1.0 finding in rice is biologically meaningful rather than a methodological artifact.

### Supplementary Note S8: GraphPop CLI Architecture

GraphPop provides a self-contained command-line interface (`graphpop`) that enables users to perform the entire population genomics workflow—from database setup through analysis to publication-ready figures—without knowledge of graph databases, graph query languages, or Python. The 12 stored procedures are listed in Supplementary Table S8. The CLI comprises 62 commands organised into 11 functional domains (Supplementary Fig. S1):

| Domain | Commands | Key functionality |
|---|---|---|
| Infrastructure & Server | 4 | Download/configure graph database, start/stop, status check |
| Database Management | 8 | Import VCF, dump/load databases, db list/create/switch/drop/info |
| Configuration & Validation | 6 | Config init/show/set, validate, inventory |
| Procedures (FAST PATH) | 6 | diversity, divergence, sfs, joint-sfs, genome-scan, pop-summary |
| Procedures (FULL PATH) | 6 | ihs, xpehh, nsl, roh, garud-h, ld |
| Conditioned Analysis | 1 | filter (post-hoc annotation filtering of persisted statistics) |
| Annotation Lookup | 5 | lookup {gene, pathway, variant, region}, neighbors |
| Multi-Stat Integration | 3 | converge, rank-genes, compare |
| Orchestration & Aggregation | 5 | run-all, aggregate, batch, report, export-windows |
| Data Extraction & Export | 4 | extract {variants, samples, genotypes}, export-bed |
| Visualisation | 11 | plot {diversity-bar, fst-heatmap, manhattan, pinpis, sfs-plot, roh-landscape, gene-zoom, chromosome, pop-tree, pca-scatter, heatmap} |
| Utility | 1 | query (arbitrary graph queries) |

#### Design principles

Three design principles govern the CLI: (1) **Familiar interface**: commands follow the subcommand pattern of VCFtools and PLINK (`graphpop diversity chr22 EUR -o output.tsv`), producing tab-separated output to stdout or file by default. (2) **Dual output mode**: the `--persist` flag writes results to graph nodes for downstream cross-query; without it, the tool behaves as a conventional command-line program that does not require a persistent database. (3) **Annotation conditioning**: FAST PATH procedures accept `--consequence`, `--pathway`, and `--gene` flags for direct conditioning; FULL PATH procedures use a compute-then-filter workflow (`graphpop ihs --persist` followed by `graphpop filter ihs --consequence missense_variant`).

#### Key user-facing commands beyond procedures

`graphpop converge` finds genomic regions where multiple independently computed statistics (e.g., iHS, XP-EHH, $H_{12}$, Fst) simultaneously exceed user-specified thresholds—the multi-statistic convergence analysis that is a signature capability of graph-native investigation. `graphpop rank-genes` produces a ranked gene list by composite selection evidence (combining max |iHS|, max |XP-EHH|, max $H_{12}$, mean Fst, and HIGH-impact variant count per gene). `graphpop lookup` enables graph exploration without writing graph queries: users can query a gene (`graphpop lookup` gene KCNE1) to retrieve variant counts, consequence types, pathway memberships, and persisted selection statistics. graphpop report generates a self-contained HTML analysis summary. All 11 plot types follow Nature Methods figure guidelines (white background, Wong/Okabe-Ito colorblind palette, 183 mm double-column width).

##### End-to-end workflow

A complete analysis requires no tools beyond the graphpop command:

```
graphpop setup --password mypass        # install graph database
graphpop import --vcf data.vcf.gz \     # import data
  --panel panel.txt --database mydb
graphpop run-all -d results/            # full-genome analysis
graphpop converge --stats ihs,xpehh,h12 \
  --thresholds 2,2,0.3 --pop POP       # convergent signals
graphpop rank-genes --pop POP --top 50  # ranked gene list
graphpop plot manhattan ihs.tsv \
  --stat ihs --threshold 2.5 -o fig.pdf # publication figure
graphpop report -o summary.html         # automated report
graphpop dump --database mydb           # share database
```

Full documentation, including per-command manuals and two vignettes (rice 3K and human 1000 Genomes end-to-end tutorials), is available in the repository.

##### Programmatic access via MCP server

In addition to the CLI, GraphPop provides a Model Context Protocol (MCP) server (graphpop-mcp) that exposes 21 tools for programmatic access by AI agents and external applications. The MCP server wraps the same 12 procedures as the CLI, plus 7 high-level analytical tools (converge, rank_genes, lookup_gene, lookup_pathway, lookup_region, filter, inventory) and 2 utilities (status, query). All tools accept typed parameters and return JSON results, enabling large language model agents to compute population genetics statistics, condition on annotations, detect convergent selection signals, rank candidate genes, and explore graph topology autonomously. The server uses the same connection configuration as the CLI (GRAPHPOP_URI, GRAPHPOP_DATABASE environment variables or ~/.graphpop/config.yaml), ensuring consistent behaviour across interfaces.

#### Supplementary Note S9: Dataset Scale and Applicability

##### The landscape of population genomics dataset sizes

The public discourse around genomic data scaling is heavily influenced by human biobank projects (UK Biobank ∼500,000 genomes[13], gnomAD 141,456 exomes/genomes[14], All of Us > 1,000,000 planned). However, the vast majority of population genomics research worldwide operates at scales well within GraphPop's current single-workstation capacity. Table S1 summarises typical dataset sizes across the major domains of population genomics.

**Table S1** Typical population genomics dataset sizes across research domains. $N$ = number of individuals/accessions; $V$ = number of variant sites. GraphPop's $O(V \times K)$ complexity reduction applies to all domains listed.

| **Domain** | **Typical** $N$ | **Typical** $V$ | **Representative datasets** |
|---|---|---|---|
| Crop breeding & genetics | 200–10,000 | 1M–30M | Rice 3K[3], Maize HapMap, Wheat 10K |
| Livestock genetics | 500–50,000 | 5M–20M | 1000 Bull Genomes[15], Sheep HapMap[16] |
| Aquaculture genetics | 100–5,000 | 1M–15M | Salmon, shrimp, tilapia breeding panels |
| Forest genomics | 200–3,000 | 2M–20M | Poplar, eucalyptus, spruce[17] |
| Conservation biology | 50–500 | 1M–10M | Endangered species resequencing panels |
| Wild population ecology | 100–5,000 | 1M–30M | Darwin's finches[18], stickleback, cichlids |
| Plant model organisms | 500–3,000 | 5M–20M | Arabidopsis 1001 Genomes[19] |
| Human pop-gen (non-biobank) | 100–10,000 | 10M–80M | 1000 Genomes[1], HGDP |
| Plant pangenomes | 500–5,000 | 10M–100M | Rice, tomato, soybean pangenomes |
| Microbial population genomics | 100–10,000 | 100K–5M | Bacterial GWAS panels |
| *Human biobanks* | *100K–1M+* | *10M–700M* | *UK Biobank, gnomAD[14], All of Us* |

GraphPop's current implementation on a single workstation (64 GB RAM, NVMe storage) comfortably handles datasets across all non-biobank domains listed above. The two datasets used in this paper—rice 3K (3,024 accessions, 29.6M SNPs) and human 1000 Genomes (3,202 samples, 70.7M variants across 22 autosomes)—are representative of the upper range of typical population genomics projects.

#### Why $O(V \times K)$ matters most for iterative, multi-population analyses

The complexity advantage of GraphPop is most pronounced in workflows that involve repeated analyses across multiple populations, chromosomes, or annotation conditions—precisely the workflows that dominate agricultural and ecological population genomics. Consider a crop breeding programme analysing 12 subpopulations across 12 chromosomes with 6 FAST PATH statistics:

- **Matrix-based approach**: $12 \times 12 \times 6 = 864$ independent $O(V \times N)$ computations, each re-reading the full genotype matrix.
- **GraphPop**: 864 $O(V \times K)$ queries, each reading only the $K$-length allele-count arrays. Import is paid once.

With $N = 3{,}000$ and $K = 12$, GraphPop's per-query runtime is $\sim 250\times$ lower, and the total workflow completes in minutes rather than days. Adding annotation conditioning (e.g., missense vs synonymous) doubles the number of queries to 1,728 but adds negligible marginal cost to GraphPop, while doubling the total computation time for matrix-based tools.

##### Scaling path to larger datasets

The $O(V \times K)$ complexity reduction is a property of the pre-aggregation strategy, not of any specific graph database engine. GraphPop's stored procedures depend only on the labelled property graph model and Cypher query language, both of which are supported by multiple graph database engines with different scaling characteristics:

- **NebulaGraph**[20]: open-source, distributed graph database designed for billion-edge graphs with horizontal scaling across commodity hardware. Supports the property graph model and a Cypher-compatible query language (nGQL).
- **Neo4j AuraDB**[21]: fully managed cloud service with auto-scaling and enterprise-grade performance. Supports the same Cypher query language used by GraphPop.
- **Amazon Neptune** (with openCypher), **TigerGraph**, **FalkorDB**: additional engines supporting variants of the property graph model.

Migration to any of these engines would require only deployment configuration changes—not modifications to GraphPop's analytical logic. This engine-independence means that the $O(V \times K)$ complexity advantage demonstrated on medium-scale datasets in this paper could, in principle, be extended to biobank-scale datasets ($> 100{,}000$ individuals) through distributed graph database deployment, with the pre-aggregation step parallelised across partitioned chromosome segments.

#### Supplementary Tables

#### Supplementary Figures

#### References


[1] 1000 Genomes Project Consortium. A global reference for human genetic variation. *Nature* **526**, 68–74 (2015).

[2] Gillespie, M. *et al.* The Reactome pathway knowledgebase 2022. *Nucleic Acids Research* **50**, D588–D592 (2022).

**Table S2** Per-population diversity statistics for the rice 3K dataset (genome-wide means across 12 chromosomes).

| Rank | Population | Mean $\pi$ | Mean $\theta_W$ | Mean Tajima's $D$ | Mean $F_{IS}$ |
|---|---|---|---|---|---|
| 1 | admix | 0.07770 | 0.11278 | −1.117 | 0.616 |
| 2 | XI-adm | 0.06052 | 0.11325 | −1.493 | 0.632 |
| 3 | XI-2 | 0.05707 | 0.09338 | −1.528 | 0.632 |
| 4 | cA-Aus | 0.05488 | 0.12711 | −0.939 | 0.648 |
| 5 | XI-3 | 0.05418 | 0.08262 | −1.092 | 0.615 |
| 6 | XI-1B | 0.05064 | 0.09863 | −1.306 | 0.598 |
| 7 | XI-1A | 0.04691 | 0.08591 | −0.859 | 0.541 |
| 8 | cB-Bas | 0.04489 | 0.07592 | −2.043 | 0.663 |
| 9 | GJ-trp | 0.03239 | 0.05454 | −1.312 | 0.700 |
| 10 | GJ-adm | 0.03234 | 0.06210 | −0.988 | 0.706 |
| 11 | GJ-sbtrp | 0.02579 | 0.04686 | −0.890 | 0.673 |
| 12 | GJ-tmp | 0.01891 | 0.05244 | −2.406 | 0.720 |

**Table S3** $\pi_N/\pi_S$ ratios for all 12 rice subpopulations (genome-wide means). All values $> 1.0$ indicate relaxed purifying selection (cost of domestication).

| Rank | Population | Mean $\pi_N/\pi_S$ | Interpretation |
|---|---|---|---|
| 1 | admix | 1.146 | Highest deleterious load |
| 2 | GJ-adm | 1.145 | Admixed accumulation |
| 3 | GJ-trp | 1.107 | Tropical japonica |
| 4 | XI-adm | 1.079 | Indica admixed |
| 5 | cB-Bas | 1.078 | Basmati |
| 6 | GJ-sbtrp | 1.077 | Subtropical japonica |
| 7 | cA-Aus | 1.076 | Aus group |
| 8 | XI-2 | 1.061 | South Asian indica |
| 9 | XI-3 | 1.058 | SE Asian indica |
| 10 | XI-1B | 1.054 | Modern indica |
| 11 | XI-1A | 1.053 | East Asian indica |
| 12 | GJ-tmp | 1.018 | Most constrained |


[3] Wang, W. *et al.* Genomic variation in 3,010 diverse accessions of Asian cultivated rice. *Nature* **557**, 43–49 (2018).

[4] Naithani, S. *et al.* Plant Reactome: a knowledgebase and resource for comparative pathway analysis. *Nucleic Acids Research* **48**, D1093–D1103 (2020).

[5] Narasimhan, V. *et al.* BCFtools/RoH: a hidden Markov model approach for detecting autozygosity from next-generation sequencing data. *Bioinformatics* **32**, 1749–1751 (2016).

**Table S4** Annotation-conditioned Fst: missense vs. synonymous variant classes for three major population pairs.

| Population pair | Fst (missense) | Fst (synonymous) | Ratio | Interpretation |
|---|---|---|---|---|
| GJ-tmp vs GJ-trp | 0.252 | 0.228 | 1.106 | Adaptive protein divergence |
| GJ-tmp vs XI-1A | 0.604 | 0.579 | 1.044 | Adaptive protein divergence |
| XI-1A vs cA-Aus | 0.366 | 0.356 | 1.028 | Adaptive protein divergence |

**Table S5** Top 10 most differentiated rice population pairs by W&C Fst (genome-wide mean).

| Population pair | Mean Fst |
|---|---|
| GJ-tmp vs XI-1A | 0.710 |
| GJ-tmp vs XI-1B | 0.694 |
| GJ-tmp vs cA-Aus | 0.678 |
| GJ-tmp vs XI-2 | 0.663 |
| XI-3 vs GJ-tmp | 0.652 |
| XI-1A vs GJ-sbtrp | 0.636 |
| GJ-trp vs XI-1A | 0.622 |
| XI-1B vs GJ-sbtrp | 0.615 |
| GJ-trp vs XI-1B | 0.603 |
| XI-1A vs GJ-adm | 0.596 |

**Table S6** Known domestication genes recovered at selection scan peaks via graph traversal.

| Gene | Function | Statistic | Value | Comparison | Method |
|---|---|---|---|---|---|
| *GW5* | Grain width | XP-EHH | 4.91 | XI-1A vs cA-Aus | Graph traversal |
| *Hd1* | Heading date | XP-EHH | −4.15 | GJ-trp vs XI-1A | Graph traversal |
| *PROG1* | Prostrate growth | XP-EHH | 4.53 | GJ-tmp vs GJ-trp | Graph traversal |
| *OsC1* | Hull color | Garud's $H_{12}$ | 0.250 | GJ-tmp | Graph traversal |
| *Wx* | Amylose (waxy) | Garud's $H_{12}$ | 0.138 | GJ-adm | Graph traversal |
| *DRO1* | Deep rooting | Garud's $H_{12}$ | 0.139 | GJ-tmp | Graph traversal |


[6] Voight, B. F., Kudaravalli, S., Wen, X. & Pritchard, J. K. A map of recent positive selection in the human genome. *PLoS Biology* **4**, e72 (2006).

[7] Miles, A. *et al.* scikit-allel: a Python package for exploring and analysing genetic variation data. https://github.com/cggh/scikit-allel (2024). V1.3.7.

**Table S7** Consequence-level selection bias in rice: mean Fst by VEP impact class (GJ-tmp vs XI-1A).

| Impact class | $n$ variants | Mean Fst | Median Fst | Mann–Whitney $p$ vs LOW |
|---|---|---|---|---|
| HIGH | 18,284 | 0.185 | 0.018 | $< 10^{-300}$ |
| MODERATE | 26,318 | 0.129 | 0.005 | $7.5 \times 10^{-150}$ |
| LOW | 33,710 | 0.092 | 0.004 | — |

**Table S8** GraphPop procedure reference: 12 stored procedures with descriptions.

| Procedure | Path | Description |
|---|---|---|
| graphpop.diversity | FAST | $\pi$, $\theta_W$, Tajima's $D$, Fay & Wu's $H$, $H_e$, $H_o$, $F_{IS}$ |
| graphpop.divergence | FAST | Hudson Fst, W&C Fst, $D_{xy}$, $D_a$, PBS |
| graphpop.sfs | FAST | Site frequency spectrum (folded/unfolded) |
| graphpop.joint_sfs | FAST | 2D joint SFS |
| graphpop.genome_scan | FAST | Sliding-window scan (materializes GenomicWindow nodes) |
| graphpop.pop_summary | FAST | Whole-chromosome summary (writes to Population node) |
| graphpop.ld | FULL | Pairwise $r^2$, $D'$ (writes LD edges) |
| graphpop.ihs | FULL | Integrated haplotype score (AF-bin standardized) |
| graphpop.xpehh | FULL | Cross-population EHH (genome-wide standardized) |
| graphpop.nsl | FULL | Number of segregating sites by length |
| graphpop.roh | FULL | Runs of homozygosity (HMM or sliding-window) |
| graphpop.garud_h | FULL | Garud's $H_1$, $H_{12}$, $H_2/H_1$ (sweep detection) |


[8] Sabeti, P. C. *et al.* Genome-wide detection and characterization of positive selection in human populations. *Nature* **449**, 913–918 (2007).

[9] Ferrer-Admetlla, A., Liang, M., Korneliussen, T. & Nielsen, R. On detecting incomplete soft or hard selective sweeps using haplotype structure. *Molecular Biology and Evolution* **31**, 1275–1291 (2014).

[10] Purcell, S. *et al.* PLINK: a tool set for whole-genome association and population-based linkage analyses. *American Journal of Human Genetics* **81**, 559–575 (2007).

[11] Danecek, P. *et al.* The variant call format and VCFtools. *Bioinformatics* **27**, 2156–2158 (2011).

[12] Chang, C. C. *et al.* Second-generation PLINK: rising to the challenge of larger and richer datasets. *GigaScience* **4**, 7 (2015).

**Table S9** Novel analyses enabled by GraphPop's co-located individual genotype, annotation, and selection graph.

| Analysis | Classical approach | GraphPop |
|---|---|---|
| Pathway-level population Fst | VCFtools Fst → BEDtools gene overlap → pathway API → manual join | Single graph traversal through `IN_PATHWAY` |
| Convergent sweep detection | 5 XP-EHH runs → 5 output files → file intersection → coordinate join | Graph pattern match on stored `xpehh_*` node properties |
| Individual process-space embedding | PLINK ROH → custom het script → VEP → rare variant filter → manual join | Pre-stored FROH + single haplotype array pass |
| Outlier → driver variant trace | Extract single-sample VCF → VEP → pathway join | 3-hop graph traversal: Sample → CARRIES → HAS_CONSEQUENCE → IN_PATHWAY |
| Pathway-stratified purging gradient | Not feasible (requires genotype × consequence × pathway × FST) | Single session: haplotype arrays + `IN_PATHWAY` + prior Fst results |


[13] Bycroft, C. *et al.* The UK Biobank resource with deep phenotyping and genomic data. *Nature* **562**, 203–209 (2018).

[14] Karczewski, K. J. *et al.* The mutational constraint spectrum quantified from variation in 141,456 humans. *Nature* **581**, 434–443 (2020).

[15] Hayes, B. J. & Daetwyler, H. D. 1000 Bull Genomes Project to map simple and complex genetic traits in cattle: applications and outcomes. *Annual Review of Animal Biosciences* **7**, 89–102 (2019).

[16] Kijas, J. W. *et al.* Genome-wide analysis of the world's sheep breeds reveals high levels of historic mixture and strong recent selection. *PLoS Biology* **10**, e1001258 (2012).

[17] Neale, D. B. & Kremer, A. Forest tree genomics: growing resources and applications. *Nature Reviews Genetics* **12**, 111–122 (2011).

[18] Lamichhaney, S. *et al.* Evolution of Darwin's finches and their beaks revealed by genome sequencing. *Nature* **518**, 371–375 (2015).

[19] Zhao, K. *et al.* An Arabidopsis example of association mapping in structured populations. *PLoS Genetics* **3**, e4 (2007).

[20] Vesoft Inc. NebulaGraph: an open-source distributed graph database. https://nebula-graph.io/ (2025). Accessed 2026-04-08.

[21] Neo4j, Inc. Neo4j AuraDB: fully managed cloud graph database. https://neo4j.com/cloud/auradb/ (2025). Accessed 2026-04-08.

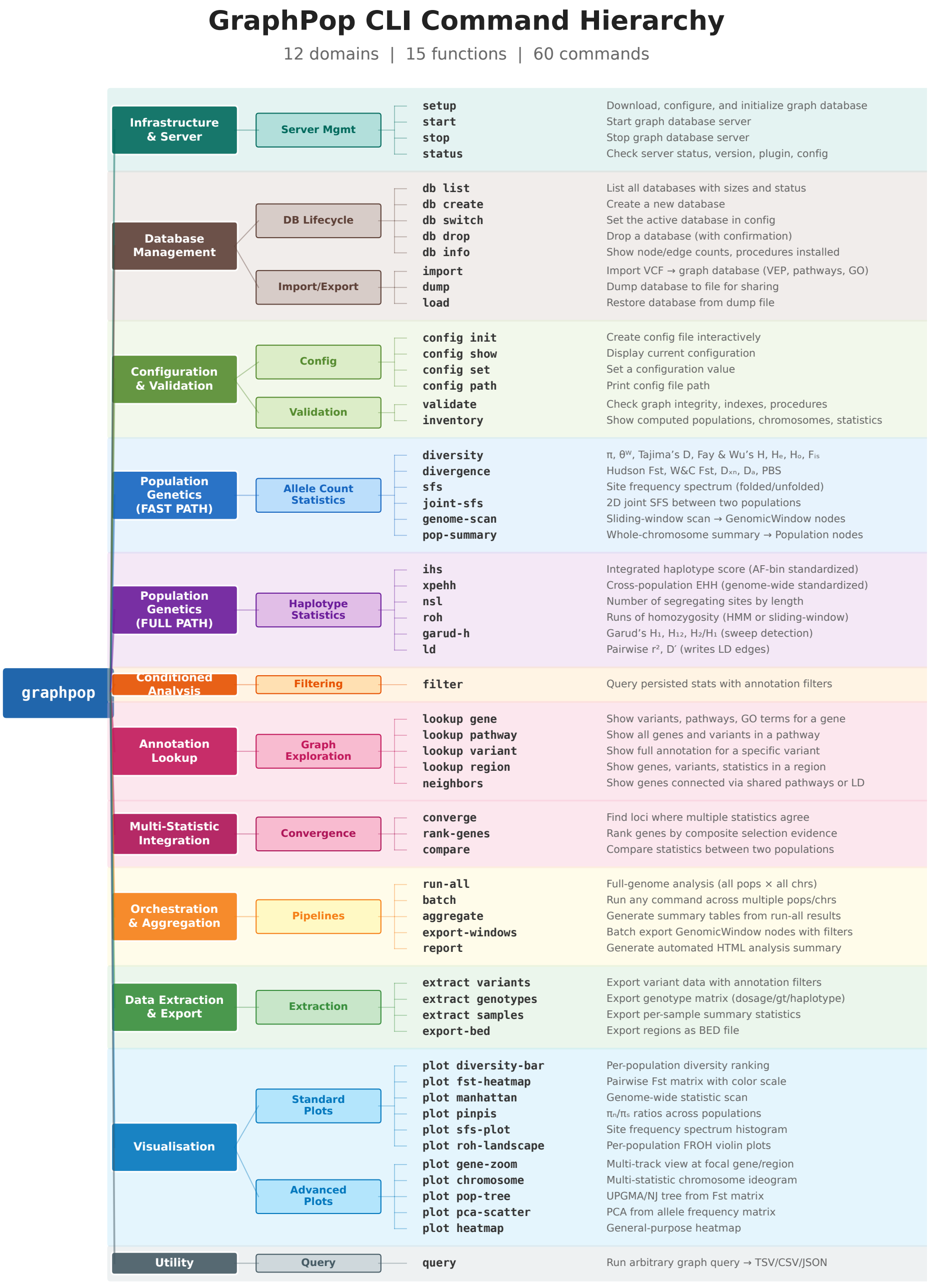


**Fig. S1 GraphPop CLI command hierarchy.** The 62 commands are organised into 11 functional domains spanning the complete analytical lifecycle: infrastructure and server management, database lifecycle, configuration and validation, 12 population genetics procedures (6 FAST PATH on pre-aggregated allele counts, 6 FULL PATH on bit-packed haplotypes), annotation-conditioned filtering, annotation lookup and graph exploration, multi-statistic integration, orchestration and aggregation, data extraction and export, and 11 publication-ready visualisation types. Users require no knowledge of graph databases, graph query languages, or Python—the graph architecture is fully abstracted behind a conventional command-line interface.

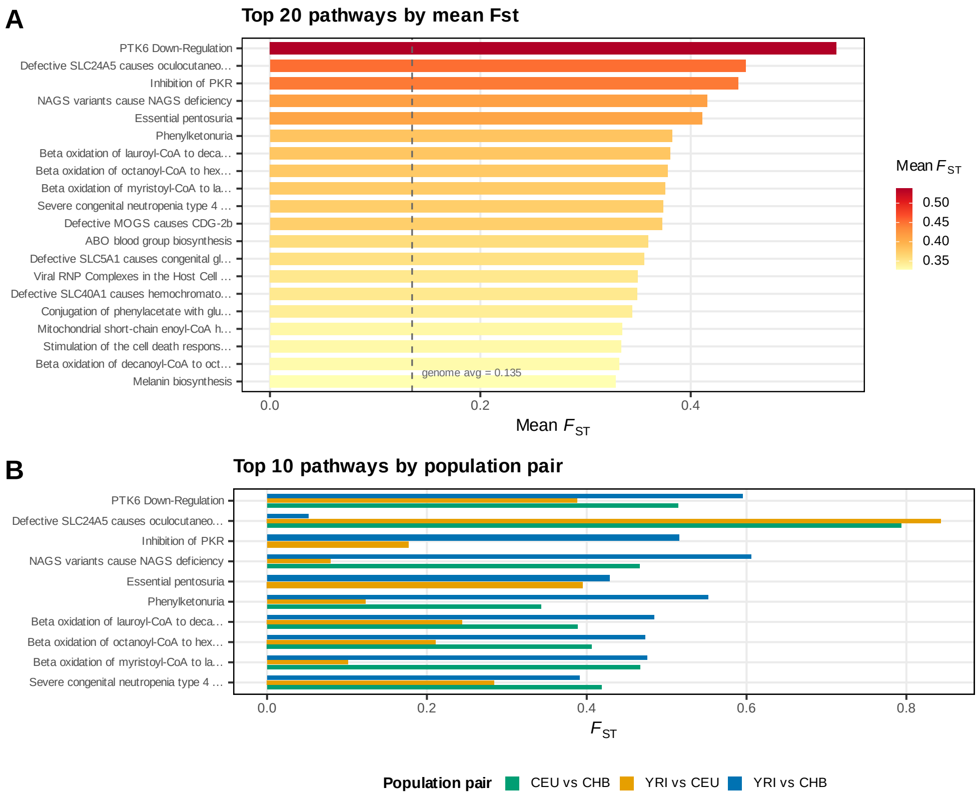


**Fig. S2 Pathway-level population differentiation (YRI vs CEU).** Mean Hudson Fst per Reactome pathway, ranked by magnitude. The *SLC24A5*-linked oculocutaneous albinism pathway shows the highest differentiation (mean Fst = 0.452). Cardiac ion channel pathways (TREK/TWIK, Phase 3 repolarisation) form a functionally coherent cluster in the top quartile.

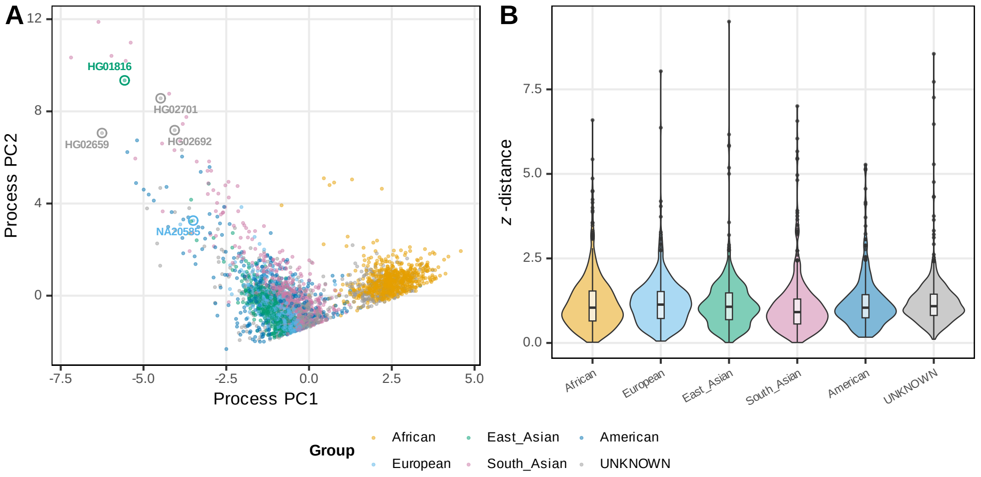


**Fig. S3 Outlier individual analysis.** Z-distance from population centroid for all 3,202 individuals. HG01816 (CDX, Chinese Dai) is the most extreme outlier ($z = 9.50$), with genome-wide FROH = 0.125—18× the CDX population mean—consistent with recent close-relative mating.

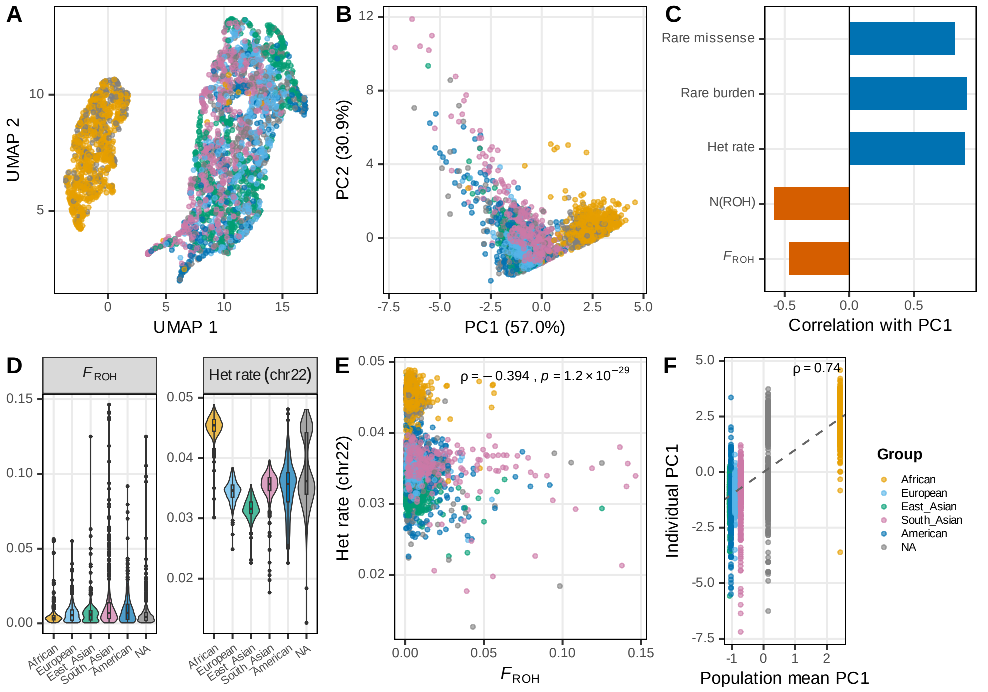


**Fig. S4 Individual-level evolutionary trajectory in process space.** Each of 3,202 individuals is characterised by five evolutionary process statistics (genome-wide FROH, ROH count, chr22 heterozygosity, rare burden, rare missense burden) and embedded via PCA and UMAP. PC1 (57.0%) separates populations along a diversity axis (African highest); PC2 (30.9%) separates along an inbreeding axis (South Asian highest). UMAP reveals clear continental clustering in process space. Individual positions strongly predict their population's position ($\rho = 0.74$).

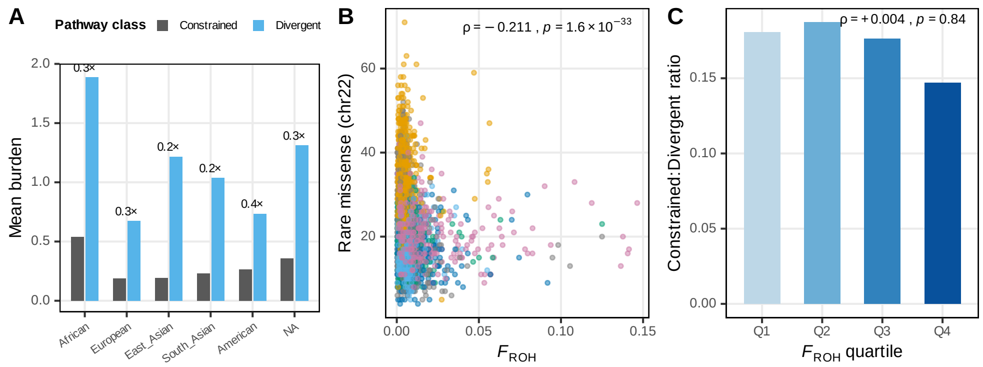


**Fig. S5 Pathway-stratified purifying selection gradient.** Rare missense burden per individual, stratified by constrained (low-Fst) vs. divergent (high-Fst) pathways. The 2× African enrichment holds at equal magnitude in both pathway classes, confirming that bottleneck-driven depletion of rare functional variants is genome-wide and uniform. FROH predicts individual burden ($\rho = -0.211$) but not the constrained:divergent ratio ($\rho = +0.004$).

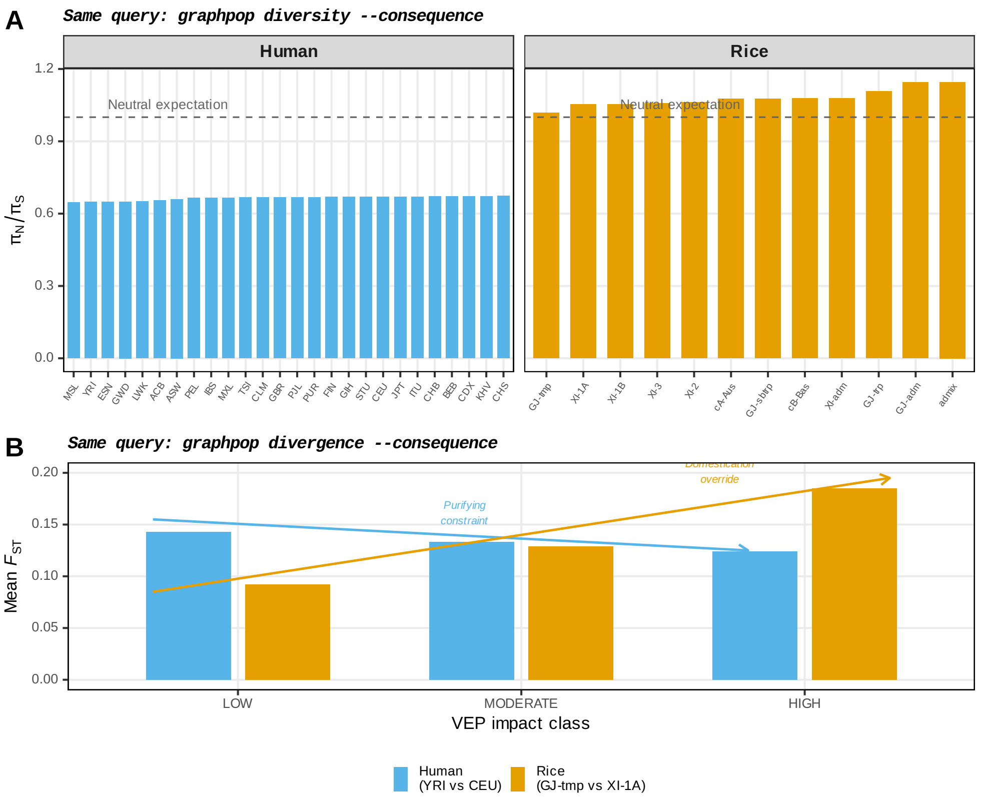


**Fig. S6 Same annotation-conditioned query applied to two species. a**, $\pi_N/\pi_S$ computed with the identical `graphpop diversity --consequence` command for all human (26) and rice (12) populations. All rice subpopulations exceed the neutral expectation ($\pi_N/\pi_S > 1.0$), indicating universal relaxation of purifying selection under domestication; all human populations fall below 1.0, confirming efficient purifying selection under natural evolution. **b**, Mean Fst by VEP impact class for one representative population pair per species, computed with the same `graphpop divergence --consequence` command. Humans show decreasing Fst with increasing impact severity (purifying constraint), while rice shows increasing Fst (domestication override). The cross-species reversal emerges from the same single-line query applied to two datasets stored in the same graph schema.

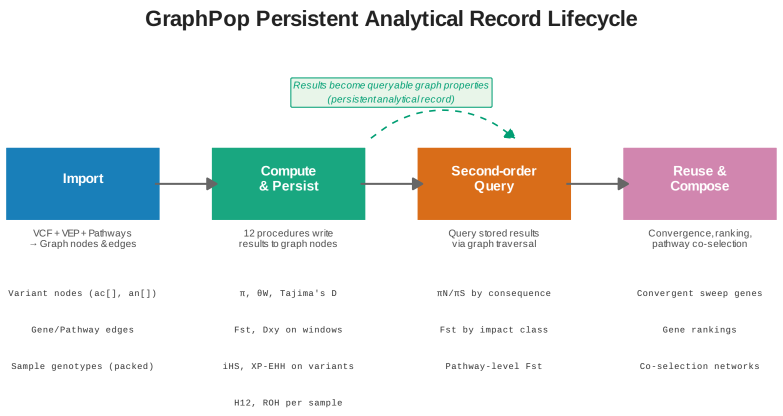


**Fig. S7 Persistent analytical record lifecycle.** GraphPop's four-stage analytical lifecycle. (1) Import: VCF genotypes, VEP annotations, and pathway data are loaded as graph nodes and edges. (2) Compute & Persist: 12 stored procedures write results (diversity, Fst, iHS, etc.) as properties on the same nodes that carry the biological data. (3) Second-order Query: stored results are queried by graph traversal for annotation-conditioned analyses (e.g., $\pi_N/\pi_S$ by consequence class, Fst by impact level). (4) Reuse & Compose: results computed for one purpose are reused for qualitatively different analyses (convergent sweep detection, gene ranking, pathway co-selection networks) without re-computation. The persistent analytical record—computed statistics stored as graph node properties—is the key architectural property that enables stages 3 and 4.

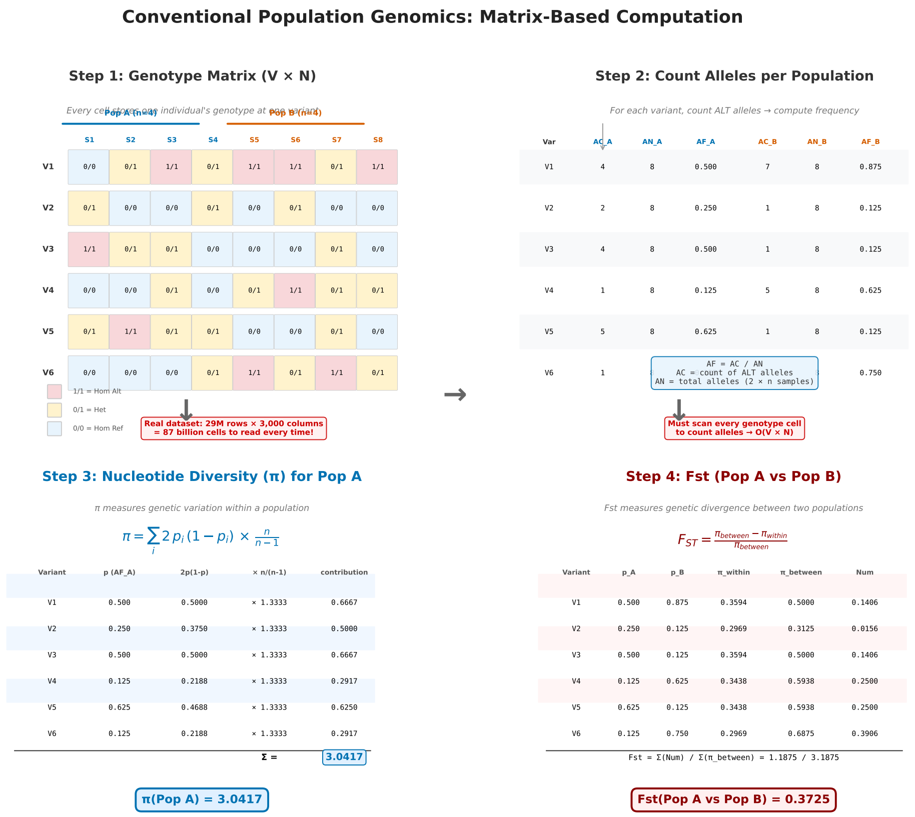


**Fig. S8 Conventional matrix-based computation of $\pi$ and Fst—worked example.** Step-by-step illustration of the classical population genomics pipeline applied to a small example dataset (6 variants × 8 samples, split into Pop A and Pop B). **Step 1**: the genotype matrix stores one genotype per cell (`0/0`, `0/1`, `1/1`); the matrix size scales as $V \times N$ and reaches 87 billion cells for the rice 3K dataset. **Step 2**: every cell is scanned to count alternative alleles per population, yielding allele counts (`AC`), total alleles (`AN`), and frequencies (`AF`). **Step 3**: nucleotide diversity $\pi$ for Pop A is computed by summing $2p_i(1-p_i) \times n/(n-1)$ over variants. **Step 4**: Fst between Pop A and Pop B is computed from $\pi_{within}$ and $\pi_{between}$. Every step requires reading the entire genotype matrix—O$(V \times N)$ complexity—making repeated queries across populations and chromosomes expensive.

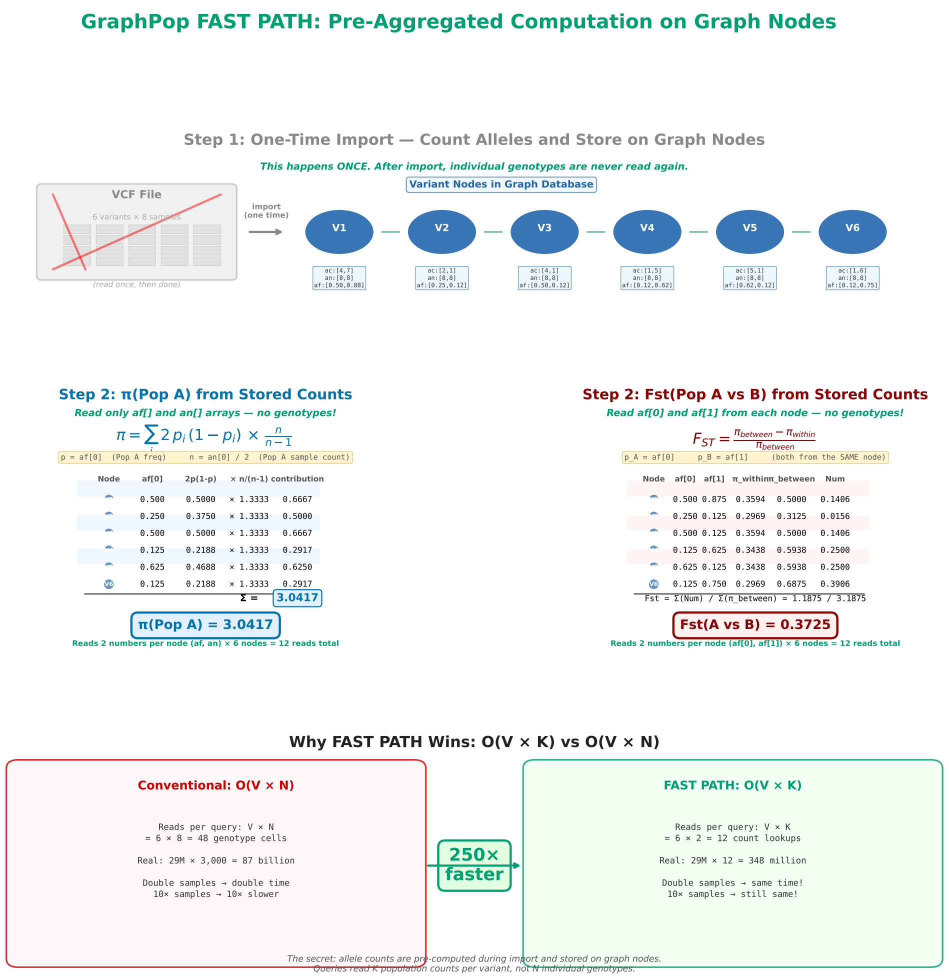


**Fig. S9 GraphPop FAST PATH computation of $\pi$ and Fst—same example, different complexity.** The same six-variant example dataset as in Supplementary Fig. S8, computed via GraphPop's FAST PATH. **Step 1** (one-time import): allele counts per population are pre-aggregated and stored as node properties (`ac[]`, `an[]`, `af[]`) on each Variant node; individual genotypes are never read again after import. **Step 2**: $\pi$(Pop A) is computed by reading only the `af[0]` and `an[0]` arrays from each Variant node—12 numerical reads instead of 48 genotype-cell scans. The result is identical to the conventional method ($\pi = 3.0417$). **Step 3**: Fst(Pop A vs Pop B) requires only `af[0]` and `af[1]` per node—another 12 reads. The bottom panel contrasts the two scaling regimes: $O(V \times N)$ reads grow with sample count, while $O(V \times K)$ reads stay fixed at the population dimension. For the rice 3K dataset, this is a $\sim$250$\times$ reduction in operations per query.

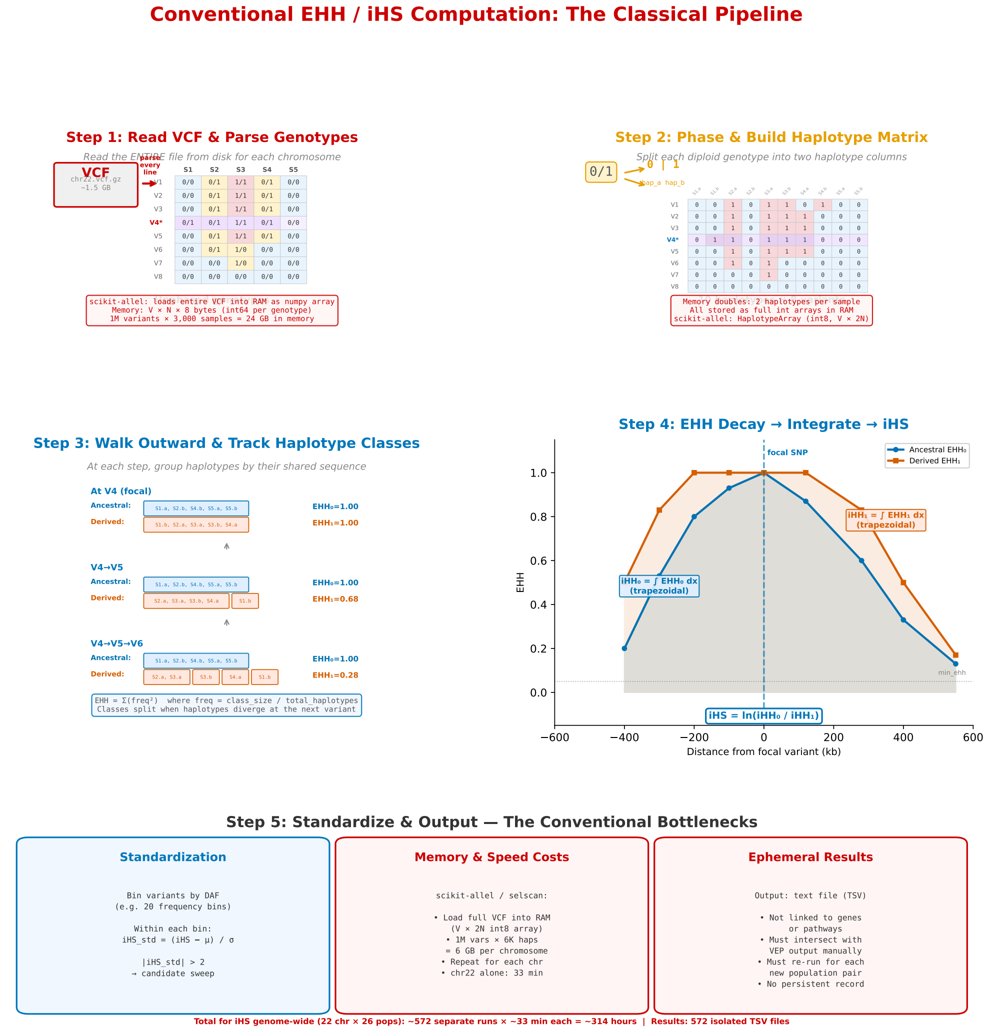


**Fig. S10 Conventional EHH and iHS computation—worked example.** The classical pipeline for haplotype-based selection scans. **Step 1**: parse the entire VCF and load genotypes into RAM as a $V \times N$ int array (24 GB for 1M variants $\times$ 3,000 samples in scikit-allel). **Step 2**: phase each diploid genotype into two haplotype columns, doubling the memory footprint (HaplotypeArray of $V \times 2N$). **Step 3**: starting from a focal variant, walk outward in both directions; at each step, group haplotypes by their shared sequence and recompute the EHH statistic (sum of squared class frequencies). **Step 4**: integrate EHH curves for ancestral and derived alleles to obtain $\mathrm{iHH}_0$ and $\mathrm{iHH}_1$, then compute iHS as the log ratio. **Step 5**: standardize iHS within derived allele frequency bins. The classical approach loads the entire VCF into memory, processes one chromosome at a time, and writes results to isolated TSV files with no link back to gene or pathway annotations. Genome-wide iHS for 22 autosomes $\times$ 26 populations requires $\sim$572 separate runs and $\sim$314 hours of compute.

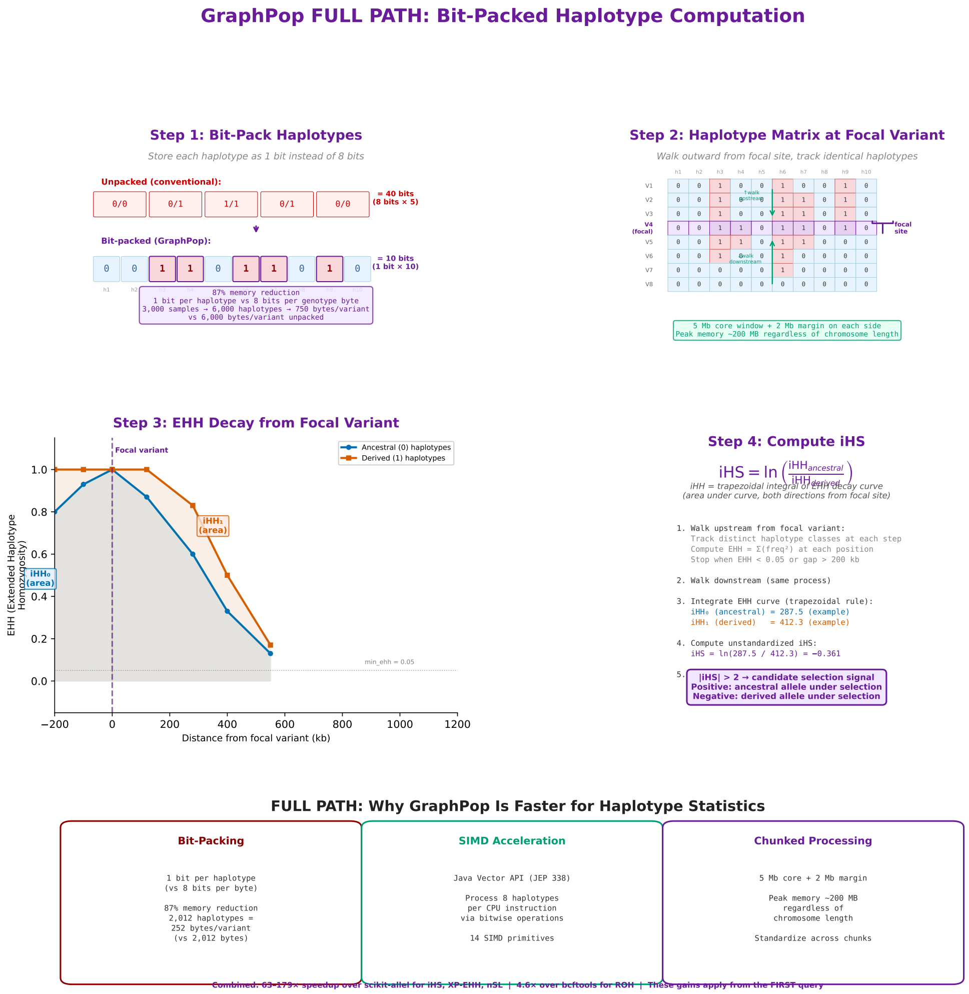


**Fig. S11 GraphPop FULL PATH computation of EHH and iHS—bit-packed haplotype computation.** GraphPop's FULL PATH for haplotype-based statistics. **Step 1**: bit-pack each haplotype as a single bit instead of 8 bits per byte, achieving 87% memory reduction (252 bytes per variant for 2,012 haplotypes vs 2,012 bytes unpacked). **Step 2**: load the haplotype matrix from the graph database into a dense bit-packed structure; walk outward from the focal variant. **Step 3**: compute EHH at each step using SIMD-accelerated bitwise operations (Java Vector API, JEP 338). **Step 4**: integrate EHH curves and compute iHS, then standardize within derived allele frequency bins. The bottom strip summarises the three pillars of FULL PATH performance: bit-packing (87% memory reduction), SIMD acceleration (8 haplotypes per CPU instruction), and chunked processing (5 Mb core windows with 2 Mb EHH margins, peak memory ∼200 MB regardless of chromosome length). Combined, these yield 63–179× speedups over scikit-allel for iHS, XP-EHH, and nSL—advantages that apply from the first query, independent of pre-aggregation.

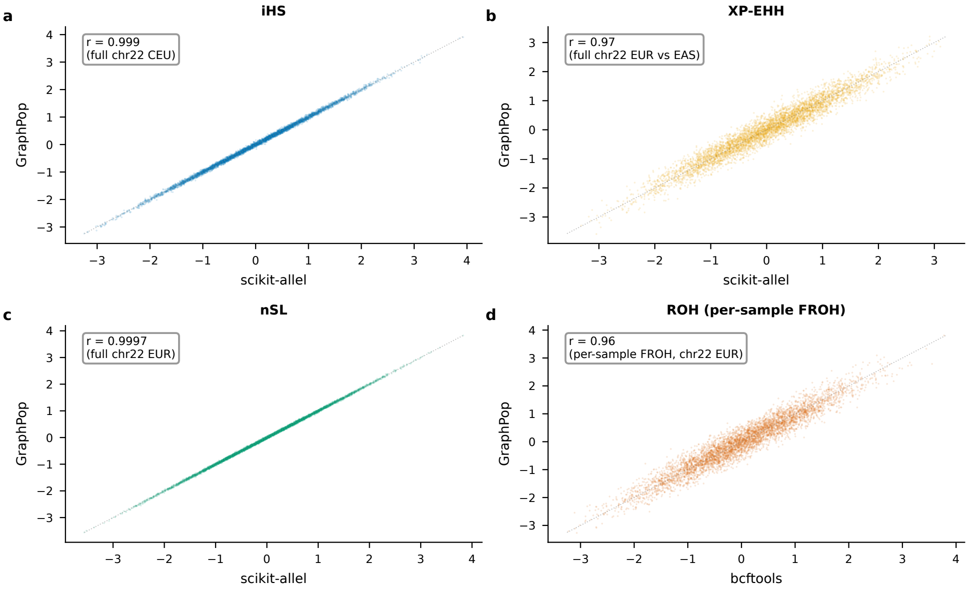


**Fig. S12 Per-locus validation of FULL PATH statistics.** Scatterplots comparing GraphPop FULL PATH output against competitor tools on 1000 Genomes chr22. **a**, iHS (GraphPop vs scikit-allel, CEU population, $r = 0.999$). **b**, XP-EHH (GraphPop vs scikit-allel, EUR vs EAS, $r = 0.97$). **c**, nSL (GraphPop vs scikit-allel, EUR, $r = 0.9997$). **d**, ROH (per-sample FROH, GraphPop vs bcftools, EUR, $r = 0.96$). The near-unity correlations for iHS and nSL confirm numerical equivalence. The lower XP-EHH correlation ($r = 0.97$) reflects differences in variant filtering (GraphPop requires MAF $\geq 0.05$ in at least one population) and EHH truncation thresholds. The ROH correlation ($r = 0.96$) reflects differences in HMM transition rate parameterization between GraphPop and bcftools. In all four panels, the identity line (grey dotted) and regression line (black dashed) are shown.
